# Comprehensive locus-specific L1 DNA methylation profiling reveals the epigenetic and transcriptional interplay between L1s and their integration sites

**DOI:** 10.1101/2023.01.03.522582

**Authors:** Sophie Lanciano, Claude Philippe, Arpita Sarkar, David Pratella, Cécilia Domrane, Aurélien J. Doucet, Dominic van Essen, Simona Saccani, Laure Ferry, Pierre-Antoine Defossez, Gael Cristofari

**Affiliations:** University Cote d’Azur, Inserm, CNRS, Institute for Research on Cancer and Aging of Nice (IRCAN), Nice, France; University Paris Cité, CNRS, Epigenetics and Cell Fate, Paris, France

**Author notes:** These authors contributed equally (Sophie Lanciano, Claude Philippe).

**Keywords:** LINE-1, L1, transposable elements, mobile genetic element, transposon, transposition, retrotransposon, retrotransposition, mobile element insertion, DNA methylation, methyl-cytosine, nanopore sequencing, chromatin states, transcription, L1 chimeric transcripts, LCTs, YY1, ESR1, transcription factor, nuclear receptor, genomic profiling

## Abstract

Long interspersed element-1 (L1) retrotransposons play important roles in human disease and evolution. Their global activity is repressed by DNA methylation, but studying the regulation of individual copies has been difficult. Here, we combine short- and long-read sequencing to resolve the DNA methylation profiles of these repeated sequences in a panel of normal and cancer cells genome-wide at single-locus resolution. We unveil key principles underpinning L1 methylation heterogeneity among cell-types, families and integration sites. First, intronic L1 methylation is intimately associated with gene transcription. Conversely, L1s can influence the methylation status of the upstream region over short distances (300 bp). This phenomenon is accompanied by the binding of specific transcription factors, which drive the expression of L1 and chimeric transcripts. Finally, L1 hypomethylation alone is generally insufficient to trigger L1 expression due to redundant silencing pathways. Our results illuminate the epigenetic and transcriptional interplay between retrotransposons and their host genome.

**GRAPHICAL ABSTRACT:** 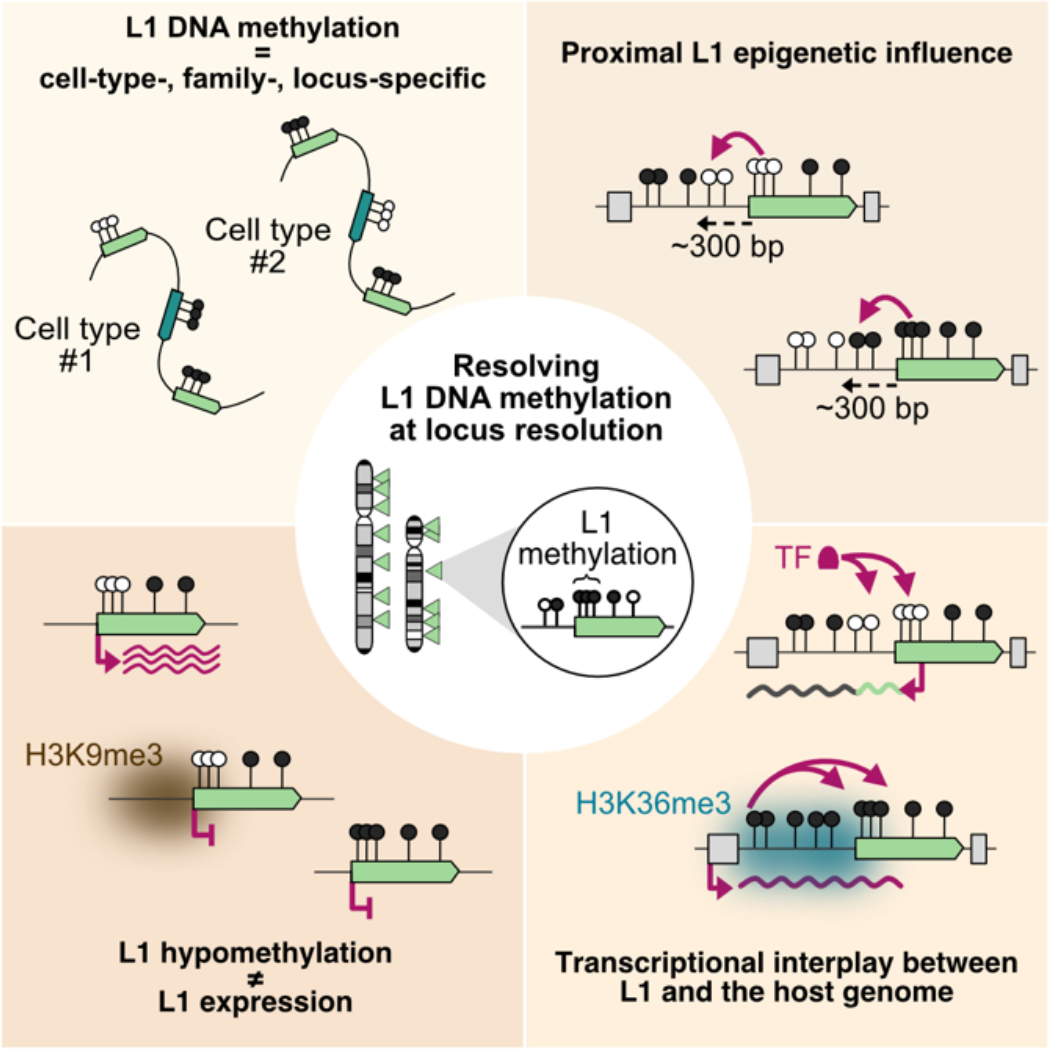

**HIGHLIGHTS:**

- Bs-ATLAS-seq profiles L1 position and methylation genome-wide
- L1 has a frequent but short-range (300 bp) influence on the DNA methylation status of the upstream sequence
- Hypomethylated L1s are bound by tissue-specific transcription factors which drive L1 and chimeric transcripts synthesis
- L1 hypomethylation alone is insufficient to enable its transcription at most loci

## INTRODUCTION

Transposable elements represent a considerable fraction of mammalian genomes and contribute substantially to their gene regulatory networks ^1, 2^. In humans, the long interspersed element-1 (LINE-1 or L1) retrotransposon accounts for at least 17% of the genome, and is the sole autonomously active transposable element ^3^. Its dramatic expansion to approximately half a million copies is driven by a copy-and-paste mechanism named retrotransposition. This process is mediated by two L1-encoded proteins, ORF1p and ORF2p, which associate with the L1 mRNA to form a ribonucleoprotein particle (RNP), considered as the core of the retrotransposition machinery. The L1 RNP cleaves the host DNA and directly synthesizes a new L1 copy at integration sites through target-primed reverse transcription ^4–7^. Retrotransposition thus depends on L1 transcription, which initiates at an internal CpG-rich bidirectional promoter located in its 5’-untranslated region (UTR) (**Figure 1A**). The sense promoter (SP) drives the synthesis of the L1 mRNA, with a significant fraction of read-through into the downstream genomic sequence ^8–12^. As for the antisense promoter (ASP), it can act as an alternative promoter for neighboring genes, leading to spliced chimeric transcripts, or even to fusion proteins with an L1-encoded antisense ORF, called ORF0 ^13–20^.

**Figure 1.**
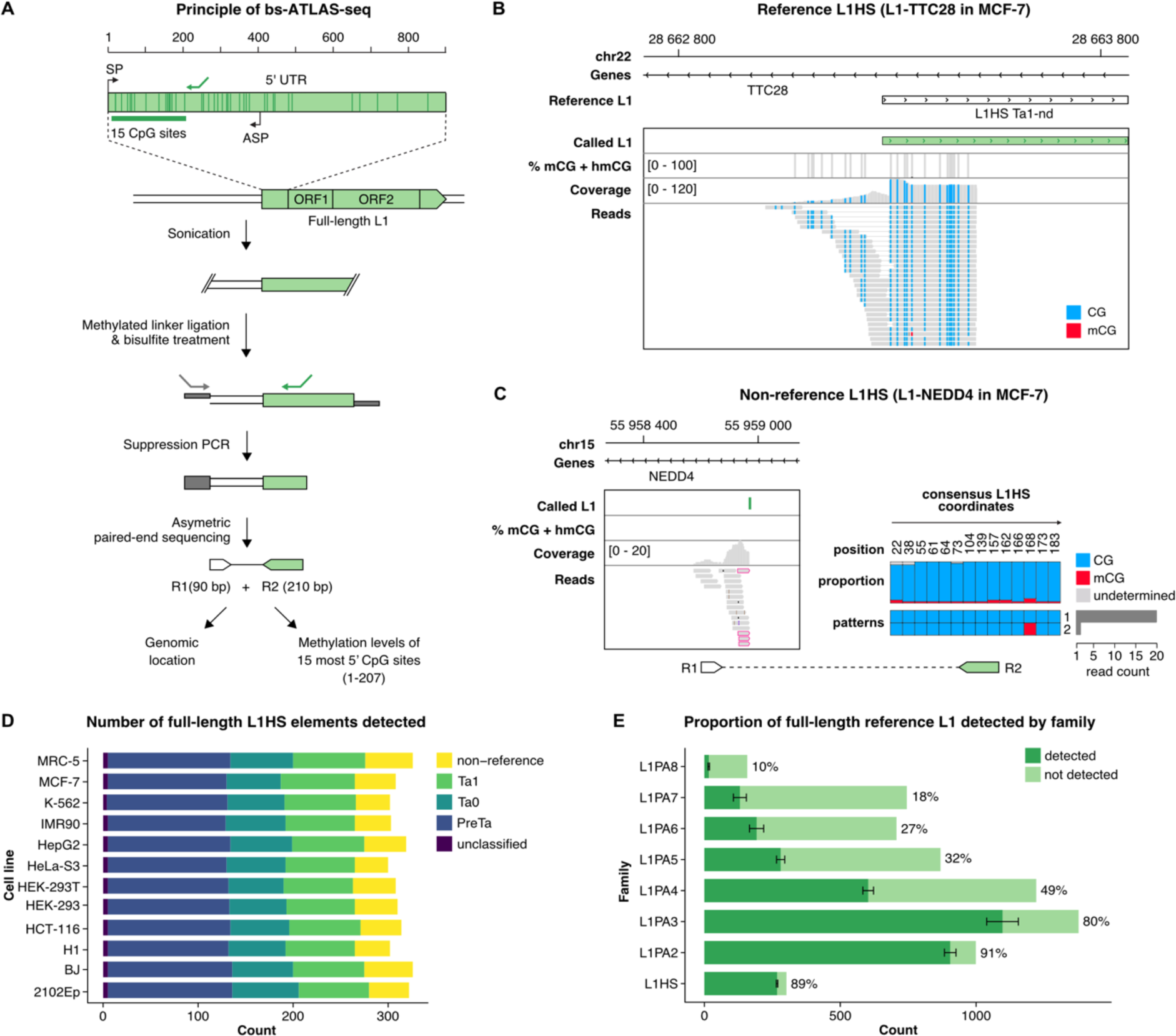
Bisulfite-ATLAS-seq (bs-ATLAS-seq) profiling of human L1 element promoters. **(A)** Principle of the bs-ATLAS-seq method. The internal L1 promoter region (∼ 900 bp) is illustrated (top). Transcription start sites for the sense (SP) and antisense (ASP) promoters are represented as broken arrows and overlap with the L1 5’UTR. Bs-ATLAS-seq interrogates the first 15 CpG sites of the L1 promoter, shown as vertical bars in the magnified view (bottom). The L1-specific primer used to amplify L1 junctions is shown as a green arrow. Genomic DNA is fragmented by sonication and ligated to a single-stranded methylated linker. Linker-ligated DNA is then treated with bisulfite and L1-containing fragments are specifically amplified by suppression PCR. In this approach, the linker is single-stranded and possesses the same sequence as the linker-specific primer (not its complementary sequence, grey arrow). Consequently, amplification only occurs upon prior extension from the L1-specific primer (green arrow) and synthesis of the linker complementary sequence (not shown). This strategy prevents linker-to-linker amplification. The L1-specific primer was designed to enrich for the L1HS family, but older related L1PA elements are also amplified (see **Figure 1E**). Finally, asymmetric paired-end sequencing provides the genomic location as well as the methylation levels of each L1 locus. R1 and R2 refers to reads #1 and #2, respectively. Note that 5-methylcytosine (5mC) and 5-hydroxymethylcytosine (5hmC) are both protected from bisulfite-induced deamination, thus bs-ATLAS-seq cannot discriminate between these two DNA modifications. **(B, C)** Genome browser view of bs-ATLAS-seq results in the breast cancer cell line MCF-7 for a reference (B) and a non-reference (C) L1HS element in the TTC28 and NEDD4 genes, respectively. In the track showing the percentage of methylation, called CpG are indicated by a vertical gray bar, and the percentage of methylation as an overlapping black bar. In the ‘coverage’ and ‘reads’ tracks, vertical colored bars correspond to non-methylated CpG (blue) and methylated CpG (mCG, red). Since bisulfite-sequencing-based methods cannot discriminate between hydroxymethylated CpG (hmCG) and methylated-CpG (mCG), methylation status is indicated as mCG + hmCG. (C) For non-reference L1HS, only the genomic flank covered by read #1 (R1; bottom left) is visible in the genome browser view. Soft-clipped reads supporting the 5’ L1 junction (split reads) are framed in pink. The proportion of mCG at each site and the frequency of the most common methylation patterns deduced from read 2 (R2; bottom right) are indicated on the charts (right). The positions of the CpGs are given relative to the L1HS consensus sequence (see Methods). **(D)** Number of full-length L1HS elements detected in the different cell lines by bs-ATLAS-seq, and their subfamilies. Pre-Ta, Ta0 and Ta1 represent different lineages of the L1HS family, from the oldest to the youngest, and were deduced from diagnostic nucleotides in L1 internal sequence (Boissinot et al., 2000) and thus could only be obtained for reference insertions as bs-ATLAS-seq provides only limited information on L1 internal sequence. **(E)** Fraction and count of full-length reference L1 elements detected by bs-ATLAS-seq for each L1 family. Bars represent the average number of full length reference L1 elements detected by bs-ATLAS-seq (dark green, mean ± s.d., n=12 cell lines), as compared to the total number of these elements in the reference genome (light green). Full length elements were defined as elements longer than 5,900 bp as annotated in UCSC repeatmasker track. The ratio of detected/total elements is indicated as a percentage on the right of each bar. Note that any given sample only contains a subset of reference L1HS due to insertional polymorphisms in the human population. See also **Figure S1** and **Table S1**.

The propagation of L1 elements throughout mammalian evolution occurred in successive waves of expansion and extinction implicating a limited number of concurrent families. In anthropoid primates, a single lineage, L1PA, has been active, leading to the sequential emergence of the L1PA8 to L1PA1 families (from the oldest to the youngest) in the last 40 million years ^21^. L1PA1 - which is human-specific and also known as L1HS - is the only autonomous transposable element remaining active in modern humans ^22, 23^, with an estimated rate of one new insertion every 60 births ^24^. Such additional insertions not catalogued in the reference genome are referred to as non-reference elements and contribute to our genetic diversity ^23, 25–29^. As a result, two individual human genomes differ on average at ∼300 sites with respect to L1 presence or absence ^30^. Although all L1PA sequences are related, the 5’ UTR and ORF1 regions are subject to rapid adaptive evolution, likely resulting from an arms-race with the host genome ^31–33^. Moreover, following their integration, L1 sequences accumulate alterations such as mutations, indels, or nested transposable element insertions, and therefore diverge relative to their original progenitor ^34^. Thus, although L1 elements are highly repeated sequences, they also exhibit a significant level of heterogeneity, between and within families, particularly in their promoter region, suggesting that they may be subject to distinct regulatory mechanisms. In support of this possibility, a variety of Krüppel-associated box domain zinc finger proteins (KZFPs) can specifically bind L1PA8 to L1PA3 elements in different cell types, leading to TRIM28-dependent silencing ^33, 35, 36^, whereas younger L1PA2 and L1HS elements are presumably silenced through TRIM28-independent mechanisms, such as DNA methylation ^35, 37–42^. Similarly, the HUSH complex represses a subset of the youngest L1 families in various cell types ^43–45^.

Despite these repressive mechanisms, L1HS can mobilize in the germline and during early embryonic development, creating heritable genetic variations and potentially causing genetic diseases ^46^. In addition, L1HS can also retrotranspose in a few somatic tissues such as the brain ^46^, as well as in many epithelial tumors ^47, 48^. Of note, even if L1 elements older than L1HS have lost the ability to achieve a full cycle of retrotransposition, they can be transcribed, and their transcriptional reactivation is not without consequence. By providing alternative promoters and forming chimeric transcripts, they can be responsible for oncogene activation in some tumors ^49–51^ and lead to the formation of long non-coding RNA (lncRNA) in many tissues ^52^. Nuclear L1 transcripts and transcriptional activity have also been implicated in regulating nuclear architecture, chromatin accessibility and totipotency in mammals ^53–55^.

A prevailing idea is that the global loss of DNA methylation commonly seen in tumor cells is responsible for the frequent and generalized reactivation of L1 elements in cancer ^56^. Consistently, targeted analyses of selected progenitor L1 elements confirmed their hypomethylation in tumors compared to normal tissues ^12, 22, 48, 57, 58^. A similar association between L1 hypomethylation and activity was found in human pluripotent and neuronal progenitor cells ^39, 59–64^. However, early studies also suggested that DNA methylation can be heterogeneous between different L1 loci, particularly in tumor cells ^37, 42^. Such heterogeneity was also reported at the level of L1 expression, with only a restricted subset of L1 loci, and among them a handful of L1HS, being robustly expressed in transformed cells ^9, 22, 65^ and ultimately acting as source elements for new retrotransposition events in tumors ^12, 22, 28, 48^. Altogether, these observations suggest that DNA methylation must be lifted to permit L1 reactivation. Yet the extent of L1 DNA hypomethylation in tumor cells, and whether DNA demethylation is sufficient to promote L1 reactivation remains unclear. As L1s can efficiently insert into regions with a wide range of chromatin states ^66, 67^, it is conceivable that the locus of integration could influence the capacity of a given L1 element to be reactivated upon demethylation. Addressing these questions in a systematic way has been challenging so far, given the technical difficulties in assessing L1 DNA methylation genome-wide at single locus resolution, particularly for the most recent L1HS family ^68^.

Here, we employed a novel genome-wide approach, termed bs-ATLAS-seq, as well as targeted nanopore sequencing, to map the position of individual full-length L1 elements and reveal their methylation levels in a panel of normal, embryonal and tumoral human cell lines. This strategy uncovered the heterogeneity of L1 DNA methylation, which was previously masked by aggregate analyses, and further revealed that L1 methylation patterns are in fact highly cell-type-, family- and locus-specific. Yet, in most cell types, including cancer cells, the majority of L1HS remains hypermethylated. We observed that gene body methylation of transcriptionally active genes is associated with the methylation of intronic L1 elements in many cell types. Inversely, L1s can frequently affect DNA methylation in the neighboring genomic region within 300 bp. This epivariation is associated with unique transcription factors binding profiles controlling L1 expression and the synthesis of L1 chimeric transcripts with neighboring genes. Finally, L1 hypomethylation or pharmacological demethylation is not sufficient alone to trigger expression at most loci. Together, our results highlight the interplay between L1 retrotransposons and their integration sites with respect to DNA methylation and expression, and reveal the existence of redundant layers of epigenetic regulation at individual loci.

## RESULTS

### Genome-wide and locus-specific human L1 DNA methylation profiling

The L1 bidirectional promoter is 910 bp long and possesses in its first half a CpG island which coincides with its sense promoter activity (**Figure 1A**) ^69–71^. To individually measure the DNA methylation levels of each full-length L1HS copy inserted in the genome, we adapted to bisulfite-treated DNA the ATLAS-seq approach, a method originally developed to map L1HS elements in the human genome ^9, 72^. This strategy, termed bisulfite-ATLAS sequencing (bs-ATLAS-seq, **Figure 1A**), provides both the location of L1HS insertions and the methylation state of the most distal region of their promoter at single-locus, single-molecule and single-nucleotide resolutions (**Figure 1B** and **Figure S1A**). Importantly, these data are obtained both for reference and non-reference insertions (**Figure 1C** and **Figure S1B**). The amplified region within the L1 5’UTR covers the first 15 CpG dinucleotides - including 7 being considered as critical for L1 regulation ^73^ - and its methylation level is representative of the broader internal promoter (see below and **Figure S1I**). Therefore, for the sake of simplicity, and unless otherwise specified, we will hereafter use the term “L1 methylation” to refer to the DNA methylation of this distal region of the L1 promoter. Bs-ATLAS-seq only requires 10 million read pairs to recover 97% of detectable L1HS elements and 20 million read pairs to recover 100% of them (**Figure S1C**), and it is highly reproducible (**Figure S1D** and **Figure S1E**).

We applied bs-ATLAS-seq to a panel of 12 human cell lines, which includes normal primary fibroblasts (BJ, IMR90 and MRC-5), artificially immortalized and transformed cells (HEK-293, adenovirus-immortalized embryonal kidney cells; HEK-293T, a derivative of HEK-293 further transformed by SV40 Large T antigen), cancer cell lines (HepG2, hepatoblastoma; K-562, chronic myeloid leukemia; MCF-7, breast cancer; HeLa-S3, cervical cancer; HCT-116, colon cancer), as well as cells of embryonal origins (H1, embryonic stem cells; 2102Ep, embryonal carcinoma cells) (**Figure S1F, Table S1**). We compiled the results into a comprehensive database containing the position of each L1 copy in the 12 cell lines and their DNA methylation levels at single locus-, nucleotide-, and molecule-resolutions (**Table S3** and **Data S1**). These data can be easily and interactively interrogated through a dedicated web portal (https://L1methdb.ircan.org). On average, we identified 312 full-length L1HS in each cell line, of which 42 are non-reference insertions and assumed to be full-length (**Figure 1D**). In addition to these copies, we also detected an average of 81 L1HS reference elements with an amplifiable 5’ end but annotated as 3’ truncated (**Table S3**). Among them, half contain Alu insertions (AluY or AluS) or other forms of internal rearrangements, or represent chimeras with older L1PA elements. We identified approximately 90% of all full-length reference L1HS copies (**Figure 1E**), the undetected elements being likely absent from the assayed samples. All detected non-reference L1HS were either identified in previous studies by distinct methods, or experimentally validated (see Methods and **Figure S1G**), indicating that false-positive insertions are virtually absent (**Table S3**). Although we designed bs-ATLAS-seq to profile the youngest and human-specific L1 family, L1HS, a significant proportion of older primate-specific families (L1PA2 to L1PA8) are also amplified given the high levels of sequence identity between them and the reduced sequence complexity associated with bisulfite treatment (**Figure 1E**).

To further validate bs-ATLAS-seq, we compared the DNA methylation levels of select L1 copies obtained by this approach with results obtained by methylated DNA immunoprecipitation (MeDIP) followed by quantitative PCR (qPCR), or by direct nanopore sequencing of methylated CpGs (ONT-seq), two orthogonal techniques that do not rely on bisulfite treatment. We found a good correlation between bs-ATLAS-seq and MeDIP or ONT-seq, which independently verifies the accuracy of bs-ATLAS-seq DNA methylation values (**Figure S1H, Table S2** and **Figure S1I**). Long-read nanopore sequencing confirmed that methylation levels of the first 15 CpG reflect those of the entire L1 CpG island (**Figure S1I**). At the genome-wide level, bs-ATLAS-seq data obtained in MCF-7 cells were also cross-validated by a combination of Hi-C with methylated DNA immunoprecipitation (Hi-MeDIP) ^74^. Thus, bs-ATLAS-seq is a cost-effective method to accurately profile L1 position and methylation genome-wide.

### L1 promoter DNA methylation is cell type- and family-specific

At first sight, the profiles we obtained across the twelve cell types are globally consistent with the prevalent view that L1 DNA methylation is relaxed in cancer cells ^37, 75–77^ (**Figure S2A**). However, a closer examination of the L1 methylation landscape across cell types and L1 families offers a more refined picture and reveals a heterogeneity not detectable by aggregate analyses (**Figure 2**). In most cell types, even cancer cells, the youngest L1HS family appears hypermethylated as compared to the bulk of older L1 families (**Figure 2** and **Figure S2B**). Within the L1HS family, reference and non-reference copies (considered to include the youngest elements) show similar methylation levels (**Figure S2C**). CpG sites are progressively mutated into TpG over time owing to spontaneous deamination ^78^, and thus older L1 families contain fewer CpG sites than younger ones ^27, 79^ (**Figure S2D**). However, the fraction of methylated CpG measured by bs-ATLAS-seq for these fixed elements is unaffected by the actual number of CpG in the L1 sequence as reads are mapped against the reference genome and not a consensus sequence. Thus, our data suggest that L1 families with a lower CpG density could be more prone to inter- and intra-locus heterogeneity, as previously observed for Alu elements ^80^.

**Figure 2.**
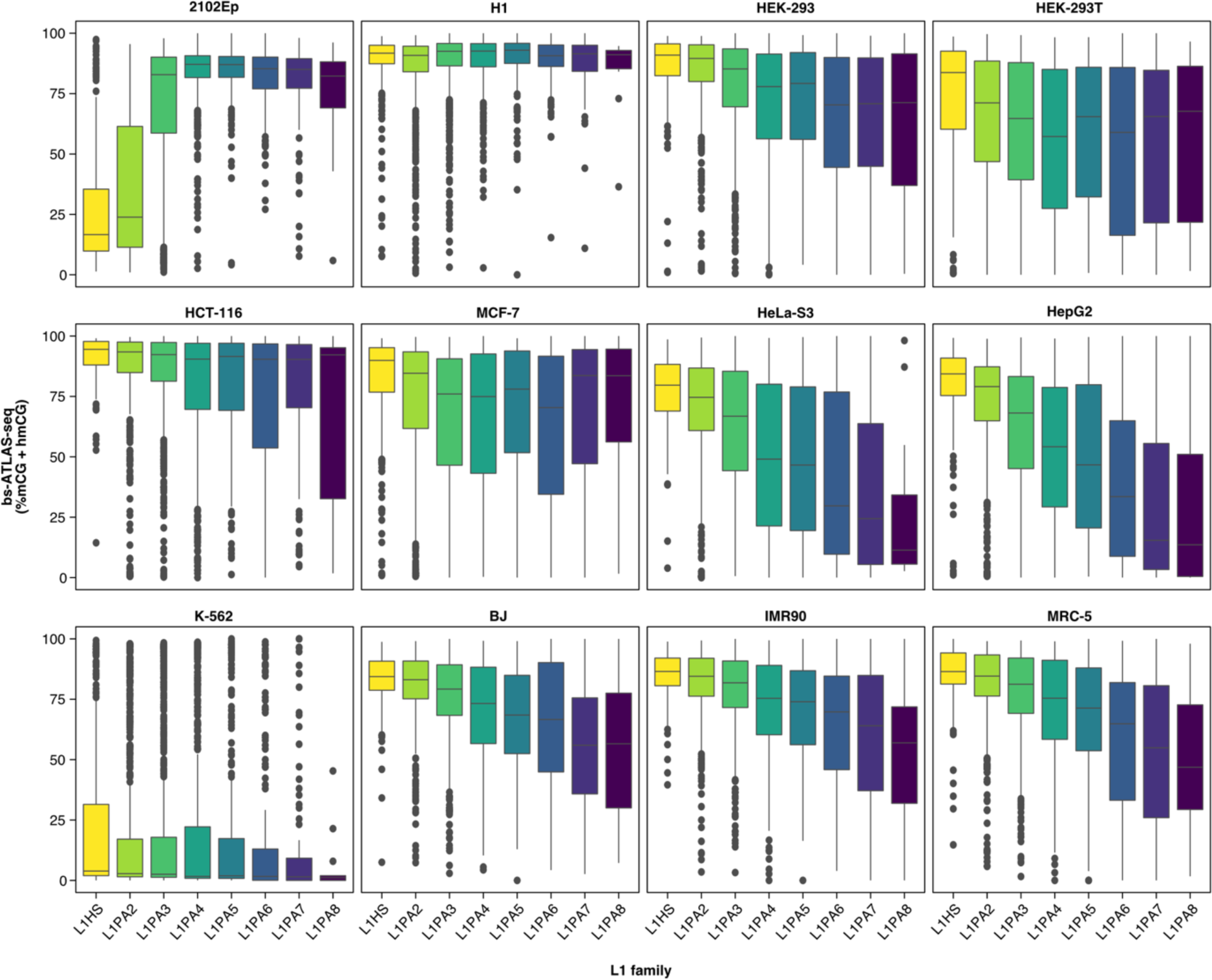
L1 promoter DNA methylation is cell-type- and family-specific. Distribution of DNA methylation levels (% mCG + hmCG) obtained by bs-ATLAS-seq, by family and cell type. Boxplots represent the median and interquartile range (IQR) ± 1.5 * IQR (whiskers). Outliers beyond the end of the whiskers are plotted individually. See also **Figure S2** and **Table S3**.

In two cell lines, the embryonal carcinoma cells 2102Ep and the chronic myeloid leukemia cells K-562, L1HS elements are exceptionally hypomethylated but with very distinct epigenetic contexts. In 2102Ep cells, hypomethylation is restricted to the young L1 families (L1HS, L1PA2, and to a lesser extent L1PA3), while older L1 elements and the rest of the genome show high levels of methylation (**Figure 2** and **Figure S2E**). In contrast, K-562 cells display a global hypomethylation phenotype that affects all L1 families and reflects genome-wide hypomethylation, down to levels as low as those observed in HCT-116 cells with *DNMT1* and *DNMT3B* inactivating mutations (**Figure 2** and **Figure S2E**).

Altogether, these data show that accessing L1 DNA methylation at individual loci reveals their heterogeneity and demonstrates that L1 DNA methylation is cell type- and family-specific.

### L1 DNA methylation can be influenced by genic activity

To test whether particular subsets of L1 elements are systematically hypo- or hyper-methylated, we compared the methylation levels of the youngest L1 loci (L1HS and L1PA2) across the different cell lines (**Figure 3A** and **Figure S3A**). Two hundred eighty-eight L1HS copies (including full-length and 3’ truncated elements) are shared by all cells. Consistent with the above results, the majority of them are highly methylated in most cell types, including cancer cells (**Figure 2** and **Figure 3A**). Excluding 2102Ep and K-562 cells, only a small subset of 59 L1HS loci shows variable methylation levels, and none is invariably unmethylated. We obtain similar results if polymorphic insertions are included (**Figure S3B**). Interestingly, in cells where L1HS elements are globally unmethylated (2102Ep and K-562), we still observe methylated copies (n=54 and n=46, respectively) (**Figure 3A**).

**Figure 3.**
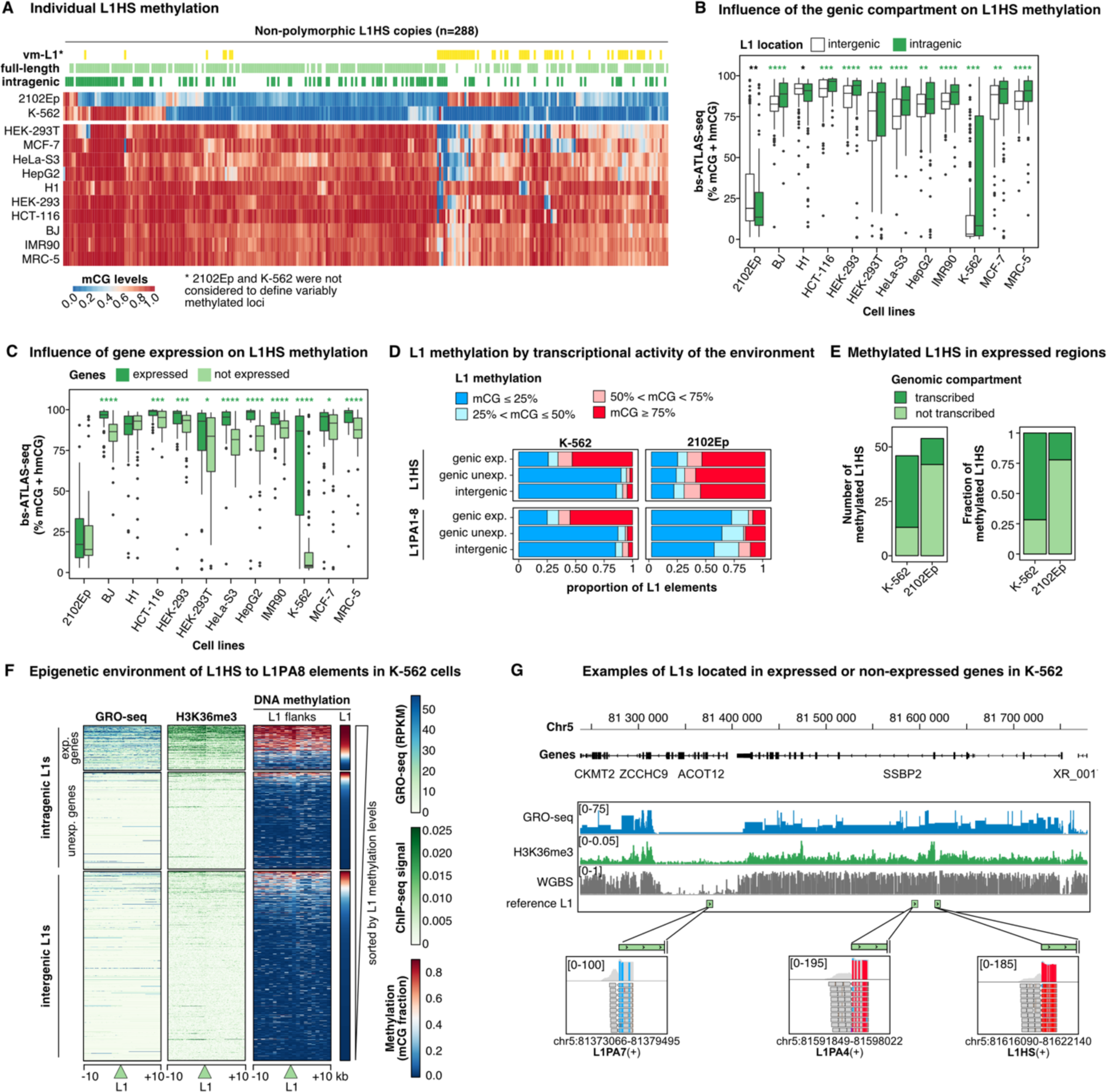
L1HS promoter DNA methylation is locus-specific and influenced by the local environment. **(A)** Heatmap of bs-ATLAS-seq methylation levels (% mCG + hmCG, from blue for unmethylated to red for fully methylated) for individual L1HS loci across a panel of 12 cell lines. Each column represents an L1HS locus. Only loci shared by all cell lines are shown here. For a heatmap including polymorphic L1HS elements, see **Figure S3B**. Rows and columns were ordered by hierarchical clustering based on methylation values. Intragenic L1HS are indicated in dark green above the heatmap and full-length L1HS copies in light green. Variably methylated L1 elements (vm-L1s, yellow) were defined as showing a greater difference between the second highest and the second lowest values of DNA methylation at this locus. K-562 and 2102Ep values were not considered to define vm-L1s. **(B)** Comparison of L1HS methylation levels obtained by bs-ATLAS-seq for intra-(green) vs inter-genic (white) elements. **(C)** Comparison of the methylation levels of intragenic L1HS obtained by bs-ATLAS-seq for expressed (dark green; TPM≥1) vs non-expressed genes (light green; TPM<1) **(D)** Proportion of L1 elements in different methylation level categories according to their genic environment and activity in K-562 (left) or 2102Ep (right) cells. Note that L1PA1 is synonymous with L1HS. **(E)** Distribution of methylated L1HS elements in transcribed (dark green) vs non-transcribed (light green) genomic compartments in K-562 and 2102Ep cells. **(F)** Heatmaps illustrating nascent transcription (GRO-seq), H3K36me3 histone modifications (ChIP-seq), and DNA methylation (whole genome bisulfite sequencing, WGBS), in 10 kb-windows upstream and downstream L1 elements from the L1HS to the L1PA8 family (green triangle). Loci are separated according to their position relative to genes (left), separating expressed and unexpressed genes, and sorted by decreasing L1 methylation levels (bs-ATLAS-seq, right). Data are shown in 10 bp-bins. A similar heatmap restricted to the L1HS family is shown in **Figure S3C**. **(G)** Genome browser view of a region on chromosome 5 encompassing three full-length L1 elements (green rectangles) with distinct promoter DNA methylation profiles, integrated with nascent transcription (GRO-seq), H3K36me3 histone modifications (ChIP-seq), and DNA methylation (WGBS) datasets in K-562 cells. The bottom inserts show L1 methylation as measured by bs-ATLAS-seq. The methylated L1HS and L1PA4 elements are inserted in a transcribed gene marked by high H3K36me3 and DNA methylation levels over the entire gene body, while the unmethylated L1PA7 is inserted in an unexpressed gene with low gene body DNA methylation and H3K36me3 levels. In panels (B) and (C), boxplots represent the median and interquartile range (IQR) ± 1.5 * IQR (whiskers). Outliers beyond the end of the whiskers are plotted individually. *p < 0.05, **p < 0.01, ***p < 0.001, and ****p < 0.0001, one-sided two-sample Wilcoxon test (green for increase and black for decrease). See also **Figure S3** and **Table S3**.

To understand why individual loci adopt a methylation profile distinct from the bulk of L1HS, we compared their genomic environment with that of unmethylated copies. In K-562 cells, methylated L1HS elements are enriched in genes (**Figure 3A**). L1HS elements inserted in genes are more methylated than in intergenic regions (**Figure 3B**), and those in expressed genes are more methylated than in unexpressed genes (**Figure 3C**). Moreover, two thirds of L1HS are methylated in expressed genes, but less than 10% are methylated in non-transcribed compartments (non-expressed genes or intergenic regions, **Figure 3D**, top left). This contrasts with 2102Ep cells, which exhibit a similarly low proportion of methylated L1HS loci as the K-562 cells (**Figure 3A**), but for which the relative proportion of methylated and unmethylated copies remains similar regardless of the transcriptional state of the integration locus (**Figure 3D**, top right). Similar results were obtained when also including L1PA2 to L1PA8 families (**Figure 3D**, bottom). In absolute numbers, 33 out of 46 (72%) of hypermethylated L1HS (mCG≥75%) are located in a transcribed genomic compartment in K-562 cells, but only 12 out of 54 (22%) in 2102Ep cells (**Figure 3E**).

In mammals, DNA methylation is widely targeted to gene bodies through transcription-coupled deposition of H3K36me3 histone marks and the subsequent recruitment of the DNA methyltransferase Dnmt3b ^81–84^. Our observation thus suggests that, in K-562 cells, most L1HS DNA methylation results from their co-transcription with genes as previously described for intragenic CpG islands ^85^. Consistently, methylated L1s are predominantly embedded in transcribed regions enriched in H3K36me3 histone modification and DNA methylation (**Figure 3F, Figure 3G** and **Figure S3C**). Thus, at the global level, L1HS elements are not directly targeted by DNA methylation in K-562 cells, but locally, some are methylated due to their co-transcription with genes. We also observe a significant - yet modest - association of genic transcription with L1HS DNA methylation in all other but embryonic cell types (**Figure 3B** and **Figure 3C**), in agreement with the previous observation showing no correlation between gene body methylation and expression in embryonic stem cells ^82^. We note that high levels of young L1 DNA methylation in the H1 embryonic stem cells, which were grown in medium containing LIF and serum, are consistent with a primed state, as previously reported ^86^.

### L1 elements drive local but short-range epivariation

Transposable elements have been proposed to function as “methylation centers” from which methylation can propagate into the flanking sequences in many species, including mammals (REF Turker). However, this possibility has never been tested systematically for L1 elements, particularly in human cells. Bs-ATLAS-seq provides not only the DNA methylation state of L1 promoters but also that of proximal CpGs upstream of L1 elements, giving us the opportunity to assess the association between L1 DNA methylation and that of its 5’ flanking genomic region. DNA methylation of L1 upstream sequences is remarkably similar to the methylation of the L1 promoter itself (**Figure S4A**). Unmethylated L1HS to L1PA4 are associated with a hypomethylated 5’ flank. Conversely, methylated elements are associated with methylated upstream sequences. For L1HS and L1PA2, hypomethylation of the flank progressively decreases as the distance from L1 increases and disappears around 300 bp, a situation previously observed for transgenic CpG islands in reporter genes and termed “sloping shores” ^87^.

To investigate whether L1 elements instruct the methylation state of their flanks, we compared allelic epivariation in the region upstream of L1 at heterozygous loci (*i.e.* having one filled allele - with L1 - and one empty allele - without L1 - in the same cell line). By design, bs-ATLAS-seq only probes the filled alleles. Therefore, to gain access to the methylation states of both alleles at heterozygous loci, and to extend the region analyzed around the L1 promoter, we performed long-read nanopore Cas9-targeted sequencing ^88^, with direct calling of 5-methyl-cytosine (5mC) (**Figure 4A**). Sequencing starts at the single guide RNA (sgRNA) binding site in the region downstream of L1 and common to both alleles, and progresses towards the target L1 in the antisense direction ^89^. By applying this strategy to a subset of 124 potentially polymorphic loci in 4 cell lines (2102Ep, MCF-7, K-562 and HCT116), we genotyped and obtained the methylation profile of each allele, including the entirety of the L1 element when present, and several kilobases upstream, with a mean coverage of 45x (**Figure 4A** and **Table S4**). The target L1s were selected as present in less than half of the 12 cell lines of the panel and therefore more likely to be heterozygous. On average, 40 loci are heterozygous (40 ± 4; mean ± s.d.) and 21 are homozygous and filled (21 ± 0.5) in each cell type, with a total of 162 heterozygous alleles and 83 homozygous filled alleles sequenced (**Figure 4C** and **Table S4**). In agreement with bs-ATLAS-seq results, we observe that cell lines differ by their L1 methylation profiles and levels (**Figure S1I**) and that L1 and their proximal upstream sequence adopt a similar methylation state (**Figure S4B**). Moreover, unmethylated L1 promoters surrounded by hypermethylated distant genomic flanks, such as those frequently observed in 2102Ep cells, inform us on the extent of the non-methylated domain within L1 elements and on the width of sloping shores. Methylation starts decreasing approximately 300 bp upstream of L1 to reach a minimum at the beginning of the L1 promoter. Methylation stays low over the first 500 bp of L1 5’ UTR, and increases again to reach a plateau around position +800 with high levels of methylation over the entire L1 body (**Figure S4B** and **Figure 4A**).

**Figure 4.**
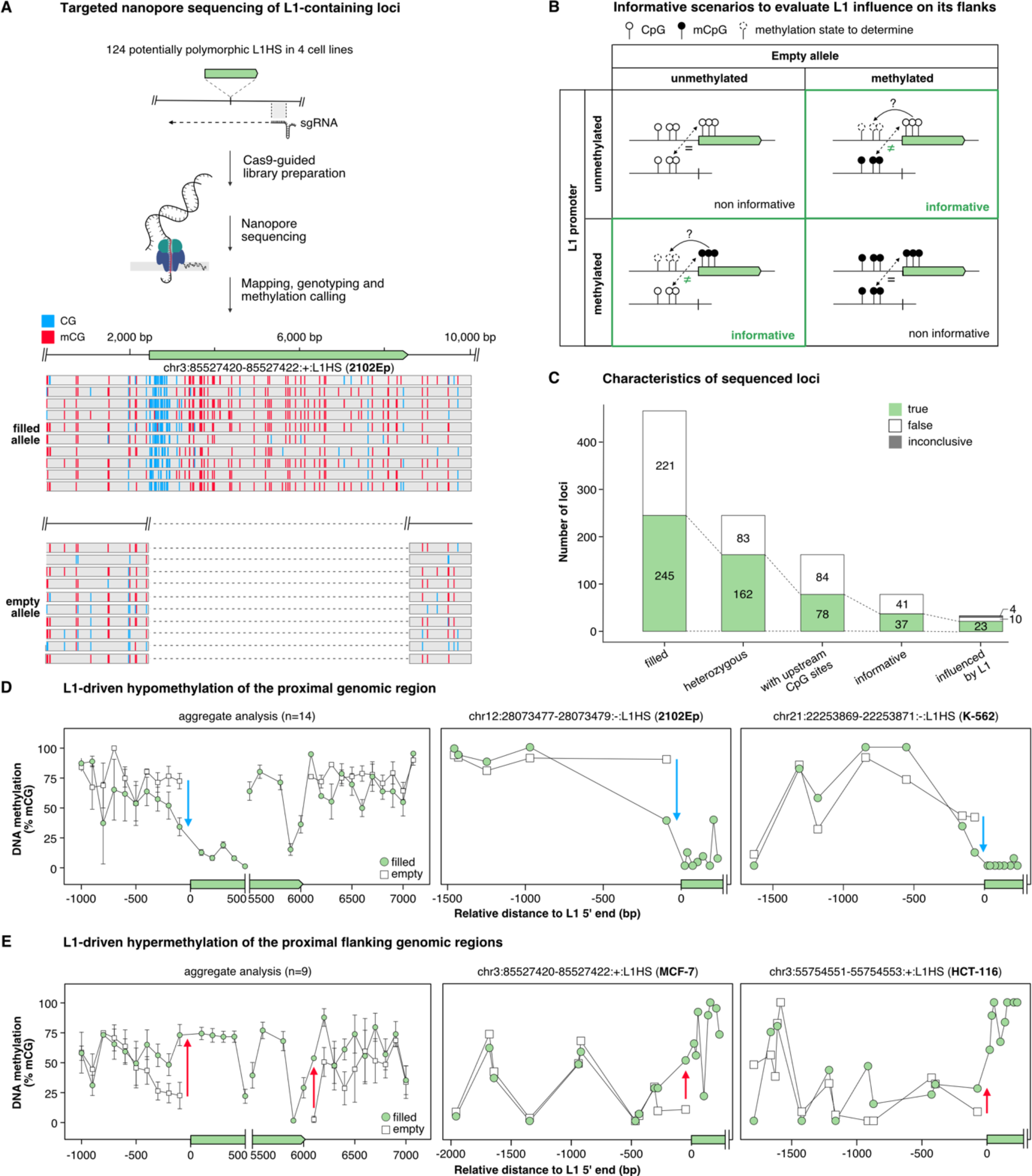
L1s frequently induce proximal epivariation. **(A)** Scheme summarizing the strategy to genotype and assess DNA methylation profiles of loci containing polymorphic L1HS elements by Cas9-guided nanopore sequencing. Single-guide RNAs (sgRNAs) designed to bind ∼ 1 kb downstream of each L1HS insertion (green solid arrows), were synthesized *in vitro* as a pool and assembled with recombinant Cas9. Cleavage of genomic DNA with the pool of Cas9 RNPs allows the subsequent ligation of sequencing adapters downstream of L1s and targeted nanopore sequencing of the selected loci (see Methods). As an example, a genome browser screenshot is shown for the insertion chr3:85527420-85527422:+:L1HS:NONREF in 2102Ep cells (top: filled allele, bottom: empty allele). The targeted L1HS (green), as well as methylated (red) and unmethylated (blue) CpG are indicated. Only the first 10 reads are shown for each allele, and the 3’ end of the reads (on the left) have been truncated for layout. **(B)** Theoretical methylation profiles at heterozygous L1 insertions (green solid arrow). The influence of L1 methylation on the surrounding genomic region can only be detected in situations where the methylation state of L1 differs from that of the empty locus (assumed to represent the pre-integration state). For these informative scenarios, the DNA methylation level of the region upstream of L1 is then analyzed and compared to the empty locus. **(C)** Characteristics of the loci profiled by nanopore sequencing. The y-axis represents the number of loci characterized in the 4 cell lines as filled, heterozygous and with CpG sites upstream of L1 (∼ 300 bp). Among them, 37 loci were considered as informative: 23 with a profile consistent with L1-induced epivariation, 10 showing no evidence of L1-induced epivariation, and 4 being inconclusive (see Methods). **(D)** L1-driven hypomethylation of the proximal upstream genomic region. Left, average DNA methylation levels in 100 bp-bins at loci with an upstream slopping shore (n=14, mean ± s.d.). Middle and right, examples of L1-induced upstream flank hypomethylation. **(E)** L1-driven methylation of the proximal flanking genomic region. Left, average DNA methylation levels in 100 bp-bins at loci with DNA methylation spreading from L1 to the external flanks (n=9, mean ± s.d.). Middle and right, examples of L1-induced flanking sequence DNA methylation. For (**D**) and (**E**), the empty (white squares) and filled (green circles) alleles are overlaid. Blue and red arrows denote hypo- and hyper-methylation relative to the empty locus, respectively. The x-axis represents the relative distance to L1 5’ end (bp) and the y-axis the percentage of DNA methylation. See also **Figure S4** and **Table S4**.

Next, we systematically examined potential L1HS-driven allelic epivariation. We reasoned that the influence of an L1 element on the flanking region could only be assessed if the methylation state of the L1 promoter differs from that of its cognate empty allele (**Figure 4B**). Out of 78 heterozygous loci, 37 were found informative as they exhibit a significant difference of DNA methylation levels between the L1 promoter and the empty allele (> 30 %) and possess at least one CpG site 300 bp upstream of the insertion site (**Figure 4C**). Upon manual inspection, 4 loci were further excluded as they show high variability and were inconclusive. Remarkably, more than two third of the remaining loci (23/33) exhibit short-range L1-mediated epivariation. Fourteen hypomethylated L1 elements have sloping shores (demethylation of the proximal upstream CpGs, **Figure 4D**), while 9 hypermethylated elements propagate methylation in their upstream sequence (**Figure 4E**). Of note, methylation spreading was detected at both extremities of methylated L1s (**Figure 4E**, left, and **Figure S4C**), while sloping shores were only observed at the 5’ junction of unmethylated L1s. In most cases, we can confidently conclude that the presence of the L1 insertion directly causes the observed allelic epivariation, as epivariation is not just associated with the filled locus, but also forms a gradient starting from the L1 element. For one locus in K-562 cells, we observe broad allelic epivariation between the empty and filled alleles, but it is unclear whether the methylation difference is due to the presence or absence of the L1HS insertion, or simply reflects the inclusion of an L1 element in an already existing epiallele (**Figure S4D** and **Figure S4E**). Finally, for 10 L1 elements, we did not detect variation of methylation between the filled and the empty alleles (**Figure S4F**). Thus, L1HS elements exert a short-range but frequent epigenetic influence on their genomic environment, which can create local epivariation.

### Local epivariation at L1 loci is associated with distinct transcription factor landscapes

To further understand what differentiates methylated from unmethylated L1 loci, we examined whether these subsets could be associated with distinct transcription factor (TF) binding profiles. We performed differential TF binding site enrichment analysis, comparing the 5’ junction of unmethylated versus methylated L1. The analyzed region spans the 300 bp of upstream sequence under L1 epigenetic influence, as well as the first 500 bp of the L1 5’ UTR which appears to be variably methylated (**Figure S4B** and **Figure 5A**). For this purpose, we screened the entire Unibind database ^90^, a catalogue of high-confidence TF binding sites (TFBS) predictions based on ∼3500 publicly available chromatin-immunoprecipitation and sequencing (ChIP-seq) experiments from various tissues and cell types, including those from ENCODE (**Figure 5A**). The number of unmethylated L1HS being relatively small in most cell types, we also included the L1PA2 family to increase the statistical power of this analysis, as it is the most recent primate-specific family after the L1HS family (**Figure S3A**) and shows a similar pattern of proximal upstream epivariation (**Figure S4A**). Note that for each cell line in our panel, we compared the methylated and unmethylated L1 subsets to all ChIP-seq data stored in Unibind, irrespective of the cell-type or conditions in which they were obtained. The rationale is that even if datasets from our cell line of interest are not present in Unibind, a similar cell type or condition may be represented. Indeed, the screen revealed several TFs specifically associated, in at least one biological condition, with the subset of methylated or unmethylated L1s found in one of the cell lines of the panel (**Figure 5B**). Closer examination of the motifs underneath the ChIP-seq peaks indicates that some of the binding sites are internal to the L1 promoter (such as for YY1), and some are found in both the upstream and internal sequences (such as for ESR1, FOXA1, or CTCF) (**Figure 5C**) ^91, 92^.

**Figure 5.**
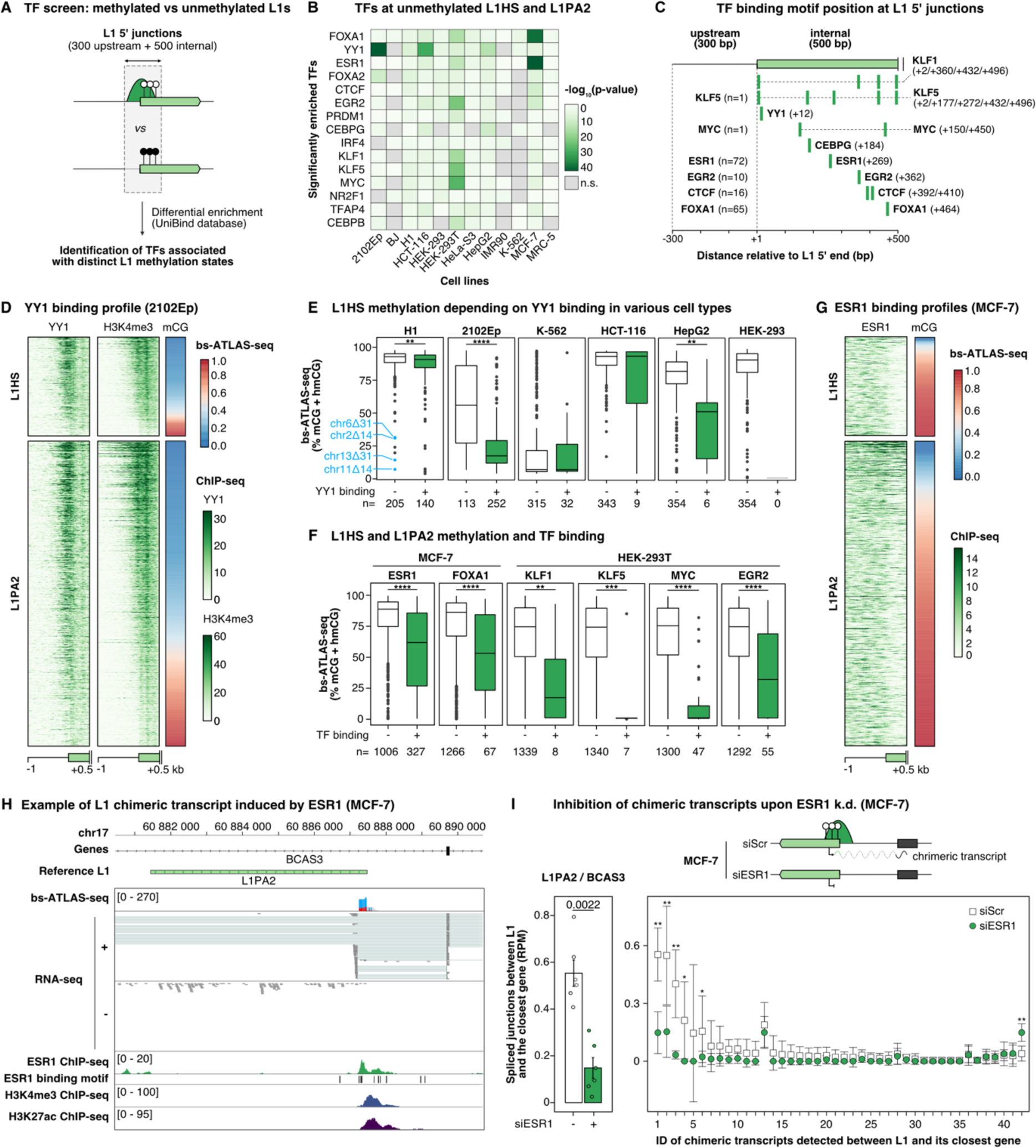
Unmethylated L1 and their flanking sequences are bound by a specific set of transcription factors (TFs) **(A)** Screening strategy to identify TFs differentially associated between unmethylated and methylated L1HS and L1PA2 copies using curated datasets publicly available in the UniBind database ^90^. Note that for each cell line in our panel, we compared each pair of methylated and unmethylated L1 subsets to all ChIP-seq data stored in Unibind (∼3500 datasets), irrespective of the cell-type or conditions in which they were obtained. The rationale was that even if our specific cell line is not necessarily represented in Unibind datasets, a similar cell type may be represented. The main hits were then subsequently confirmed using matched datasets (see panels E and F). **(B)** Heatmap showing the TF binding enrichment at hypomethylated L1HS and L1PA2 in our panel of cell types. Only the 15 most enriched TFs are shown. **(C)** Schematic representation of the location of the motifs corresponding to the TFs identified in (B). For TFs binding upstream of L1 insertions, the number of loci with an upstream peak is indicated. **(D)** Heatmap displaying L1 methylation (bs-ATLAS-seq), as well as YY1 and H3K4me3 binding (ChIP-seq), at the 5’ junction (-1 to +0.5 kb) of L1HS and L1PA2 elements in 2102Ep cells. Loci are sorted by increasing levels of L1 methylation. ChIP-seq signal represents the number of normalized reads per 10-bp bin. **(E)** DNA methylation level of L1HS bound (+) or unbound (-) by YY1 in embryonal cell lines (H1 and 2102Ep) and other cell lines for which matched YY1 ChIP-seq were also publicly available (K-562, HCT116, HepG2, HEK-293T). The number of L1HS copies in each subset (n) is indicated at the bottom of the plot. In H1 cells, the four hypomethylated loci in blue refer to those studied in ^63^. **(F)** DNA methylation levels of L1HS and L1PA2 loci bound (+) or not bound (-) by ESR1, FOXA1, KLF1, KLF5, Myc and EGFR2 in the relevant cell types. ChIP-seq data are matched to the cell line. The number of L1HS copies in each subset (n) is indicated at the bottom of the plot. **(G)** Heatmap displaying L1 methylation (bs-ATLAS-seq), as well as ESR1 binding (ChIP-seq), at the 5’ junction (-1 to +0.5 kb) of L1HS and L1PA2 elements in MCF-7 cells. Loci are sorted by increasing levels of L1 methylation. ChIP-seq signal represents the number of normalized reads per 10-bp bin. **(H)** Genome browser view of the *BCAS3* locus integrating L1 methylation (bs-ATLAS-seq), expression (poly(A)^+^ RNA-seq), ESR1 binding, as well as H3K4me3 and H3K27ac histone modifications (ChIP-seq). Note the distinctive spliced RNA-seq reads, antisense relative to the L1 element, linking L1 antisense promoter with the adjacent *BCAS3* exon. **(I)** SiRNA-mediated knock-down of ESR1 leads to reduced L1 chimeric transcripts. Top, schematic representation of chimeric transcripts initiated from L1 antisense promoter and leading to truncated or alternative isoforms of the surrounding gene. Upon siRNA-mediated knock down (siESR1), the number of L1 chimeric splice junctions is expected to decrease if ESR1 drives chimeric transcript synthesis, as compared to a scrambled siRNA control (siScr). Bottom left, chimeric transcripts at the *BCAS3* locus quantified by the number of normalized spliced-RNA-seq reads (RPM) detected in MCF-7 cells treated by an siRNA against ESR1 (+) or a control scrambled siRNA (-) (data from GSE153250). Bars represent the mean ± s.d. (n=6) and are overlaid by data of individual replicates (one-sided two-sample Wilcoxon test). Bottom right, average chimeric transcripts quantified as the normalized number of splice junctions between L1 and its closest gene in RPM for 42 loci (n=6, mean ± s.d.). The 42 loci are sorted by descending order according to the difference of chimeric transcript levels between cells treated by siESR1 and the control siScr, and the associated number corresponds to the transcript ID in **Table S5**. For 37 loci out of 42 (88%), L1 chimeric transcription is reduced upon ESR1 knock down. The difference is statistically significant for 5 loci (one-sided two-sample Wilcoxon test). In panels (E) and (F), boxplots represent the median and interquartile range (IQR) ± 1.5 * IQR (whiskers). Outliers beyond the end of the whiskers are plotted individually. *p < 0.05, **p < 0.01, ***p < 0.001, and ****p < 0.0001, two-sided two-sample Wilcoxon test. See also **Figure S5** and **Table S5.**

Among the top hits, YY1 is strongly enriched at the unmethylated subset of L1s from 2102Ep cells (**Figure 5B**). The YY1 binding site is internal to the L1 promoter, but close enough to the 5’ end (position +10 ± 2 bp) that the ChIP-seq signal largely overlaps the flanking sequence. The strong YY1 enrichment at unmethylated L1s from 2102Ep cells was detected in Unibind datasets obtained in different cell types, including two embryo-related cell lines, the H1 embryonic stem cells and NTera2/D1 embryonal carcinoma cells. To confirm this association in matched cell types, we performed YY1 ChIP-seq in 2102Ep cells (**Figure 5D**) and tested if it was also observed in other cell types of the panel for which YY1 ChIP-seq data were publicly available (**Figure 5E**). Although YY1 is relatively ubiquitously expressed (**Figure S5A**), its binding to L1HS elements mostly occurs in the embryonal cell types, H1 and 2102Ep, where YY1-bound L1HS elements represent a large fraction of all L1HS elements (**Figure 5E**, 41% and 69%, respectively). In these cells, YY1-bound L1HS elements are significantly less methylated as compared to their YY1-unbound counterparts (**Figure 5D** and **Figure 5E**). Among other cell types, only the hepatoblastoma cell line HepG2 shows a significant difference of methylation between YY1-bound and -unbound elements, but overall, the number of bound elements is extremely limited in these cells (**Figure 5E**, 9% in K-562, and <3% in HCT-116 and HepG2, 0% in HEK-293). Interestingly, the binding of YY1 is not restricted to L1HS elements. A large fraction of L1PA2 elements in H1, and the majority of them in 2102Ep cells, are bound by YY1, and again bound elements are significantly less methylated than unbound ones (**Figure S5B**). By tracking an L1 progenitor active in the brain and carrying a small 5’ truncation spanning the YY1 motif (chr13Δ31_L1_), a previous report suggested that the YY1 binding site is involved in L1 silencing, and thus proposed that YY1, or another pathway acting on its binding site, may drive L1 methylation ^63^. As YY1 was best known to activate the L1 promoter, helping to define accurate L1 transcription start site ^92–96^, this finding was unexpected. We confirmed that the same locus, as well as a small set of other elements lacking the YY1 motif, are hypomethylated in the embryonal cell types H1 and 2102Ep (**Figure 5E**, blue labeled data points, and **Figure S5C**). However, if we consider L1HS elements that are actually bound by YY1, and not only those that contain the YY1 motif, we observe that YY1 binding is in fact strongly associated with L1 hypomethylation (**Figure 5D** and **Figure 5E**). Finally, copies bound by YY1 are also marked by active chromatin marks (H3K4me3) (**Figure 5D**) and are more expressed in 2102Ep as compared to unbound elements (**Figure S5D**). Altogether, these results reinforce the idea that, genome-wide, YY1 binding is not associated with the repression of L1s, but is instead associated with L1 hypomethylation and transcriptional activity, at least in embryonic stem cells or embryonal carcinoma cells. Thus, our data are consistent with a scenario whereby alterations of the YY1 motif may incidentally affect other pathways targeting the same sequence and leading to L1 hypomethylation ^63^.

Another prominent hit in our TF search was the estrogen receptor ESR1, which is strongly enriched at unmethylated loci in the MCF-7 breast cancer cell line (**Figure 5B**). Unlike YY1, ESR1 binds not only to the internal L1 sequence, but also to the upstream region (**Figure 5C, Figure 5G** and **Figure S5G**). Altogether, almost one fourth of the loci are bound by ESR1 (**Figure 5F**). The expression level of this TFs is higher in MCF-7 as compared to other cell types (**Figure S5A**), and L1 elements associated with ESR1 binding are less methylated (**Figure 5F**). Among the 327 L1 elements with ESR1 binding, we detected 42 L1 chimeric transcripts with a neighboring gene (**Figure 5H-I** and **Table S5**). These chimeric transcripts engage both protein-coding RNA and lncRNA species, many of them being associated with cancer, either as biomarkers or as oncogenes, and can encode tumor-specific antigens such as L1-GNGT1 ^13, 49, 97, 98^. As an example, the 5’ UTR of an L1PA2 element located in the intron of the *BCAS3* gene is unmethylated, and covered by active chromatin marks (H3K4me3 and H3K27ac), while its 5’ UTR and immediate upstream region (-189 bp) are bound by ESR1 (**Figure 5H** and **Figure 5I**, left). Moreover, RNA-seq data in MCF-7 show a high proportion of spliced reads between the L1PA2 antisense promoter and the closest *BCAS3* intron, indicating that L1 demethylation and ESR1 binding are associated with antisense promoter activity, which can act as an alternative promoter for *BCAS3*. Of note, unit-length L1 transcripts are also detected at this locus, suggesting that the L1 sense promoter is also active. Finally, when ESR1 expression is experimentally reduced by siRNA-mediated knockdown ^99^, the expression of L1 elements themselves (**Figure S5F**) and their chimeric transcripts (**Figure 5I**) decreases. Given the importance of estrogen receptor (ER) status in breast cancer prognosis and management, we explored data from the Pan-Cancer Analysis of Whole Genomes (PCAWG) project to test whether ER status might be associated with increased L1 mobilization in this cancer type ^48, 100^. We found that ER^+^ tumors more frequently exhibit somatic L1 retrotransposition than ER^-^ tumors (**Figure S5H**, left). Nevertheless, the number of events is not significantly different between the two groups (**Figure S5H**, right). Consistent with the detection of L1 ORF1p in more than 90% of all breast adenocarcinoma cases, irrespective of their ER status^101, 102^, our results suggest that L1 might be activated through different sets of transcription factors in ER^-^ and ER^+^ tumors. Altogether we conclude that ESR1 directly drives L1 sense and antisense promoter activities in MCF-7 cells, and more generally, that TFs bound within or next to unmethylated L1s can drive cell-type-specific functional alterations of neighboring genes.

Finally, other TFs significantly enriched at hypomethylated L1s were bound to 5% or less of the whole set of L1HS and L1PA2 elements (**Figure 5F**). Among them, FOXA1 is highly expressed in MCF-7 cells (**Figure S5A**). As FOXA1 has pioneer activity and can drive distance-dependent local demethylation ^103^, we speculate that cell-type-specific FOXA1 expression and binding could lead to the hypomethylation of a small subset of L1 loci.

### L1HS promoter hypomethylation is not sufficient to promote expression at most loci

It is broadly accepted that hypomethylation of L1 leads to their transcriptional reactivation, at least at the global level ^39, 73^. To test this assumption at individual loci, we performed poly(A)^+^ RNA-seq for the cell lines of the panel since these cells show a broad range of L1 expression ^9^. Of note, RNA samples were prepared from cells collected from the same plate as for bs-ATLAS-seq to match methylation and expression data, with the exception of the embryonic stem cell H1 for which we used publicly available data (due to regulatory restrictions). Unambiguously measuring L1HS expression levels at individual loci is extremely challenging due to: (i) the low mappability of these repeated sequences; (ii) L1HS insertional polymorphisms; and (iii) pervasive transcription of L1 embedded in genes that greatly exceeds the autonomous transcription of unit-length L1 elements ^34^. To recognize autonomous L1 expression driven by the L1 promoter, we applied a previously devised strategy which measures readthrough transcription downstream of reference and non-reference L1HS elements, after removing potential signal from gene transcription or pervasive transcription ^9^ (**Table S6**). To exclude that some expressed copies could escape detection due to a strong polyadenylation signal at – or close to – an L1 3’ end, we also assessed L1 expression by L1EM ^104^, a software relying on the expectation-maximization algorithm to reassign multi-mapping reads (**Table S6**).

Irrespective of the strategy used to identify expressed loci, we found that only a limited number of them are expressed in a given cell type, consistent with our previous findings ^9^ (**Figure 6A, Figure 6B** and **Table S6**). Of note, some show intact ORFs and published evidence of retrotransposition capability, as measured by cell culture assays or through the identification of transduction events deriving from the locus (**Figure 6C** and **Table S6**). As expected, we observe a weak but significant anti-correlation between L1HS methylation and expression in most cell types expressing detectable levels of L1HS (**Figure 6A** and **Table S6**). Hence, the top-expressed loci are unmethylated (**Figure 6A** and **Figure 6D**, top left) while fully methylated loci are not expressed (**Figure 6A** and **Figure 6D**, top right). However, most unmethylated loci show no evidence of expression, indicating that hypomethylation of L1HS elements alone is insufficient to permit their expression (**Figures 6A, Figure 6B** and **Figure 6D**, bottom left). K-562 cells represent an extreme case of this configuration as only one single element shows detectable expression while most copies are hypomethylated. Inversely, some L1HS with relatively high levels of methylation are expressed in MCF-7 and HepG2 cells (**Figure 6A** and **Figure 6D**, bottom right). Methylated copies could escape silencing by using upstream alternative promoter ^63, 105^. Alternatively, epiallele heterogeneity in the cell population or even between alleles could account for their expression. Indeed, even if these elements are in the upper range of methylation, they show a significant proportion of unmethylated reads (**Figure 6D**, bottom right). Beside the activity of the L1 sense promoter, we also detect antisense transcription, and some unmethylated L1HS elements only produce antisense transcripts (**Figure 6D**, bottom left). These observations suggest that the absence of DNA methylation at L1HS elements may not always be sufficient to trigger their transcriptional activation.

**Figure 6.**
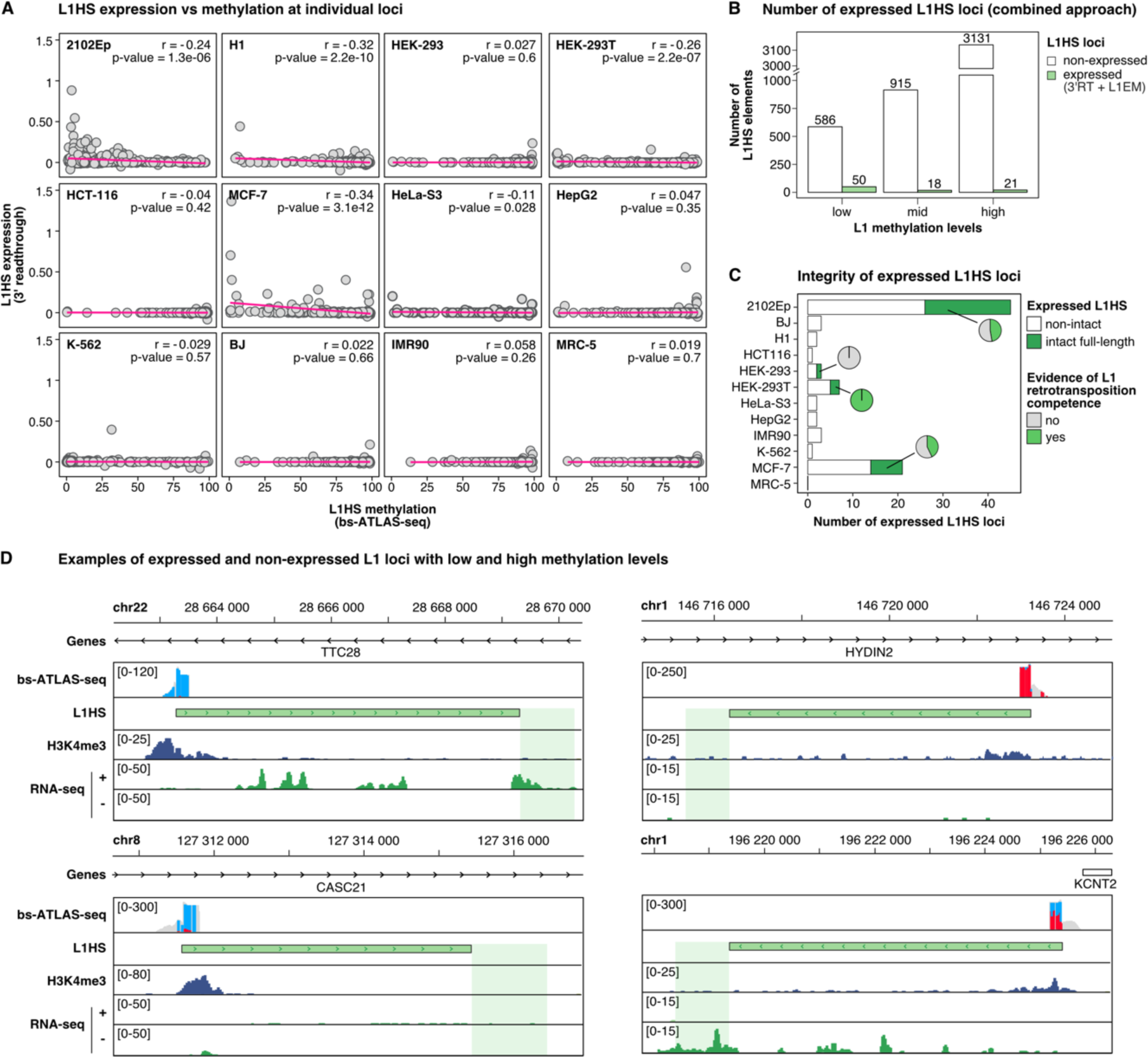
L1HS promoter hypomethylation is not sufficient to trigger L1HS expression at the locus level. **(A)** Genome-wide correlation between L1HS expression and methylation at individual L1HS loci in the entire cell line panel. L1 expression level is estimated through 3’ readthrough transcription, a consequence of the weak L1 polyadenylation signal ^9^. Briefly, we calculated the number of unique RNA-seq reads mapped within a 1 kb-window downstream of L1 and on the same strand, and subtracted the number of unique reads mapped within a 1 kb-window upstream of L1 to eliminate signal from surrounding pervasive transcription. Then, the value was normalized by the total number of mapped reads (RPKM). Negative values were set to 0. Most L1HS are not expressed and hypermethylated (lower right corners). r and p represent Pearson’s correlation coefficient and p-value, respectively. A regression line is indicated in pink. **(B)** Barplots indicating the absolute number of L1HS elements with low (mCG ≤ 25%), medium (25% < mCG < 75%) or high (mCG ≥ 75%) methylation levels obtained by bs-ATLAS-seq, and detected as unexpressed (white) or expressed (light green) by the combined 3’ readthrough (3’ RT) and L1EM ^104^ approaches. **(C)** Barplots indicating the absolute number of expressed L1HS elements across the different cell lines, for non-intact (white) and intact copies (dark green). Among the expressed L1 elements, the associated pie charts show the proportion of copies with published evidence of retrotransposition competence. **(D)** Genome browser views of 4 L1HS loci with distinct configuration of DNA methylation and expression in MCF-7 cells. L1 DNA methylation (bs-ATLAS-seq) is integrated with poly(A)^+^ RNA-seq and H3K4me3 ChIP-seq data. An unmethylated and expressed (top left), a methylated and unexpressed (top right), an unmethylated and unexpressed (bottom left) and a half-methylated and expressed L1HS (bottom right) are shown. See also **Table S6**.

To experimentally test this hypothesis, we measured L1HS methylation and expression in HCT-116 cells treated by the DNA methyltransferase inhibitor 5-aza-2’-deoxycytidine (5-aza) ^106^ (**Figure 7**). Under our experimental conditions, 5-aza homogenously decreases methylation by 50% for all L1HS elements (**Figure 7A**), with completely unmethylated reads at most loci (**Figure 7B**), as a likely consequence of passive loss of DNA methylation during replication. As expected, this acute and massive reduction of DNA methylation allows the reactivation of several transposable element families including L1HS to L1PA8 (**Figure 7C**). However, it appears that only a subset of L1HS copies (22%, 85 out of 379 loci) are detected as expressed (**Figure 7D**). 5-aza-induced demethylation acts not only on promoter regions, but also on gene bodies, with opposing effects on gene expression ^107^ – and thus possibly on the expression of intragenic L1 elements. To exclude unpredictable effects of gene body demethylation on intragenic L1 expression, we analyzed separately L1 loci located outside genes, within genes that are expressed, and within genes that are not expressed (**Figure 7E**). The result shows that the proportion of derepressed L1 loci upon 5-aza treatment is slightly superior in the expressed genes as compared to non-transcribed regions. However, overall, the majority of L1 elements stay silent. We thus conclude that acute demethylation is not sufficient at most L1 loci to ensure their transcriptional reactivation.

**Figure 7.**
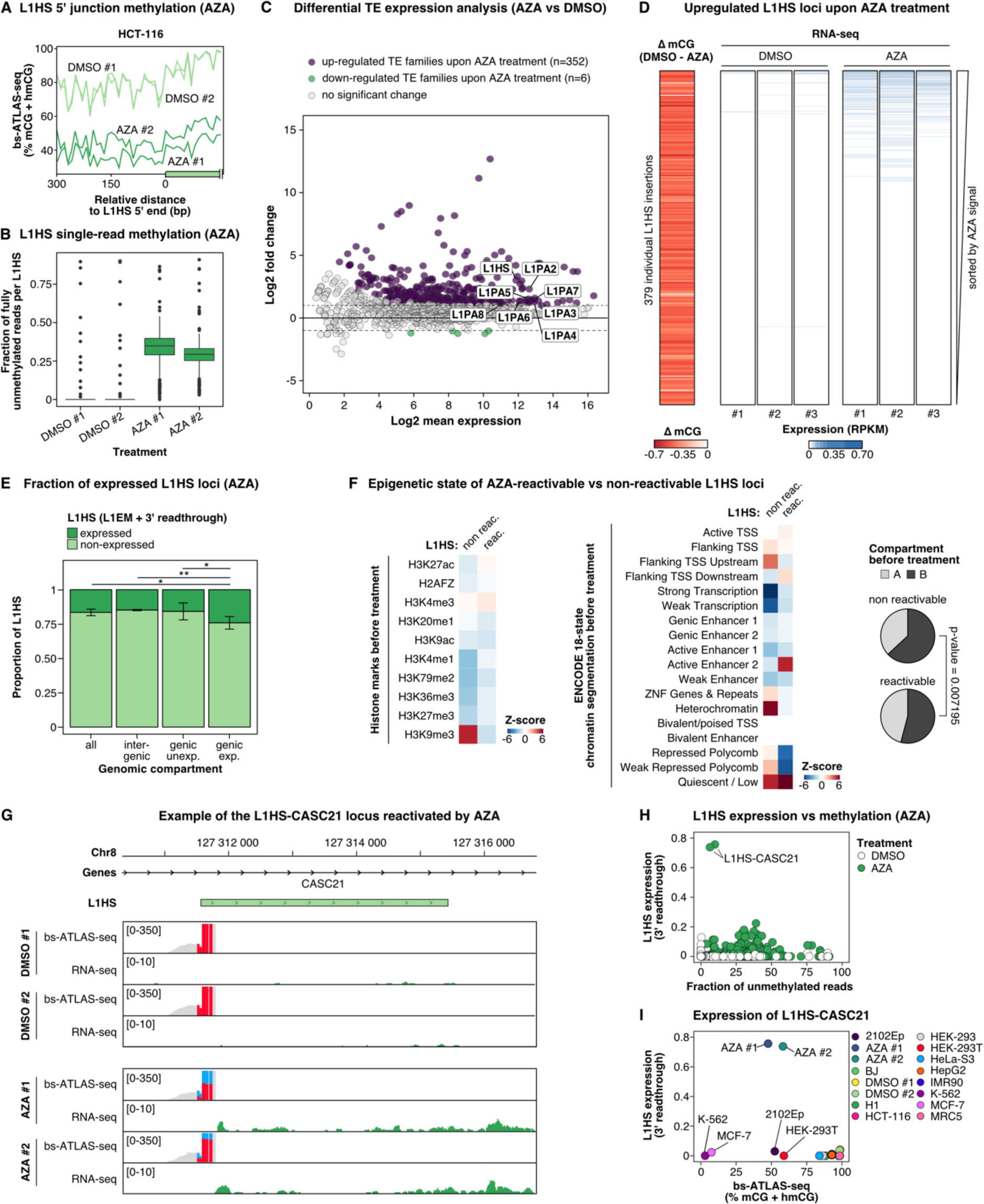
Acute DNA demethylation only reactivates the expression of a minor fraction of L1HS loci. **(A)** Metaplot of L1 DNA methylation profiles (bs-ATLAS-seq) at the L1HS promoter and upstream flanking region (300 bp) in HCT-116 treated with 5-aza-2-deoxycytidine (two replicates, AZA #1 and AZA #2; dark green) or with DMSO as negative control (two replicates: DMSO #1 and DMSO #2; light green). **(B)** Fraction of fully unmethylated reads in 5-aza- or DMSO-treated HCT-116 cells. Boxplots represent the median and interquartile range (IQR) ± 1.5 * IQR (whiskers). Outliers beyond the end of the whiskers are plotted individually. See (C) for legend. **(C)** Differential expression of transposable element (TE) families between 5-aza- or DMSO-treated HCT-116 cells measured by poly(A)^+^ RNA-seq using TEtranscripts ^142^. In the MA-plot, each data point represents an aggregated TE family. TE families found significantly up- or down-regulated upon 5-aza treatment are colored in purple and green, respectively, and data points corresponding to the L1HS to L1PA8 families are labelled (of which all but L1PA8 are upregulated). **(D)** Heatmaps showing the average difference of L1HS methylation (ΔmCG, bs-ATLAS-seq, 2 replicates) between HCT-116 cells treated by DMSO and 5-aza (AZA), as well as the expression levels of each L1HS in RNA-seq replicates (L1 3’ readthrough, see legend of **Figure 6A**). Heatmaps were sorted by decreasing L1 expression (average of the 3 replicates). **(E)** Barplot indicating the proportion of unexpressed (light green) and expressed (dark green) L1HS loci under 5’-aza treatment according to their genic environment and activity in HCT-116 cells. *p < 0.05, **p < 0.01, two-sided two-proportions z-test. **(F)** Association of L1HS elements that can be reactivated or not by 5-aza treatment, with pre-treatment histone modifications (left), chromatin states (18-state chromHMM, middle), or A/B-compartments (right) in HCT-116 cells. Heatmaps display the magnitudes of overlaps between L1 flanking sequences (±100 bp) and the genomic feature of interest expressed as Z scores (blue for depletion and red for enrichment). Pie charts represent the proportion of L1HS elements in the A or B-type of compartment, for reactivable vs non-reactivable copies (p-value, two-sided two-proportions z-test). Chromatin segmentation and genome compartmentalization data for HCT-116 cells were obtained from ENCODE and ^143^, respectively. Reactivated loci correspond to the 85 loci detected as expressed under 5’-aza treatment in HCT-116 by the two methods (union of 3’ readthrough and L1EM). **(G)** Genome browser view of the L1HS-CASC21 locus treated (AZA) or not (DMSO) by 5-aza, integrating L1 methylation (bs-ATLAS-seq) and expression (poly(A)+ RNA-seq). **(H)** L1HS expression vs methylation in 5-aza-treated HCT-116 cells. L1 methylation is defined here as the mean fraction of fully unmethylated reads per L1HS locus and per condition (AZA or DMSO) and expression is estimated through L1 3’ readthrough as described in the legend of **Figure 6A**. Each point represents an L1HS locus and a replicate and was colored in light (DMSO) or dark (AZA) green. **(I)** Comparison of the expression and methylation levels of an intronic L1HS insertion located in the CASC21 gene, across cell types and conditions. This element is only expressed upon 5-aza treatment, but not in other cell types with similar or even completely abolished methylation (MCF-7, K-562, 2102Ep or HEK-293T).

To understand what differentiates reactivable loci from those that remain repressed, we measured the association of each group of loci with histone modifications and chromatin segmentation states obtained from the ENCODE project ^108, 109^. Non-reactivable L1HS loci are associated with H3K9me3-rich regions (**Figure 7F**, left), as well as with heterochromatin (**Figure 7F**, middle). Consistently non-reactivable elements are enriched in B-compartments as compared to reactivable ones (**Figure 7F**, right). These observations suggest that multiple layers of epigenetic repression coexist in the same cell type, on the same family, and even on the same locus, and may persist at the majority of loci after acute DNA demethylation by 5-aza, at least in HCT-116 colon carcinoma cells. Interestingly, an L1HS element inserted in the intron of *CASC21* (L1-CASC21) shows strong levels of expression after 5-aza treatment in HCT-116 (**Figure 7G** and **Figure 7H**). In K-562 and in MCF-7, this element is also unmethylated but not expressed (**Figure 7I**) supporting the idea that cell-type specific factors are necessary to activate L1HS expression, or that alternative epigenetic pathways may supplant DNA methylation in the latter cell lines.

## DISCUSSION

Understanding the impact and regulation of L1 elements in humans requires genome-wide strategies able to profile the DNA methylation of full-length elements. Those belonging to the human-specific L1 (L1HS) family are especially relevant, as they form the only family capable of autonomous retrotransposition in our genome, but other young primate-specific L1 elements also constitute an important source of genetic innovation. These families are notoriously difficult to study as all copies are almost identical to each other - an issue accentuated by the C-to-T conversion in bisulfite sequencing protocols - and because individual genomes significantly differ from the reference human genome with respect to the presence or absence of L1HS insertions ^34, 68^. To overcome these bottlenecks, we have developed bs-ATLAS-seq, which gives access to the position and methylation state of L1HS, including non-reference insertions, as well as those of many L1PA elements, at single-locus, single-nucleotide and single-molecule resolutions (**Figure 1**). Bs-ATLAS-seq offers specific advantages, including excellent cost-effectiveness, since it requires only 10-20 million reads per sample, and versatility with regard to genomic DNA quality, since it works with partially fragmented genomic DNA, as is typically the case in clinical samples. It can be used in conjunction with other emerging approaches based on nanopore long-read sequencing, which for their part can haplotype-resolve DNA methylation over the entire locus ^89, 110, 111^, as illustrated by our allele-specific methylation analysis (**Figure 4**).

We comprehensively located young full-length L1 elements and characterized their DNA methylation in a panel of twelve normal, embryonal, or cancerous cell lines, providing one of the most detailed catalogues of L1 DNA methylation in human cells so far (https://L1methdb.ircan.org). Most cells studied here belong to ENCODE top-tier cell lines, enabling integration with a wide variety of publicly available functional genomics data, and thus facilitating the exploration of retrotransposon-host genome interactions. We observed that in most cell types but embryonic cells, the methylation of intragenic L1s is largely influenced by gene expression (**Figure 3**). Global L1 DNA methylation has been extensively used as a cancer biomarker, often as a surrogate for measuring global genome methylation levels ^112^. Our findings suggest that deconvoluting L1 methylation signal at the level of individual loci, particularly those inserted in genes, may represent an alternative source of DNA-based biomarkers capturing cell-type-specific gene expression.

Early observations at select loci showed that exogenous retroviruses and transposable element insertions could be targeted by DNA methylation, which then spreads to neighboring regions ^113–116^. However, a more systematic search of mouse endogenous retroviruses (ERVs) capable of spreading DNA methylation to nearby gene promoters has detected only a single example ^117, 118^. By contrast, hundreds of transposable element loci may affect the methylation status of their adjacent sequence in *Arabidopsis thaliana* ^119^. In most cases, DNA methylation propagates over a few hundred base pairs, but can reach several kilobases in *A. thaliana* ^119^. In addition, hypomethylated CpG islands can induce so-called methylation “sloping shores” in their upstream sequence, a property previously reported for a GFP reporter gene containing a CpG island and mobilized by an engineered mouse retrotransposon ^87^, and for an active progenitor L1 element ^63^. The extent and significance of these phenomena remained uncertain for human L1 elements. In our survey, we found that approximately one third of informative loci exhibit spreading of DNA methylation from a methylated L1 to the adjacent sequence, while another third shows demethylation of the flanking region upstream of hypomethylated L1s (**Figure 4**). We note that, in theory, allele-specific alterations of DNA methylation associated with L1 insertions could reflect L1 insertions into pre-existing epialleles. However, in most cases, methylation follows a gradient starting from the L1 element, suggesting that the insertion is directly responsible for the proximal epivariation in the flanking sequence, and not the contrary. Finally, we found that L1-mediated alterations of DNA methylation do not extend beyond 300 bp. This is contrasting with long-distance DNA methylation propagation that can reach several kilobases in *A. thaliana*, and may involve plant-specific pathways such as RNA-directed DNA methylation ^119, 120^, or even longer distance effects driven by retrotransposon transcriptional activity ^121^. Although L1 affects nearby DNA methylation only at short distances, epivariation in the zone of influence is associated with differential binding of transcription factors and can affect the host transcriptome (**Figure 5**). These findings parallel recent observations in mice indicating that polymorphic ERVs and L1s can alter local chromatin accessibility_122._

One of our original questions was to test whether all methylated L1s were repressed and whether, reciprocally, all unmethylated L1s were expressed. We found that the majority of unmethylated L1s remains silent (**Figure 6**). Consistently, only a fraction of L1s appear reactivable upon acute DNA demethylation by a demethylating agent (**Figure 7**). We conclude that, for most loci, L1 hypomethylation alone is insufficient to induce its expression and that other mechanisms prevent L1 reactivation in the absence of DNA methylation. We uncovered two non-exclusive scenarios. First, L1 silencing pathways can function redundantly and cohabit with DNA methylation at individual loci. Consistently, we found that L1HS elements not reactivated upon 5-aza treatment are enriched in H3K9me3-bound heterochromatic regions and B-compartments *before* demethylation in the HCT-116 colon cancer cell line (**Figure 7F**). Deposition of this repressive mark could involve Setdb1 or other histone methyl transferases ^123–126^, depending on the cell type, and be tethered by KZFPs-TRIM28 ^127^ or the HUSH complex ^43–45^. In other cell types, repression could rather involve SIN3A and the local recruitment of histone deacetylases ^128^. Incidentally, we note that HUSH-mediated L1 silencing was first discovered through a CRISPR screen in K-562 cells ^43^, in which L1PA elements appear to be virtually devoid of DNA methylation (**Figure 2**). Knocking out any component of the HUSH complex in K-562 cells leads to a massive increase of L1 expression, but has more modest effects in other cell types ^43^. Second, the expression of cell-type-specific TFs binding within or nearby L1, such as ESR1, can be required to switch from an activable – but quiescent – state to an active state ^129^. Accordingly, knocking down ESR1 expression limits L1 expression in the breast cancer cell line MCF-7 (**Figure 5**).

Retrotransposons and their chimeric transcripts represent a rich source of tumor-specific antigens, which can be specifically recognized by infiltrating T cells or targeted by CAR-T cells ^130–133^. Furthermore, they have the ability to induce a viral mimicry state, which stimulates innate immunity through the sensing of double-stranded RNA and cytosolic DNA species ^45, 134–138^. By modulating retrotransposon expression, epigenetic drugs can thus enhance both specific and innate antitumoral immune responses. Our findings underscore the need to precisely delineate the specific pathways controlling the expression of individual loci. This knowledge will be critical for the development of rational drug combinations capable of specifically re-expressing L1 elements and L1-derived tumor-specific antigens of interest for immunotherapy, while minimizing off-targets.

### Limitations of the study

In our study, we identified a set of TFs associated with distinct L1HS or L1PA2 methylation states. Given the high degree of identity between individual copies of these young L1 families and the resolution of ChIP-seq experiments, only TFs binding sufficiently close to the 5’ end of the element can be unambiguously assigned to individual elements. Therefore, TFs that potentially bind L1s more internally might be missed, even if documented in Unibind. As approaches capable of mapping TF binding sites within repeated sequences, such as PAtChER ^74^, DiMeLo-seq ^139^, nanopore-DamID ^140^, or nanopore-based NOMe-seq ^141^, become more widely available, this problem will become less acute in the future. Additionally, we present data showing association between L1 methylation state and other genomic features. In some cases, we could confidently infer causality (L1-mediated epivariation of the proximal genomic region, transcription-mediated methylation of intronic L1s, ESR1-driven L1 chimeric transcript synthesis). However, in other cases this remains an experimental challenge. A preponderant difficulty is that L1 methylation could have been established in a different cellular context through factors not necessarily present anymore, and this methylation pattern could have been maintained since then.

## STAR*METHODS

Detailed methods are provided in the online version of this paper and include the following:

- Key resources table
- Resource availability

`o` Lead contact
`o` Materials availability
`o` Data and code availability
- Experimental model and subject details
- Method details

`o` Bs-ATLAS-seq
`o` Cas9-targeted nanopore sequencing
`o` Methylated DNA immunoprecipitation (MeDIP)
`o` LUMA
`o` Infinium Human Methylation 450K BeadChip analysis
`o` RNA-seq
`o` 5-aza-deoxycitidine treatment
`o` ChIP-seq
`o` Transcription factor enrichment at unmethylated L1
- Quantification and statistical analysis

## ACKNOWLEDGMENTS

This research was funded, in whole or in part, by the Agence Nationale de la Recherche, Grant ANR-21-CE12-0001. A CC-BY 4.0 public copyright license has been applied by the authors to the present document and will be applied to all subsequent versions up to the Author Accepted Manuscript arising from this submission, in accordance with the grant’s open access conditions.

We thank C. Baudoin and the IRCAN Genomics Core Facility (GenoMed) for sequencing, O. Croce and IRCAN bioinformatics service for computing resources, B. Meyer for deploying the web portal on IRCAN server, and members of the Cristofari lab for helpful discussions. We are grateful to the ENCODE Consortium for RNA-seq and ChIP-seq data, and to J-L Garcia-Perez (Univ. of Edinburgh, UK) for providing H1 genomic DNA. This work was supported by joint grants awarded to PAD and GC, and to DvE and GC from the Agence Nationale de la Recherche (RETROMET, ANR-16-CE12-0020; ActiveLINE, ANR-21-CE12-0001), as well as other grants awarded to GC from the Fondation pour la Recherche Médicale (FRM, DEQ20180339170), the Agence Nationale de la Recherche (Idex UCAJEDI, ANR-15-IDEX-01; Labex SIGNALIFE, ANR-11-LABX-0028; ImpacTE, ANR-19-CE12-0032), the Institut National du Cancer (INCa PLBIO 2020-095), the Canceropôle Provence-Alpes-Côte d’Azur, INCa and the Provence-Alpes-Côte d’Azur Region (Projet Emergence), Inserm (GOLD cross-cutting programme on genomic variability), CNRS (GDR 3546). Equipment acquisition for the GenoMed facility was supported by FEDER, Région Provence Alpes Côte d’Azur, Conseil Départemental 06, ITMO Cancer Aviesan (plan cancer) and Inserm. Work in the laboratory of PAD was also supported by LabEx “Who Am I?” (ANR-11-LABX-0071), Université Paris Cité IdEx (ANR-18-IDEX-0001) funded by the French Government through its “Investments for the Future” program, Fondation pour la Recherche Médicale, and Fondation ARC (Programme Labellisé PGA1/RF20180206807).

## AUTHOR CONTRIBUTIONS

GC and PAD conceived the study and secured funding. CP developed the bs-ATLAS-seq procedure. SL and GC designed and conducted the computational analyses. DP developed the web interface to interrogate the data. CP, AS, CD, LF, AJD, DvE and SS contributed to other experiments. SL, PAD and GC wrote the manuscript with inputs from all other authors. GC supervised the project.

## DECLARATION OF INTERESTS

GC is an unpaid associate editor of the journal *Mobile DNA* (Springer-Nature).

---

**Figure S1 related to Figure 1.**
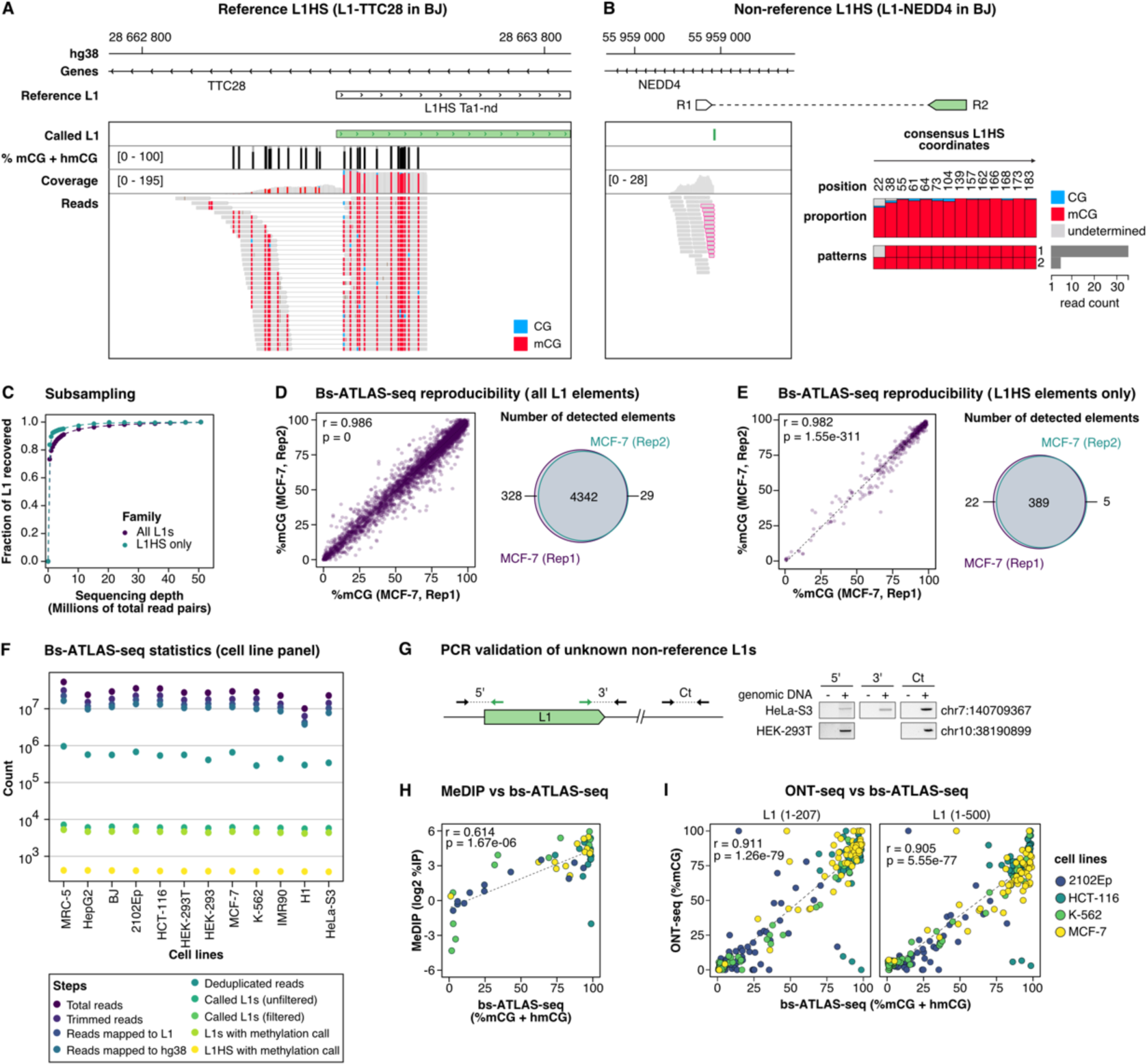
Optimization, sequencing statistics and validation of bs-ATLAS-seq. (**A, B**) Genome browser view of bs-ATLAS-seq results in BJ foreskin fibroblasts at the reference L1HS-TTC28 locus (**A**) and at the non-reference L1HS-NEDD4 locus (**B**). In the track showing the percentage of methylation, called CpG are indicated by a vertical gray bar, and the percentage of methylation as an overlapping black bar. In the ‘coverage’ and ‘reads’ tracks, vertical colored bars correspond to non-methylated CpG (blue) and methylated CpG (mCG, red). For non-reference L1HS, only the genomic flank covered by read #1 (R1; bottom left) is visible in the genome browser view. Soft-clipped reads supporting the 5’ L1 junction (split reads) are framed in pink. The proportion of mCG at each site and the frequency of the most common methylation patterns deduced from read #2 (R2; bottom right) are indicated on the charts (right). Positions of CpGs are related to L1HS consensus sequence (see Methods). (**C**) Sensitivity analysis. Computational down-sampling of high depth bs-ATLAS-seq sequencing data (MCF7) shows that L1 recovery reaches a plateau above 10 million of total read pairs. Thus, all samples were subsequently sequenced to a depth greater than 10 million of total reads. (**D, E**) Reproducibility of bs-ATLAS-seq. Two independent libraries of MCF-7 (from two subsequent MCF-7 passages) and sequencing runs are compared with respect to L1 elements of all families (**E**) or to L1HS elements only (**F**). Replicate 1 (MCF7, Rep1) was down sampled to the sequencing depth of replicate 2 (MCF7, Rep2) for comparison purpose. Left panels, correlation of methylation levels for shared detected L1 loci. Right panels, Venn diagram showing the overlap of detected L1 loci between the two libraries. **(F)** Statistics of bs-ATLAS-seq for the 12-cell line panel. See also **Table S1**. **(G)** PCR validation of unknown non-reference insertions. PCR was done with genomic DNA of the indicated cell lines (+) or with water as non-template control (-). The insertion in chr10 is pericentromeric and embedded in other repeats. Therefore, only primers to amplify the 5’ junction could be designed. Ct, unrelated locus used as PCR control. **(H)** DNA methylation level of selected L1 elements was profiled using an antibody-based enrichment of DNA methylation (MeDIP) and compared with bs-ATLAS-seq data. The DNA methylation level of MeDIP data is expressed as log2 of the percentage of immunoprecipitation (log2 %IP). See also **Table S2**. **(I)** Comparison of bs-ATLAS-seq with PCR-free targeted sequencing and methylation calling by Oxford Nanopore Technology sequencing (ONT-seq) for the L1 region common to both methods (1-207) or the full CpG island (1-500). See also **Table S4**. For all correlation analysis, r and p represent Pearson correlation coefficient and p-value, respectively.

**Figure S2 related to Figure 2.**
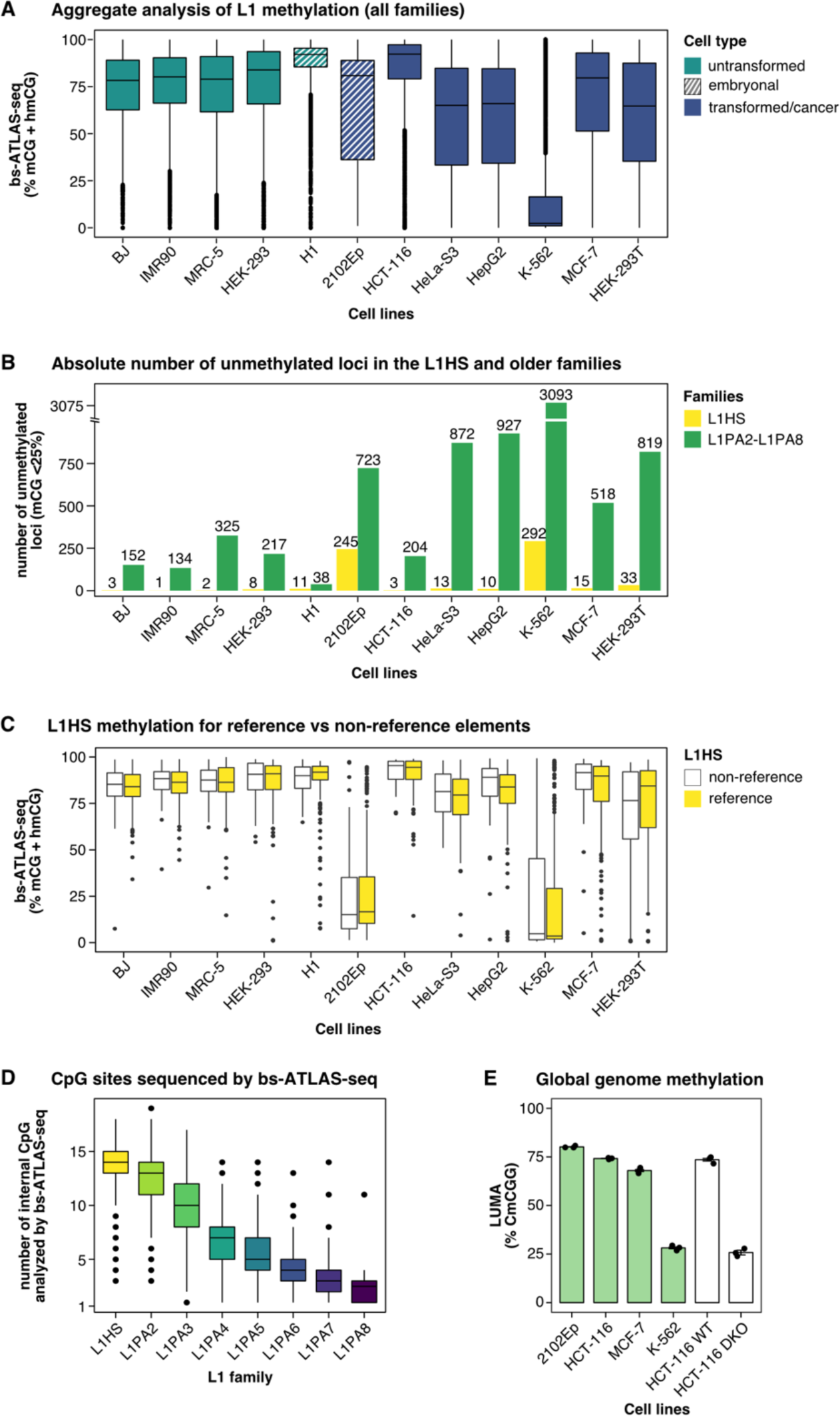
DNA methylation at different scales. **(A)** Aggregate DNA methylation level of the L1 promoter for all L1 elements detected by bs-ATLAS-seq (L1HS to L1PA8) across normal (green), embryonal (hatched), or transformed and cancer (blue) cells. **(B)** Barplot indicating the absolute numbers of unmethylated L1 copies (% mCG<25% according to bs-ATLAS-seq) across the different cell lines, for L1HS (light green) and older copies (L1PA2 to L1PA8, dark green). **(C)** Comparison of methylation levels for reference and non-reference L1HS elements across the different cell lines. Differences are not significant (two-sided two-sample Wilcoxon test). **(D)** Number of CpGs per element analyzed by bs-ATLAS-seq for each L1 family. **(E)** Genome-wide global CpG methylation measured by LUminometric Methylation Assay (LUMA). A subset of the cell line panel assayed by bs-ATLAS-seq showing low- or high-levels of L1HS methylation were tested for genome-wide global CpG methylation by LUMA (green). HCT-116 DKO and WT refers to a double *DNMT1* and *DNMT3B* knock-out, and its parental cell line, respectively (white). These additional cell lines were used as controls in the LUMA assay. Bars represent the average percentage of methylated CCGG sites (mean ± sem, n=3 technical replicates). In panels (**A**), (**C**) and (**D**), boxplots represent the median and interquartile range (IQR) ± 1.5 * IQR (whiskers). Outliers beyond the end of the whiskers are plotted individually.

**Figure S3 related to Figure 3.**
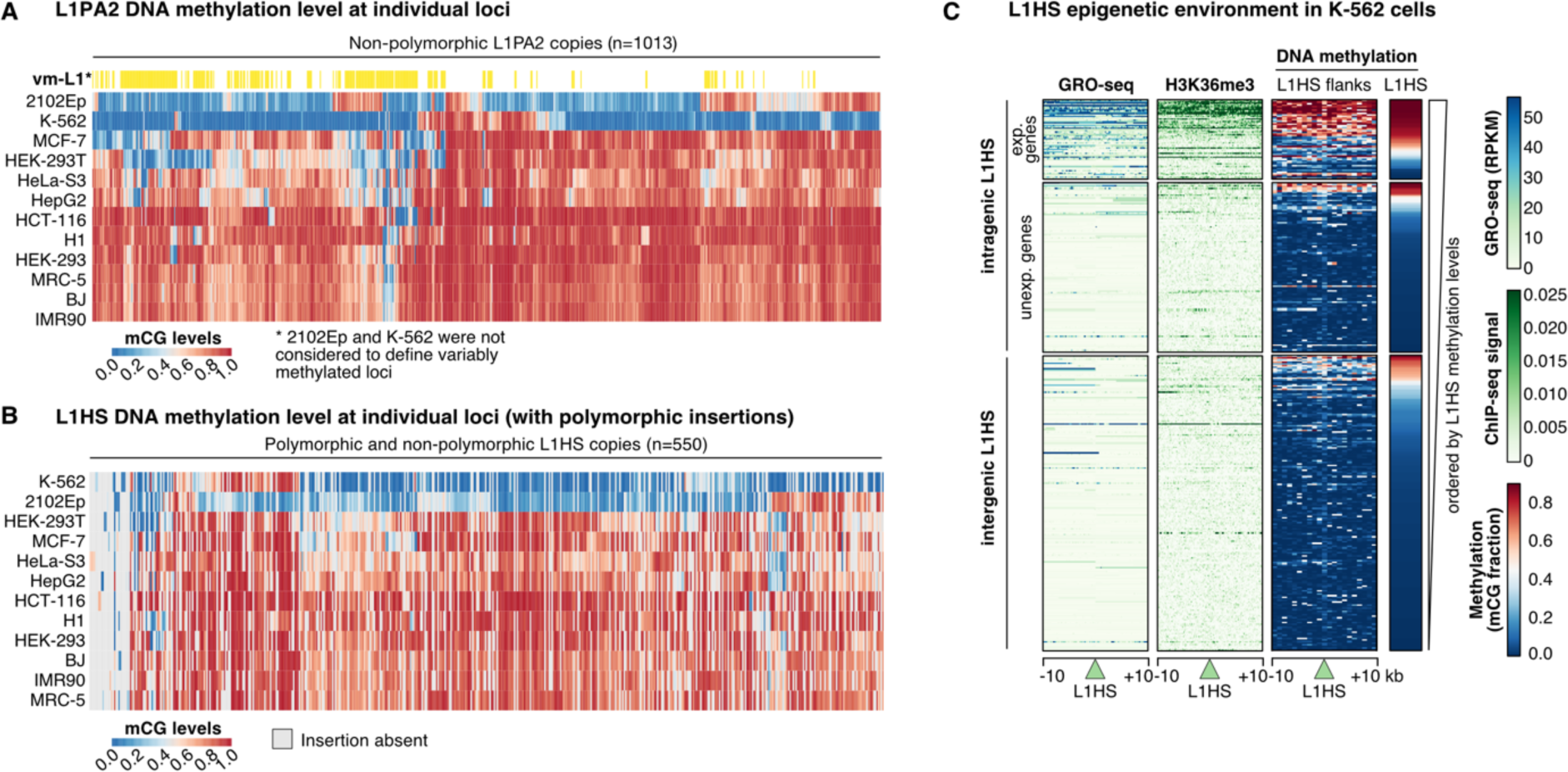
Methylation level of the youngest L1 subfamilies. **(A)** Heatmap of bs-ATLAS-seq methylation levels (% mCG + hmCG) for individual L1PA2 loci across a panel of 12-cell lines. Vm-L1, variably-methylated L1 loci (see **Figure 3** for definition). **(B)** Heatmap of bs-ATLAS-seq methylation levels (% mCG + hmCG) displaying values for both reference and non-reference L1HS across cell lines. When an insertion is absent in a given cell line, the heatmap cell is colored in grey. **(C)** Heatmaps illustrating nascent transcription (GRO-seq), H3K36me3 histone modifications (ChIP-seq), and DNA methylation (whole genome bisulfite sequencing, WGBS), in 10 kb-windows upstream and downstream L1HS elements (green triangle). Loci are separated according to their position relative to genes (left), separating expressed and unexpressed genes, and sorted by decreasing L1 methylation levels (bs-ATLAS-seq, right). A similar heatmap including all families (from L1HS to L1PA8) is shown in **Figure 3F**.

**Figure S4 related to Figure 4.**
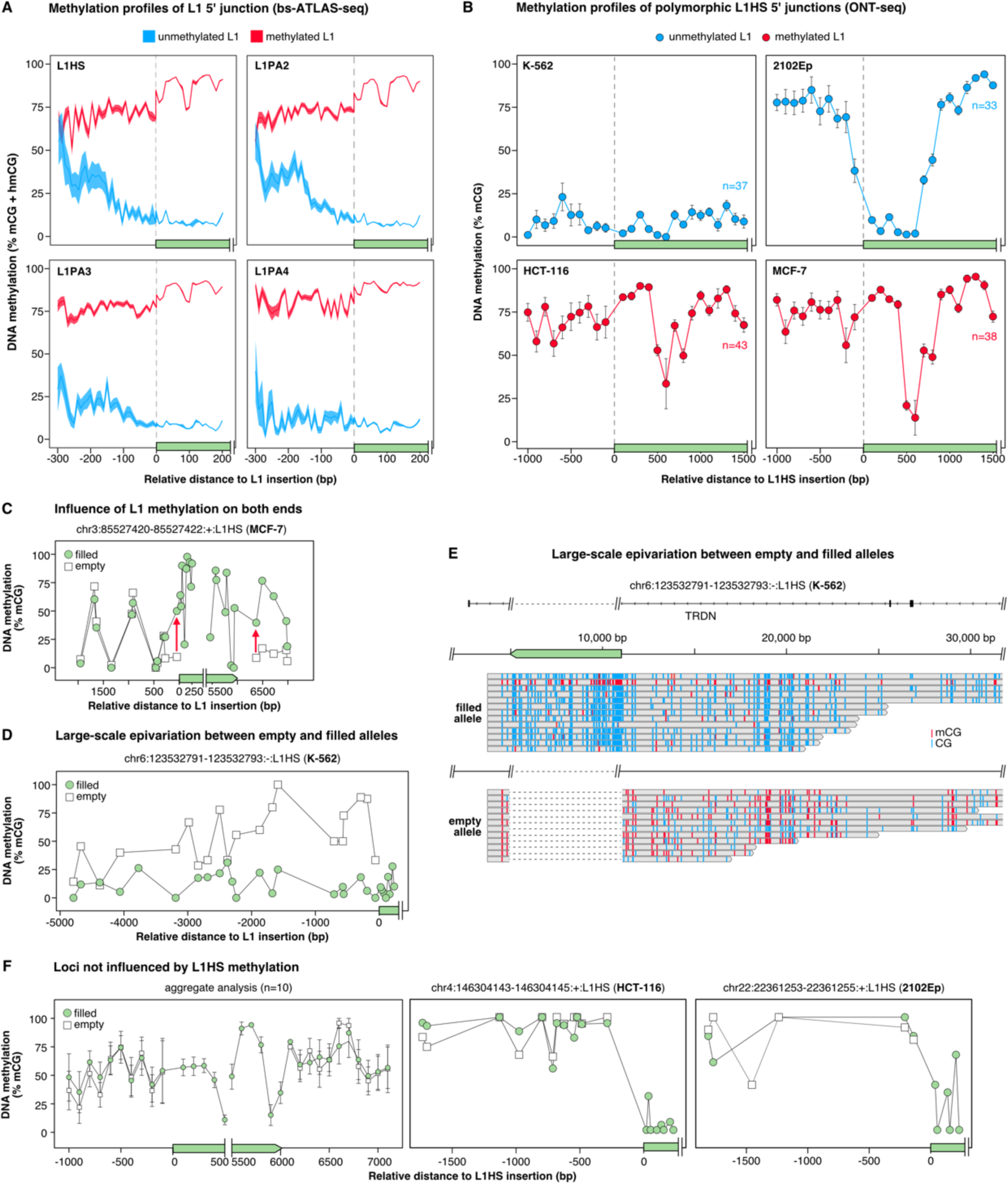
DNA methylation profiles of L1s and their target loci. **(A)** DNA methylation profiles of methylated (mCG ≥ 75%, red) and unmethylated (mCG ≤ 25%, blue) L1 (L1HS to L1PA4) 5’ junctions obtained by bs-ATLAS-seq data. Data represent average DNA methylation levels in 10 bp-bins aggregated from the 12 cell lines (mean ± 95% C.I.). **(B)** DNA methylation profiles of methylated (mCG ≥ 75%; red) and unmethylated (mCG ≤ 25%; blue) L1HS 5’ junctions obtained by ONT-seq in 4 different cell lines (K-562, 2102Ep, HCT-116, MCF-7). Note that for the sake of comparison, the distinction of methylated vs unmethylated L1 is based on the first 15 CpG, as for bs-ATLAS-seq. Given the small numbers, unmethylated L1 in HCT-116 and MCF-7 (n=3 and n=3, respectively), and methylated L1s in K-562 (n=4) were not plotted. Data points represent average mCG levels in 100 bp-bins for each cell line (mean ± s.d.). **(C)** Example of locus with DNA methylation spreading from L1 to the external flanks. Red arrows denote hypermethylation relative to the empty locus. **(D, E)** Large-scale allele-specific epivariation associated with an L1 insertion. (D) Methylation levels and (E) genome browser view (top: filled allele, bottom: empty allele). The L1 insertion is depicted as a green solid arrow. Methylated and unmethylated CpG are indicated in red and blue, respectively. **(F)** Loci not influenced by L1 methylation state. Left, average DNA methylation levels in 100 bp-bins (n=10, mean ± s.d.). Middle and right, examples of loci not influenced by L1 methylation. For (A) to (D), and (F), the x-axis represents the relative distance to L1 5’ end (green) and the y-axis the percentage of DNA methylation. For (C), (D) and (F), the empty (white squares) and filled (green circles) alleles are overlaid.

**Figure S5 related to Figure 5.**
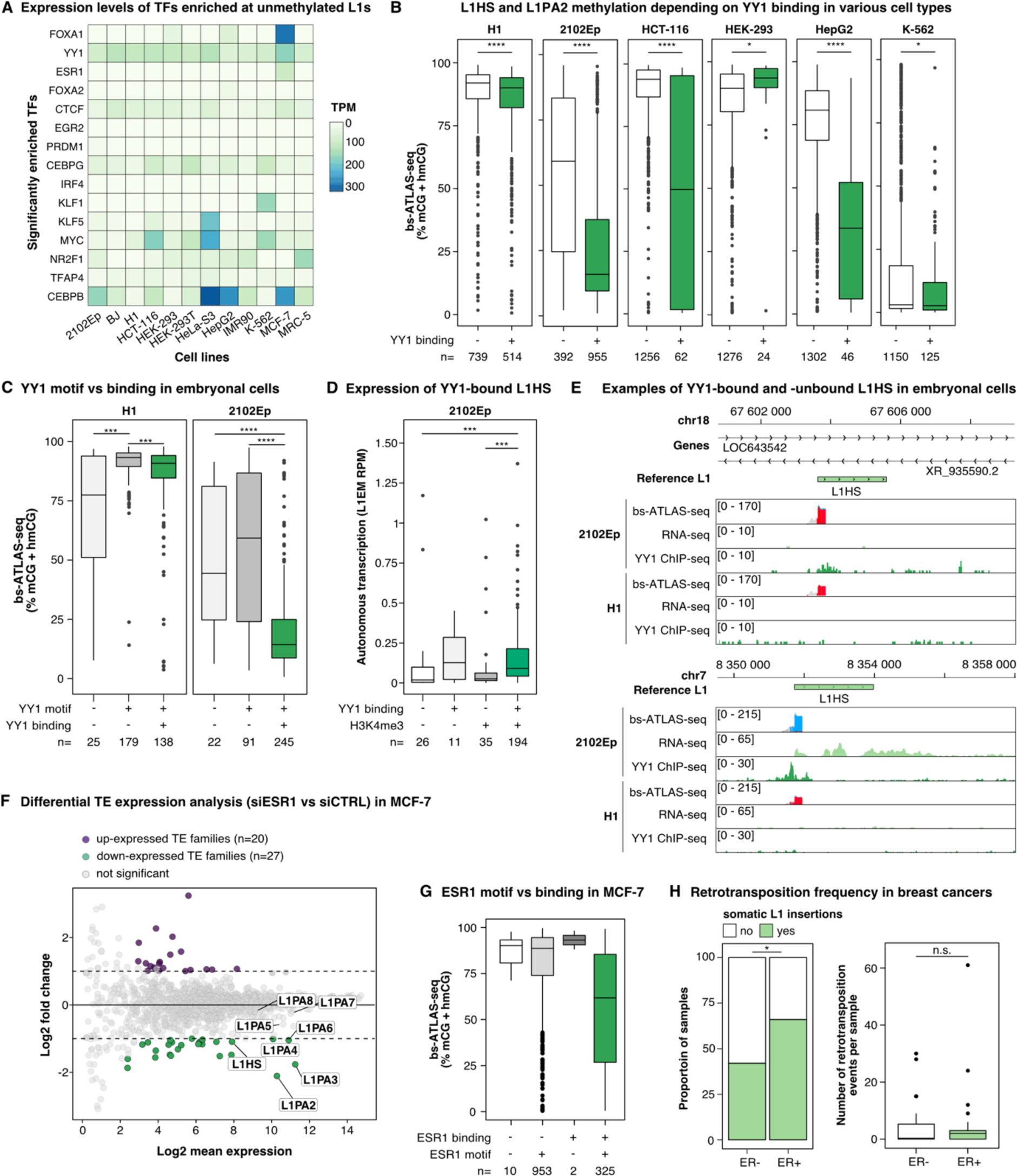
Unmethylated L1HS and L1PA2 are bound by YY1 in embryonal cells. **(A)** Heatmap showing the expression levels of the 15 most enriched TF identified in the screen across the panel cell lines. FOXA1 and ESR1 are more expressed in MCF-7 as compared to other cell types whereas YY1 is more ubiquitously expressed, even if it predominantly binds to L1 elements in embryonal cells (H1 and 2102Ep) (see **Figure 5E** and panel B). Expression level is measured as transcripts per million (TPM). **(B)** DNA methylation levels of L1HS and L1PA2 elements bound (+) or unbound (-) by YY1 in embryonal cell lines (H1 and 2102Ep) and other cell lines for which matched YY1 ChIP-seq were also publicly available (K-562, HCT116, HepG2, HEK-293T). The number of L1HS copies in each subset (n) is indicated at the bottom of the plot. **(C)** DNA methylation levels of L1HS loci with (+) or without (-) YY1 binding motifs in their 5’ UTR, and actually bound (+) or not (-) by YY1 in H1 and 2102Ep cells. **(D)** Expression level of L1HS element bound (+) or not (-) by YY1 and associated (+) or not (-) with H3K4me3 histone modification in 2102Ep cells. Locus-level expression was estimated by L1EM. The number of L1HS copies in each subset (n) is indicated at the bottom of the plot. **(E)** Genome browser view of two example L1HS loci with distinct promoter DNA methylation profiles (bs-ATLAS-seq), integrated with RNA-seq and YY1 ChIP-seq data in H1 and 2102Ep. Top, locus in chromosome 18, the YY1 signal is close to the background level, the L1HS element is hypermethylated and non-expressed. Both cell lines have similar profiles. Bottom, locus in chromosome 7, a strong YY1 peak is detected in 2102Ep cells, where the L1HS is completely unmethylated and robustly expressed. In contrast, in H1 cells, the same locus does not appear bound by YY1, is hypermethylated and non-expressed. **(F)** Differential expression of transposable element (TE) families between MCF-7 cells treated by an siRNA against ESR1 (+) or a control scrambled siRNA (-) measured by RNA-seq using TEtranscripts ^142^ (data from GSE153250). In the MA-plot, each data point represents an aggregated TE family. TE families found significantly up- or down-regulated upon ESR1 knockdown are colored in purple and green, respectively, and data points corresponding to the L1HS to L1PA8 families are labelled (of which L1HS to L1PA6 are downregulated). **(G)** DNA methylation levels of L1HS and L1PA2 loci with (+) or without (-) ESR1 binding motif in their 5’ UTR and actually bound (+) or not (-) by ESR1 in MCF-7 cells. Note that the two loci without internal ESR1 binding motif but bound by ESR1 (dark grey) have a binding motif upstream (<300 bp) of the element. **(H)** Somatic L1 retrotransposition in breast cancer according to the estrogen receptor (ER) status in PCAWG samples. Left, proportion of cancer samples with at least one somatic L1 insertion. Right, number of somatic L1 retrotransposition events per sample. ER status was obtained from ^100^ and somatic L1 retrotransposition events were identified in ^48^

## STAR*METHODS

### KEY RESOURCES TABLE

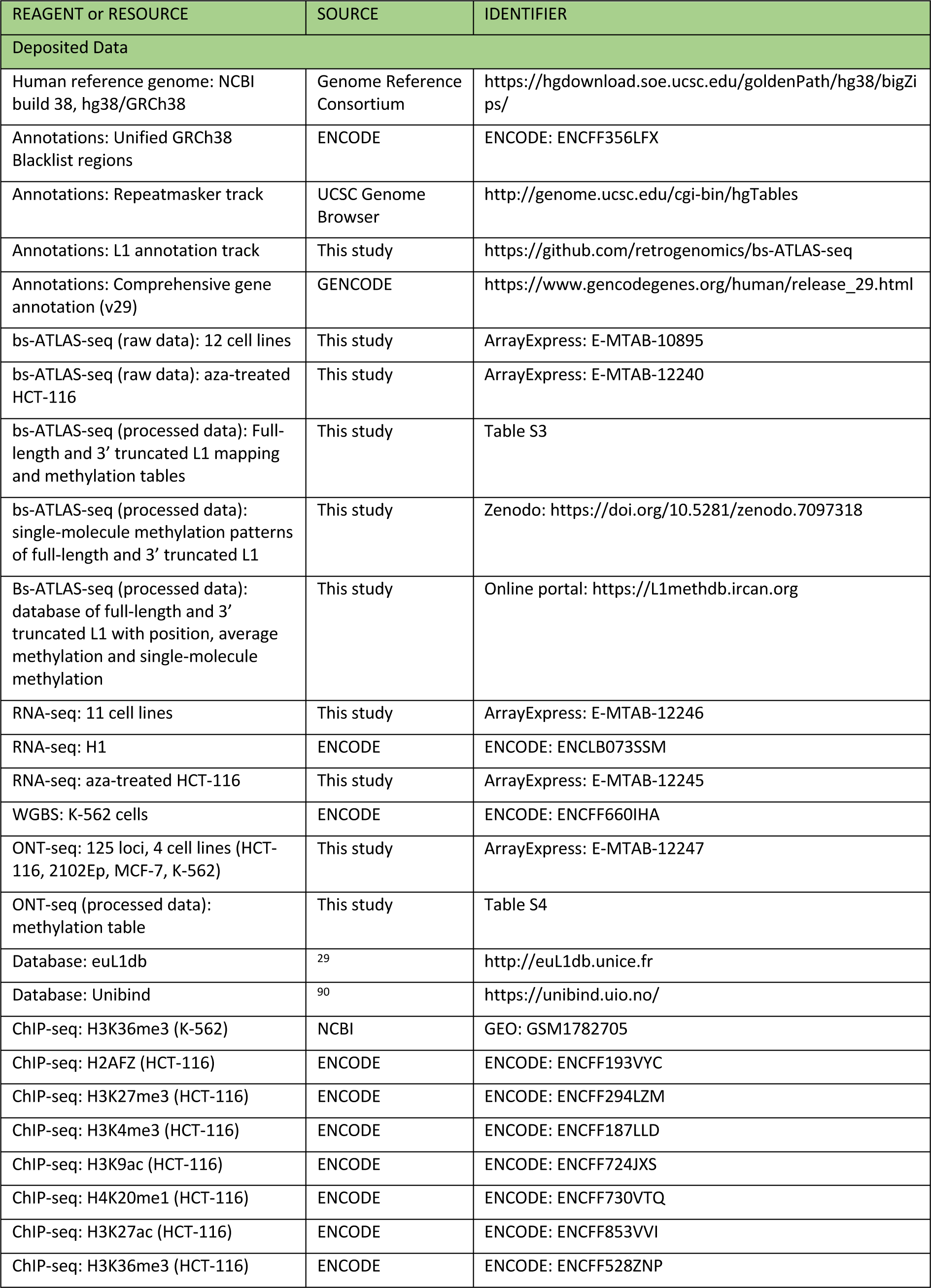
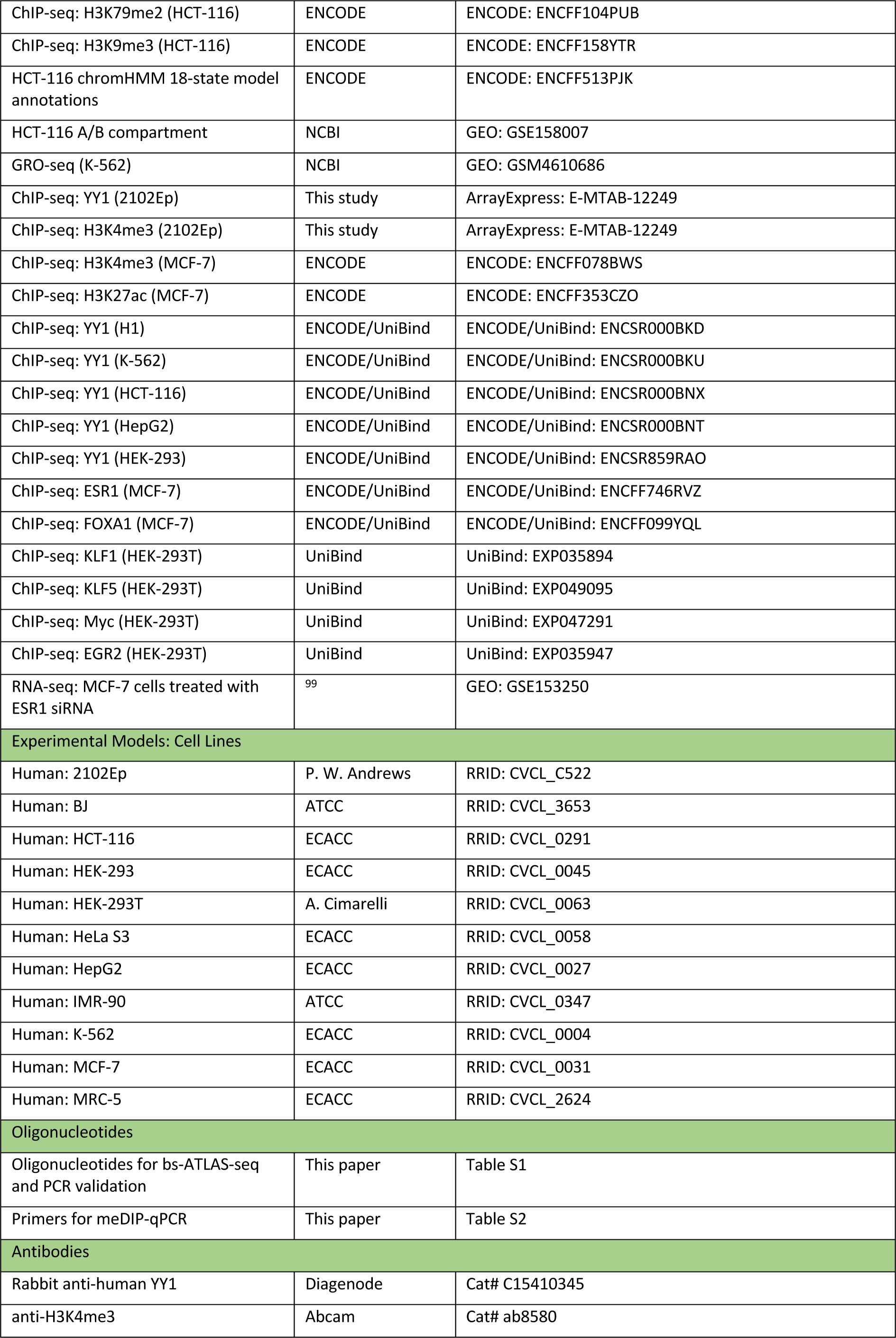
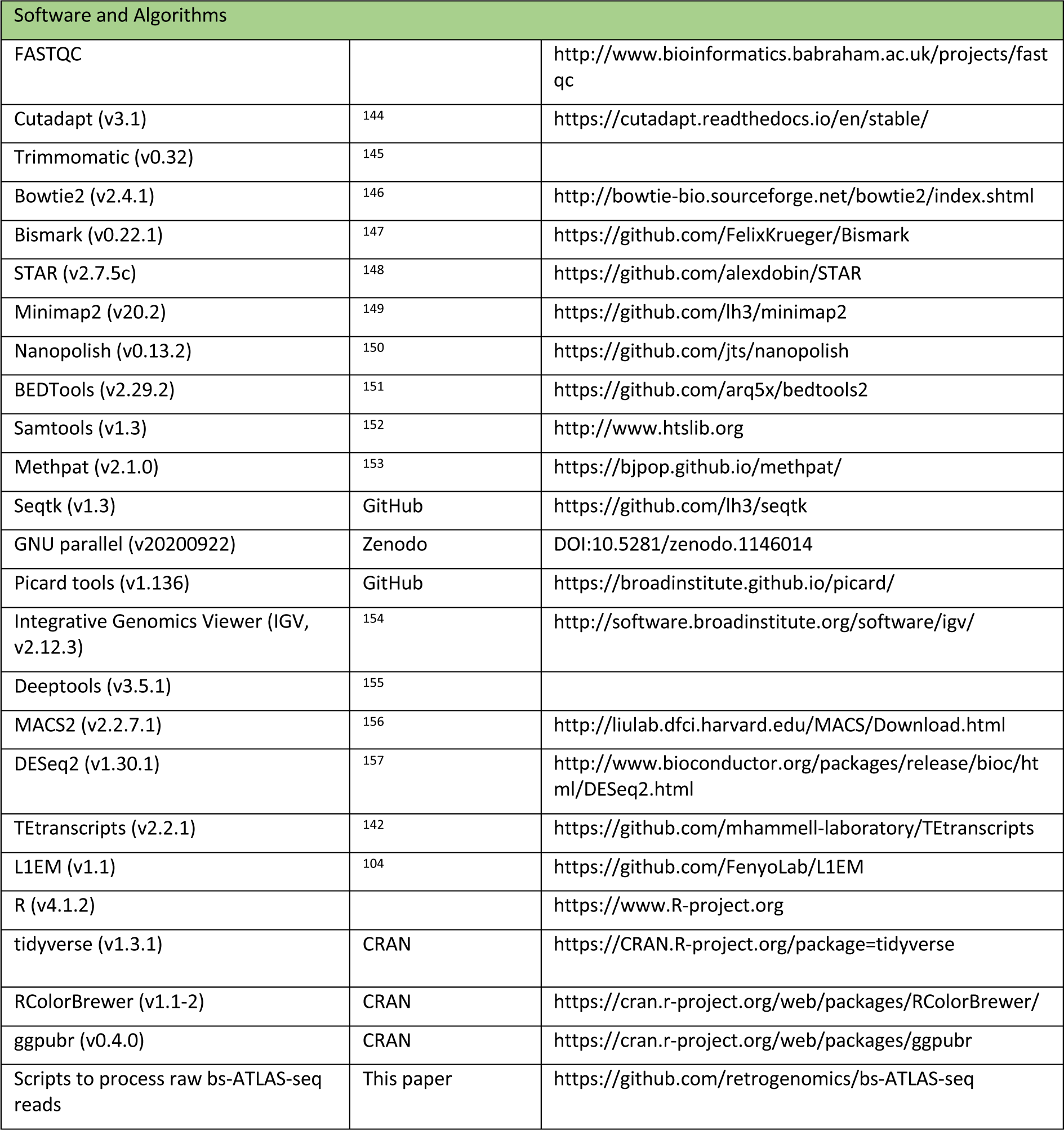

## RESOURCE AVAILABILITY

### Lead contact

Further information and requests for reagents may be directed to, and will be fulfilled by, the corresponding author, Gael Cristofari.

### Materials availability

This study did not generate any new unique reagents or materials to report.

### Data and code availability

Data were submitted to the ArrayExpress database (www.ebi.ac.uk/arrayexpress) under accession numbers E-MTAB-10895 and E-MTAB-12240 (bs-ATLAS-seq), E-MTAB-12247 (ONT sequencing), E-MTAB-12249 (ChIP-seq), and E-MTAB-12246 and E-MTAB-12245 (RNA-seq). The genomic location and methylation levels of L1 insertions are summarized in **Table S3**. Single-molecule methylation patterns for each locus are provided in Zenodo (https://doi.org/10.5281/zenodo.7097319). All L1 methylation datasets can be interactively queried, filtered and downloaded through the web portal L1MethDB (https://L1methdb.ircan.org/). The scripts written to call L1 insertions and CpG methylation from bs-ATLAS-seq data, as well as useful annotation files used in the course of this study, are available at https://github.com/retrogenomics/bs-ATLAS-seq.

## EXPERIMENTAL MODEL AND SUBJECT DETAILS

The cell lines used in this study are identical to those previously characterized in ^9^ and include primary fibroblasts (BJ, IMR90, MRC5), embryonic stem cells (H1) and cancer or transformed cell lines (HCT-116, K-562, HEK-293, HEK-293T, HeLa S3, MCF-7, HepG2, and 2102Ep). All cells were directly obtained either from ECACC (distributed by Sigma) or from ATCC (distributed by LGC Standards), apart from 2102Ep cells (a kind gift of P. W. Andrews, University of Sheffield, UK) and HEK-293T (a kind gift of Andrea Cimarelli, ENS-Lyon, France). H1 human embryonic stem cells were not grown in the laboratory for regulatory reasons but genomic DNA of H1 cells grown in presence of LIF and serum was a kind gift of J. L. Garcia-Perez (University of Granada, Spain). Cells were maintained in a tissue culture incubator at 37°C with 5% CO_2_ and grown in Dulbecco’s modified Eagle medium (DMEM), McCoy’s 5A (HCT-116) or RPMI 1640 (K-562) containing 4.5 g/L D-Glucose, 110 mg/L Sodium Pyruvate, and supplemented with 10% FBS, 862 mg/mL L-Alanyl-L-Glutamine (Glutamax), 100 U/mL penicillin, and 100 µg/mL streptomycin. Cell cultures tested negative for mycoplasma infection using the MycoAlert Mycoplasma Detection Kit (Lonza). Cell line authenticity was verified by multiplex STR analysis (Eurofins) and comparison with the DSMZ database (https://celldive.dsmz.de/) or with previously published profiles for H1 and 2102Ep cells ^158, 159^.

## METHOD DETAILS

### Bs-ATLAS-seq

A practical protocol for bs-ATLAS-seq is provided in ^160^ and his detailed below.

### DNA extraction

Genomic DNA was prepared with the QiaAmp DNA Blood mini kit (Qiagen) and quantified by fluorometry using the Quant-iT dsDNA HS Assay and a Qubit fluorometer (Thermo Fisher Scientific).

#### Mechanical fragmentation, end-repair and A-tailing

Two micrograms of genomic DNA were sonicated for 6 cycles (6 s on, 90 s off) at 4 °C with a Bioruptor NGS (Diagenode), generating average fragments of 1 kb. Fragment size was controlled by capillary electrophoresis with the DNA high sensitivity kit and a Bioanalyzer 2100 (Agilent Technologies). DNA ends were repaired using the End-It DNA End-Repair Kit (Epicentre), and A-tailed with Klenow Fragment (3’-to-5’ exo-, New Englands Biolabs) following manufacturer’s protocol. At each step, DNA was purified with Agencourt AMPure XP beads (Beckman Coulter) using a 1:1 ratio of beads to DNA solution (v/v).

#### Linker ligation

Oligonucleotides LOU2493 (with all C methylated) and LOU2494 (**Table S1**) were mixed in 5 μL of 1x T4 DNA Ligase buffer (50 mM Tris-HCl pH 7.5, 10 mM MgCl_2_, 1 mM ATP, 10 mM Dithiothreitol; New England Biolabs) at a final concentration of 80 μM each and annealed by heating at 65 °C for 15 min, followed by slow cooling down to room temperature. Four hundred nanograms of fragmented genomic DNA were ligated with a 40-fold molar excess of the duplex linker overnight at 16 °C in 50 μL of 1x T4 DNA Ligase buffer supplemented with 400 U of T4 DNA Ligase (New England Biolabs). Excess linkers were removed by two successive rounds of purification with Agencourt AMPure XP beads using a 1:1 ratio of beads to DNA solution (v/v). Note that only the single-stranded methylated oligonucleotide LOU2493 is covalently bound to the 5’ ends of the genomic DNA fragments.

#### Bisulfite conversion

Two hundred and fifty nanograms of linker-ligated genomic DNA were subjected to sodium bisulfite conversion for 210 min at 64°C using the EZ DNA Methylation Kit (Zymo Research) according to the manufacturer’s instructions. After clean-up, converted DNA was kept at 4 °C for up to 20 h.

#### Suppression PCR

L1 5’ junctions were amplified in 40 µL-reactions containing 16 ng of converted and linker-ligated genomic DNA, 0.2 μM of primers, 0.2 μM dNTPs, 1.5 mM MgCl_2_, 0.8 U of Platinum Taq DNA Polymerase in 1X PCR buffer (Invitrogen). A first primer (LOU2565, or LOU2715 to LOU2724) targets the L1-specific region with a 5’ extension corresponding to Illumina Rd2 SP and P7 sequences, with a 10-nt index specific to the sample between them. A second primer (LOU2497) targets the linker (identical to Rd1 SP) and possesses a 5’ extension corresponding to Illumina P5 sequence. Primer sequences and annotations are provided in **Table S1**. Amplification was performed under the following cycling conditions: 1 cycle at 95°C for 4 min; followed by 20 cycles at 95°C for 30 s, 53°C for 30 s, and 68°C for 1 min; and a final extension step at 68°C for 7 min. To reduce PCR stochasticity, each sample was amplified in eight parallel 40 µL-reactions and subsequently pooled. In addition, another reaction was performed in the absence of the L1-specific primer to control for the absence of linker-to-linker amplification. The amplified library was cleaned-up from primers and irrelevant products by double-sided size-selection with Agencourt AMPure XP beads using a 0.55:0.65 ratio of beads to DNA solution (v/v), to reach an average library size of 450 bp. Finally, a last purification was achieved with Agencourt AMPure XP beads using a 1:1 ratio of beads to DNA solution (v/v) to eliminate potential remaining traces of oligonucleotides.

#### Sequencing

Libraries were quantified by qPCR with KAPA library quantification kit for Illumina (Roche) and their size range was checked by capillary electrophoresis using with the DNA high sensitivity kit and a Bioanalyzer 2100 (Agilent Technologies). Libraries were diluted to 1 nM and pooled equimolarly. Pooled libraries were paired-end sequenced with a NextSeq 550 system (Illumina) using a high-output kit and 300 cycles and 20% of PhiX DNA spike-in. To gain access to the methylation state of the first 15 CpG in L1 sequence, paired-end sequencing was performed asymmetrically with 90 cycles for read #1 and 210 cycles for read #2.

#### bs-ATLAS-seq primary analysis

Illumina paired-end sequencing reads were processed to locate L1 elements and to call their methylation status, using the script `bs-atlas-seq_calling.sh` (v 1.1, available at https://github.com/retrogenomics/bs-ATLAS-seq), which steps are summarized below. In each read pair, read #1 is 90 bp long and corresponds to the 5’ flanking sequence of L1, while read #2 is 210 bp-long and corresponds to L1 5’ UTR internal sequence.

#### Read trimming, mapping, and filtering

We demultiplexed FASTQ files according to their sample-specific barcode using cutadapt (v 3.1, https://github.com/marcelm/cutadapt). We then verified the presence of bs-ATLAS-seq adapters in the reads and trimmed them with cutadapt. Once trimmed, reads #2 were mapped locally against the first 250 bp of L1HS consensus sequence (Repbase Rel. 10.01) using Bismarck (v0.22.1) ^147^ allowing soft-clipping. Only pairs for which read #2 mapped to the L1HS consensus in the correct orientation were subsequently analyzed (Samtools v1.3) ^152^. The selected pairs were mapped against hg38 reference human genome using Bismarck in end-to-end mode using the following options: `--minins 250 --maxins 1250 --score_min L,-0.6,-0.6`. At this stage, mapped read pairs support L1 elements included in the reference genome (reference L1s). To identify non-reference L1 insertions, we extracted reads #1 from unmapped pairs using seqtk (v1.3, https://github.com/lh3/seqtk) and remapped them alone against hg38 with Bismarck in local mode. This read rescue procedure allowed us to identify: (i) discordant pairs when read #1 mapped end-to-end to hg38; and (ii) split read if the 5’ end of read #1 mapped partially to hg38 but its 3’ end mapped to L1HS consensus sequence. We filtered out discordant pairs with read #2 showing more than 4.5% divergence towards L1HS consensus sequence as they correspond to artefactual chimeras formed with old L1 elements. Finally, properly mapped pairs and read #1 singletons were pooled in a single .bam file, and deduplicated with Picard tools (v1.136, https://broadinstitute.github.io/picard/). As a conservative assumption, we considered read pairs as redundant if their read #1 starts at the same genomic position, since this situation reflects an identical random break site obtained upon sonication.

#### L1 calling

We identified reference L1 by intersecting properly mapped pairs with annotated L1 elements in UCSC repeatmasker track ^161^ using BEDtools ^151^. A minimum of 10 non-redundant reads was required to call a reference L1 element. The coordinates of the elements were extracted from UCSC repeatmasker track. To identify non-reference L1 elements, we clustered reads #1 of discordant pairs and split reads less than 100 bp apart, excluding those intersecting with previously identified reference L1s, using BEDtools. A minimum of 10 non-redundant reads, including at least 2 split reads, was required to call a non-reference L1 element. We used the break point of split reads to precisely define the insertion sites at nucleotide resolution (a 2-bp interval spanning the integration point with 0-based coordinates). Finally, candidate L1 elements not in assembled chromosomes (chr1 to chr22, chrX or chrY) or falling in ENCODE Unified GRCh38 Blacklist regions (ENCFF356LFX) were filtered out with BEDtools.

#### L1 CpG methylation calling

We called CpG methylation in individual read pair for each reference and non-reference L1 locus, including any covered upstream L1 flanking sequence, using the `bismark_methylation_extractor` script from Bismarck. CpG methylation patterns for individual loci were summarized and visualized using MethPat ^153^

#### Assessment of bs-ATLAS-seq recovery rate

To estimate the fraction of elements detected by bs-ATLAS-seq in each L1 family, we compiled a list of reference L1 elements using hg38 UCSC repeatmasker track filtered to keep only the assembled chromosomes (chr1 to chr22, chrX and chrY) and to remove elements in ENCODE Unified GRCh38 Blacklist regions. The recovery rate was calculated for each sample, taking into account the sex of the donor (presence or absence of a Y chromosome). Given the ongoing activity of L1HS elements in modern humans, and the fact that the reference human genome is a composite assembly obtained from a small number of individuals, it is expected that a given sample only contains a fraction of reference L1HS elements. Thus, the calculated rate is a lower estimate. Reference L1 elements were considered as full length if their length is >5900 bp. Non-reference L1 were assumed to be full-length. L1HS subfamilies were deduced from diagnostic SNPs in the reference sequence ^162^.

#### Assessment of bs-ATLAS-seq false positive rate

To estimate the percentage of false positive L1 detected by bs-ATLAS-seq, we compared candidate L1 elements with databases of known non-reference insertions (KNR), such as euL1db ^29^, the 1000 genome project (1KGP) ^23^ or previous mapping of L1HS in the same cell lines using 3’ junction amplification and Ion Torrent-based single-end sequencing (3’-ATLAS-seq) ^9^. Only 3 candidate non-reference insertions appeared unknown (**Table S3**). chr18:15193133-15193135:-:L1HS:NONREF was validated by nanopore sequencing (**Table S4**). The two others, chr7:140709367-140709369:-:L1HS:NONREF and chr10:38190899-38190901:+:L1HS:NONREF, were validated by PCR of their junctions with the flanking sequence (**Table S1** and **Figure S1G**).

### Cas9-targeted nanopore sequencing

To sequence polymorphic L1 loci, we applied Cas9-targeted nanopore sequencing as described in ^89^ to 125 loci.

#### Extraction of genomic DNA

High molecular weight genomic DNA was extracted from freshly pelleted cells using the Monarch Genomic DNA Purification kit (New England Biolabs). Immediately after extraction, DNA was quantified by fluorometry using a Qubit fluorometer and the dsDNA HS Assay kit (Thermo Fisher Scientific). Fragment length (>10kb) was verified by resolving 100 ng of DNA on a 0.8% agarose gel. DNA was stored at 4°C until library preparation, usually the following day.

#### Design and synthesis of single guide RNAs (sgRNAs)

We designed one sgRNA for each of the 124 potentially polymorphic L1HS loci (*i.e.* empty in at least 50% of the cell lines of the panel as determined by bs-ATLAS-seq). Using precomputed SpCas9 sgRNA target prediction and scoring by the CRISPOR tool ^163^ available in the ‘CRISPR Targets’ track of the UCSC Genome Browser, we selected sgRNAs in the region 900 to 1,500 bp downstream of the targeted L1s, and with the highest scores (at least 55 for the MIT specificity score ^164^ and 35 for Moreno-Mateos (MM) efficiency score ^165^). A control sgRNA (LOU3161) targeting a unique site on chromosome 9 was included as a positive control. The 125 sgRNAs were synthesized as a pool using the EnGen sgRNA Synthesis kit (New England Biolabs), and purified with the Monarch RNA Cleanup kit (New England Biolabs). The sgRNA pool was quantified with the Qubit RNA Assay kit (Thermo Fisher Scientific), aliquoted and stored at -80°C.

#### Library preparation

Cas9 ribonucleoprotein particles (RNPs) were assembled by mixing 60 µmol of the sgRNA pool and Alt-R *S. pyogenes* HiFi Cas9 nuclease V3 (IDT) in equimolar amounts in 30 µL of 1X CutSmart Buffer (New England Biolabs) to reach a final concentration of 2 µM. After a 30min incubation step at 25°C, RNPs were kept on ice. For each cell line, 5 µg of genomic DNA was dephosphorylated by 3 µL of Quick Calf Intestinal Phosphatase (CIP, New England Biolabs) in a total volume of 30 µL for 10 min at 37°C. Then CIP was inactivated by heating the reaction at 80°C for 2 min. Cas9-mediated cut and A-tailing was achieved by adding 10 µL of the Cas9 RNP pool, 1 µL of 10 mM dATP (Thermo Fisher Scientific) and 1 µL of Taq Polymerase (5 U/µL, New England Biolabs) to the CIP reaction. Reactions were incubated at 37°C for 1 h, at 72°C for 5 min, and then kept at 4°C. As a quality control, we performed qPCR using 1 µL of the reaction saved before and after the incubation step to quantify the relative copy number of the intact *RASEF* locus (the target of sgRNA LOU3161) using a pair of primers flanking the cut site (LOU3322: TCACAGGTTGCACACTGGAA, and LOU3323: AGCTCAGCCACTTTTCAGCT) and a pair of primers in *Sox2* as loading control (LOU0695: CATGGGTTCGGTGGTCAAGT, and LOU0696: TGCTGATCATGTCCCGGAGGT). Cleavage was considered as successful if the number of intact target sites decreased by ∼10 to 15-fold. Then, sequencing adapters were ligated to the digested products using the Ligation Sequencing kit (SQK-LSK-110, Oxford Nanopore Technologies) in reactions containing 40 µL of sample, 20 µL of Ligation buffer LNB, 10 µL of NEBNext Quick T4 DNA Ligase (New England Biolabs), 5 µL of Adapter mix AMX-F and 3 µL of nuclease-free water. After 10 min incubation at 20°C, DNA was cleaned up using AMpure XP beads (Beckman Coulter) with a beads-to-sample ratio of 0.3:1 (v/v), washed with the long-fragment buffer (LFB) to retain fragments ≥ 3kb, and eluted from the beads in 13 µL of elution buffer (EB) at room temperature for 30 min to further enrich for fragments longer than 30kb. The purified eluate (∼12 µL with 40-45 fmol of DNA) was ready for sequencing on a MinION flow cell and was kept at 4°C until loading.

#### Sequencing of DNA library

A MinION flow cell (R9.4.1, Oxford Nanopore Technologies) was loaded on a Mk1B sequencer and primed with a mix of Flush buffer FB and Flush Tether FLT. Then 75 µL of the DNA library (12 µL of eluate, 37.5 µL of Sequencing buffer SBII and 25.5 µL of Loading beads LBII) were loaded into the flow cell and sequenced for 72 h with the MinKNOW interface (v20.10.6). Base-calling was performed during the sequencing run using Guppy (v4.2.3).

#### Bioinformatic analysis

To map reads obtained by the protocol described above, we prepared a custom genome including the two possible alleles (empty or filled) for each target locus. Both alleles contained 50 kb upstream and 1 kb downstream of L1, extracted from the human reference genome hg38. L1 insertion sites were deduced from bs-ATLAS-seq experiments. If the targeted L1 is present in the reference genome, the empty allele was made by removing the L1 sequence with bedtools maskfasta (v2.3). If the targeted L1 was absent from the reference genome, the filled allele was built using *ref*orm (https://github.com/gencorefacility/reform) by introducing an L1 consensus sequence at the insertion point. Thus, the custom genome comprises 250 sequences concatenated in a multifasta file. After indexation, nanopore reads were mapped to the custom genome with minimap2 (v20.2) using the following options: `-a -x map-ont` ^149^. Reads with a mapping quality score (MAPQ) of minimum 20 were sorted and filtered using samtools (`samtools view -b -q 20`). As reads partially spanning an L1 element without reaching the upstream flank tend to be soft-clipped and to be wrongly mapped, we kept only reads longer than 7 kb. Zygosity was evaluated by calculating the coverage of each allele with bedtools coverage (v2.3). If a single allele (filled or empty) was covered, the locus was considered as homozygous. Inversely, if both alleles were covered, the locus was considered as heterozygous. For each covered allele, methylation calling was performed with nanopolish (v0.13.2) ^150^. We considered only CpG covered by at least 5 reads. Alignments and methylation were visualized with IGV genome browser (v2.12.3) ^154^.

### Methylated DNA Immunoprecipitation (MeDIP)

MeDIP was performed using the Auto MeDIP Kit on an automated platform SX-8G IP–Star Compact (Diagenode). Briefly, 2.5 μg of DNA was sheared using a Bioruptor Pico to approximately 500-bp fragments, as assessed with D5000 ScreenTape (Agilent). Cycle conditions were as follows: 15 s ON / 90 s OFF, repeated 6 times. A portion of sheared DNA (10 %) was kept as input and the rest of the sheared DNA was immunoprecipitated with α-5-methylcytosine antibody (Diagenode), bound to magnetic beads, and was isolated. qPCR for selected genomic loci was performed and efficiency was calculated as % (me-DNA-IP/total input). Primer sequences are listed in **Table S2**.

### LUMA

To assess global CpG methylation, 500 ng of genomic DNA was digested with MspI+EcoRI and HpaII+EcoRI (NEB) in parallel reactions, EcoRI was included as an internal reference. CpG methylation percentage is defined as the HpaII/MspI ratio. Samples were analyzed using PyroMark Q24 Advanced pyrosequencer.

### RNA-seq

#### RNA extraction

Total RNA was purified from the same cell pellet (split in half) as the genomic DNA for bs-ATLAS-seq by two successive cycles of TRI Reagent extraction (Molecular Research Center) and recovered in 50 µL of milli-Q water. Subsequently, 8 µg of total RNA was treated with 2 U of TURBO DNase (Life technologies) for 20 min at 37°C followed by a 5 min incubation step at room temperature with the DNase Inactivation Reagent. After centrifugation at 10,000 x g for 1.5 min, the supernatants containing the RNA samples were transferred to new tube. RNA was quality-controlled and quantified by UV-spectroscopy (NanoDrop 2000), microfluidic electrophoresis (Agilent 2100 Bioanalyzer) and fluorometric Qubit RNA Assay (Life Technologies).

#### Library preparation and sequencing

Directional poly(A)+ RNA-Seq libraries were prepared using 300 ng of DNase-treated RNA using the Poly (A) mRNA Magnetic Isolation Module and NEBNext Ultra II Directional RNA Library Prep kit for Illumina (New England Biolabs) according to manufacturer’s instructions. Samples were multiplexed and sequenced with 2×75 bp pair-end reads on a NextSeq 500 instrument (Illumina).

#### RNA-seq mapping

RNA-seq raw reads were trimmed to remove fragments of sequencing adapters and regions of poor sequencing quality using the sliding-window mode of Trimmomatic (v0.32) ^145^ and parameters recommended for paired-end reads by the Trimmomatic manual. Read quality before and after trimming was then verified using FASTQC (v0.11.2) (http://www.bioinformatics.babraham.ac.uk/projects/fastqc). Trimmed reads were mapped against the human reference genome hg38 (with GENCODE comprehensive release 29), using STAR (v2.7.5c), with the following non-default parameters: `–outFilterMultimapNmax 1000` (1000 alignments allowed per read-pair), `–alignSJoverhangMin 8` (minimum overhang for unannotated junctions).

#### L1 expression measurement

The extent of L1 expression driven by L1 sense promoter was first approximated through the level of readthrough transcription in the downstream flanking sequence ^9, 34^. Because the R2 read is in the same direction as the RNA fragment, only mapped R2 reads with MAPQ≥20 were considered for the following analyses. They were first extracted from the bam file using samtools (`samtools view -b -f 128 -F 4 -q 20`). Then, the number of mapped R2 reads in a 1 kb-window upstream and downstream the L1 element and in the same orientation as L1 were counted using BEDtools (`coverageBed -s`). Annotated exons overlapping with these regions and on the same strand as L1 were masked. Finally, the 5’ signal was subtracted from the 3’ signal to remove potential noise due to pervasive transcription, and the result was normalized by the number of mapped reads to give a value as L1 reads for 1kb per million of mapped reads (RPKM). Negative values (more 5’ signal than 3’ signal) were set to zero. As a cross-validation, the expression levels of individual L1HS copies were also measured with L1EM (v1.1) and recommended parameters ^104^. To measure L1 expression aggregated at the family-level, we used TEtranscripts (v2.2.1) ^142^, combined with DESeq2 (v1.30.1) for differential expression analysis ^157^.

#### Chimeric transcript discovery

Splice junctions are counted during the mapping step and are summarized in the table `SJ.out.tab` from STAR. Each splice junction is characterized by its coordinates and the number of mapped reads which supports the junction. To detect chimeric transcript between an L1 and a neighboring gene, the “start” and the “end” are dissociated and separately analyze with bedtools intersect. Only splice junction for which one extremity mapped into L1 and the other into an exon (GENCODE comprehensive release 29), and supported by at least 2 uniquely mapped reads, are retained. Then uniquely and multi-mapped reads are summed and normalized by the number of mapped reads per million (RPM).

### 5-aza-2’-deoxycytidine treatment

HCT-116 cells were cultured in McCoy medium supplemented with 10% FBS, 100 U/mL penicillin and 100 µg/mL streptomycin. For 5-aza-2’-deoxycytidine (DAC) treatment, cells were plated at a density of 100,000 cells/well in 6-well plates and treated with DAC at a final concentration of 1 µM for a total of 5 days. Fresh medium and drug were added daily for the first 3 days.

### ChIP-seq

#### Chromatin immunoprecipitation (ChIP)

For ChIP of the transcription factor YY1, exponentially-growing 2102Ep cells were washed twice with PBS and fixed at room temperature by addition of disuccinimidyl glutarate to a final concentration of 2mM and incubation for 45 minutes, followed by two washes in PBS and addition of formaldehyde to a final concentration of 1% and incubation for 15 minutes. For ChIP of histone H3K4me3, 2102Ep cells were fixed by addition of formaldehyde to a final concentration of 1% directly to the cell growth medium. Fixation was stopped by addition of glycine to a final concentration of 125 mM. Fixed cells were washed once quickly and twice for 10 min each with ice-cold PBS, and collected by scraping and centrifugation for 10 min at 500 xg at 4 °C. Nuclei were collected by centrifugation for 5 min at 500 xg at 4 °C and resuspended at 5×10^7^ cells/mL in 900 μL of ice-cold L2 buffer (50 mM Tris pH 8.0, 5 mM EDTA, 1% SDS) containing protease inhibitors. Chromatin was fragmented by sonication to an average size of 600–700 bp (typically 9 cycles of 10 s sonication, 1 min recovery on ice, using a micro-tip sonicator) and insoluble debris was pelleted by centrifugation. A 50 μL-aliquot was removed from each sample and analyzed by agarose gel electrophoresis after DNA extraction to verify fragmentation. Fragmented chromatin was diluted with 9 volumes of buffer DB (50 mM Tris pH8, 200 mM NaCl, 5 mM EDTA, 0.5% NP40), and 1 μg of anti-YY1 antibody (C15410345, Diagenode) or anti-H3K4me3 antibody (ab8580, Abcam) was added to each 1 mL of chromatin and incubated overnight at 4 °C with rotation. Antibody-bound chromatin was pulled-down by addition of 25 μL of protein-A Dynabeads (Invitrogen) for anti-YY1 ChIP, or by addition of 15 μL of protein-A sepharose for anti-H3K4me3 ChIP, incubated for 30 min at 4°C, and collected using a magnet for dynabead-bound chromatin, or by centrifugation. Chromatin-bound beads were washed once quickly and 4 times for 5 min each with 900 μL of ice-cold buffer WB (20 mM Tris pH 8.0, 500 mM NaCl, 2 mM EDTA, 1% NP40, 0.1% SDS), followed by washing for 5 min each with 900 μL of ice-cold TE followed by 350 μL of ice-cold 10 mM Tris pH 8.0.

#### Library preparation and sequencing

For YY1 ChIP, immunoprecipitated chromatin was tagmented on beads based on the Diagenode ‘TAG kit for chipmentation’ protocol, using a total tagmentation time of 15 minutes, and sequencing libraries were prepared from tagmented samples by PCR amplification using Kapa HiFi polymerase (Roche). For H3K4me3 ChIP, immunoprecipitated chromatin was released by incubating beads three times in buffer EB (TE + 2% SDS) for 5 min at room temperature with periodic tickling, and pooling the supernatants after collection; fixation was then reverted by overnight incubation at 65 °C, and DNA was directly purified using the MinElute PCR purification kit (Qiagen), eluted in 30 μL elution buffer, and sequencing libraries were prepared using the NEBNext Ultra II DNA library kit (New England Biolabs). Samples were sequenced using a paired-end strategy on a NextSeq500 instrument (Illumina).

#### ChIP-seq analysis

Sequencing reads were trimmed with cutadapt (-q 10) and were aligned to the human reference genome hg38 using bowtie2 (v2.4.1) with options `--very-sensitive` and `-- end-to-end`. Peaks were called with MACS2 (v2.2.7.1) and the following parameter: `-g 2.9e9` using input DNA as background. For H3K4me3, the broad peak option`(--broad`) was also selected. Coverage was calculated using deeptools (`bamCoverage --minMappingQuality 10 -- normalizeUsing RPKM --binSize 10`), and visualized in IGV.

### Transcription factor enrichment at unmethylated L1

Differential transcription factor binding between unmethylated (mCG<25%) and methylated (mCG>75%) L1HS and L1PA2 subsets was analyzed for each cell line using the Unibind enrichment command line tool (UniBind_enrich.sh, available at https://bitbucket.org/CBGR/unibind_enrichment) and the entire Unibind database (Hg38_robust_UniBind_LOLA.RDS) using the `twoSets` option ^90^.

### Enrichment of genomic features

To allow a fair comparison of the associations of reactivated and non-reactivated L1HS upon 5-aza treatment with a wide range of genomic features, we used a statistical approach in which we generate a large number of controlled *in silico* randomizations of each dataset, and we express the magnitudes of each association as a z-score, which reflects the number of standard deviations by which the measured similarity of any pair of datasets differs from the similarity expected by chance, as previously performed ^67^.

## QUANTIFICATION AND STATISTICAL ANALYSIS

Statistical tests were performed in R and are explicitly stated in each Figure legend.

## SUPPLEMENTAL INFORMATION

Supplemental information includes 5 figures, 6 tables and 1 dataset, and can be found with this article online.

## SUPPLEMENTAL DATA

**Table S1 – Bs-ATLAS-seq oligonucleotides and sequencing statistics.**

Related to **Figure 1** and **Figure S1.**

**Table S2 – MeDIP results and primers.**

Related to **Figure S1**.

**Table S3 – Genomic coordinates and methylation levels of all L1 insertions recovered by bs-ATLAS-seq across all cell lines.**

All genomic coordinates are related to the hg38 reference genome. Related to **Figure 2, Figure 3, Figure S2**, and **Figure S3.**

**Table S4 – Methylation levels of polymorphic L1HS elements obtained by Cas9-guided nanopore sequencing and sgRNA sequences used in this study.**

All genomic coordinates are related to the hg38 reference genome. Related to **Figure S1, Figure 4** and **Figure S4**.

**Table S5 – Chimeric L1 transcripts associated with ESR1-bound L1 loci in MCF-7.**

All genomic coordinates are related to the hg38 reference genome. Related to **Figure 5** and **Figure S5.**

**Table S6 – L1HS expression levels across all cell lines.**

All genomic coordinates are related to the hg38 reference genome. Related to **Figure 6** and **Figure 7**.

**Data S1 – Single-molecule methylation patterns of full-length and 3’ truncated L1 obtained by bs-ATLAS-seq.**

Related to **Figure 1.** Can be downloaded from https://doi.org/10.5281/zenodo.7097318. Coordinates are related to the hg38 reference genome (REF insertions) or to L1HS consensus sequence (NONREF insertions).

## Notes

https://L1methdb.ircan.org

### Summary of Updates

Summary, Introduction and Discussion were updated to clarify; Bibliography has been updated.

## REFERENCES

1. Fueyo, R., Judd, J., Feschotte, C., and Wysocka, J. (2022). Roles of transposable elements in the regulation of mammalian transcription. Nat. Rev. Mol. Cell Biol. 10.1038/s41580-022-00457-y.

2. Modzelewski, A.J., Gan Chong, J., Wang, T., and He, L. (2022). Mammalian genome innovation through transposon domestication. Nat. Cell Biol. 10.1038/s41556-022-00970-4.

3. Mandal, P.K., and Kazazian, H.H. (2008). SnapShot: Vertebrate transposons. Cell 135, 192–192.e1. 10.1016/j.cell.2008.09.028.

4. Cost, G.J., Feng, Q., Jacquier, A., and Boeke, J.D. (2002). Human L1 element target-primed reverse transcription in vitro. EMBO J. 21, 5899–5910. 10.1093/emboj/cdf592.

5. Feng, Q., Moran, J.V., Kazazian, H.H., and Boeke, J.D. (1996). Human L1 retrotransposon encodes a conserved endonuclease required for retrotransposition. Cell 87, 905–916. 10.1016/s0092-8674(00)81997-2.

6. Kulpa, D.A., and Moran, J.V. (2006). Cis-preferential LINE-1 reverse transcriptase activity in ribonucleoprotein particles. Nat. Struct. Mol. Biol. 13, 655–660. 10.1038/nsmb1107.

7. Monot, C., Kuciak, M., Viollet, S., Mir, A.A., Gabus, C., Darlix, J.-L., and Cristofari, G. (2013). The specificity and flexibility of l1 reverse transcription priming at imperfect T-tracts. PLoS Genet. 9, e1003499. 10.1371/journal.pgen.1003499.

8. Goodier, J.L., Ostertag, E.M., and Kazazian, H.H. (2000). Transduction of 3’-flanking sequences is common in L1 retrotransposition. Hum Mol Genet 9, 653–657. 10.1093/hmg/9.4.653.

9. Philippe, C., Vargas-Landin, D.B., Doucet, A.J., van Essen, D., Vera-Otarola, J., Kuciak, M., Corbin, A., Nigumann, P., and Cristofari, G. (2016). Activation of individual L1 retrotransposon instances is restricted to cell-type dependent permissive loci. eLife 5, 166. 10.7554/elife.13926.

10. Pickeral, O.K., Makałowski, W., Boguski, M.S., and Boeke, J.D. (2000). Frequent human genomic DNA transduction driven by LINE-1 retrotransposition. Genome Res. 10, 411–415. 10.1101/gr.10.4.411.

11. Rangwala, S.H., Zhang, L., and Kazazian, H.H. (2009). Many LINE1 elements contribute to the transcriptome of human somatic cells. Genome Biol. 10, R100. 10.1186/gb-2009-10-9-r100.

12. Tubio, J.M.C., Li, Y., Ju, Y.S., Martincorena, I., Cooke, S.L., Tojo, M., Gundem, G., Pipinikas, C.P., Zamora, J., Raine, K., et al. (2014). Mobile DNA in cancer. Extensive transduction of nonrepetitive DNA mediated by L1 retrotransposition in cancer genomes. Science 345, 1251343–1251343. 10.1126/science.1251343.

13. Criscione, S.W., Theodosakis, N., Micevic, G., Cornish, T.C., Burns, K.H., Neretti, N., and Rodić, N. (2016). Genome-wide characterization of human L1 antisense promoter-driven transcripts. BMC Genomics 17, 463. 10.1186/s12864-016-2800-5.

14. Cruickshanks, H.A., and Tufarelli, C. (2009). Isolation of cancer-specific chimeric transcripts induced by hypomethylation of the LINE-1 antisense promoter. Genomics 94, 397–406. 10.1016/j.ygeno.2009.08.013.

15. Denli, A.M., Narvaiza, I., Kerman, B.E., Pena, M., Benner, C., Marchetto, M.C.N., Diedrich, J.K., Aslanian, A., Ma, J., Moresco, J.J., et al. (2015). Primate-specific ORF0 contributes to retrotransposon-mediated diversity. Cell 163, 583–593. 10.1016/j.cell.2015.09.025.

16. Faulkner, G.J., Kimura, Y., Daub, C.O., Wani, S., Plessy, C., Irvine, K.M., Schroder, K., Cloonan, N., Steptoe, A.L., Lassmann, T., et al. (2009). The regulated retrotransposon transcriptome of mammalian cells. Nat. Genet. 41, 563–571. 10.1038/ng.368.

17. Macia, A., Muñoz-Lopez, M., Cortes, J.L., Hastings, R.K., Morell, S., Lucena-Aguilar, G., Marchal, J.A., Badge, R.M., and Garcia-Perez, J.L. (2011). Epigenetic control of retrotransposon expression in human embryonic stem cells. Mol. Cell. Biol. 31, 300–316. 10.1128/mcb.00561-10.

18. Nigumann, P., Redik, K., Mätlik, K., and Speek, M. (2002). Many human genes are transcribed from the antisense promoter of L1 retrotransposon. Genomics 79, 628–634. 10.1006/geno.2002.6758.

19. Pinson, M.-E., Court, F., Masson, A., Renaud, Y., Fantini, A., Bacoeur-Ouzillou, O., Barriere, M., Pereira, B., Guichet, P.-O., Chautard, E., et al. (2022). L1 chimeric transcripts are expressed in healthy brain and their deregulation in glioma follows that of their host locus. Hum. Mol. Genet., ddac056. 10.1093/hmg/ddac056.

20. Speek, M. (2001). Antisense promoter of human L1 retrotransposon drives transcription of adjacent cellular genes. Mol. Cell. Biol. 21, 1973–1985. 10.1128/mcb.21.6.1973-1985.2001.

21. Khan, H., Smit, A., and Boissinot, S. (2006). Molecular evolution and tempo of amplification of human LINE-1 retrotransposons since the origin of primates. Genome Res. 16, 78–87. 10.1101/gr.4001406.

22. Scott, E.C., Gardner, E.J., Masood, A., Chuang, N.T., Vertino, P.M., and Devine, S.E. (2016). A hot L1 retrotransposon evades somatic repression and initiates human colorectal cancer. Genome Res. 26, 745–755. 10.1101/gr.201814.115.

23. Sudmant, P.H., Rausch, T., Gardner, E.J., Handsaker, R.E., Abyzov, A., Huddleston, J., Zhang, Y., Ye, K., Jun, G., Fritz, M.H.-Y., et al. (2015). An integrated map of structural variation in 2,504 human genomes. Nature 526, 75–81. 10.1038/nature15394.

24. Feusier, J., Watkins, W.S., Thomas, J., Farrell, A., Witherspoon, D.J., Baird, L., Ha, H., Xing, J., and Jorde, L.B. (2019). Pedigree-based estimation of human mobile element retrotransposition rates. Genome Res. 29, 1567–1577. 10.1101/gr.247965.118.

25. Belyeu, J.R., Brand, H., Wang, H., Zhao, X., Pedersen, B.S., Feusier, J., Gupta, M., Nicholas, T.J., Brown, J., Baird, L., et al. (2021). De novo structural mutation rates and gamete-of-origin biases revealed through genome sequencing of 2,396 families. Am. J. Hum. Genet. 108, 597–607. 10.1016/j.ajhg.2021.02.012.

26. Borges-Monroy, R., Chu, C., Dias, C., Choi, J., Lee, S., Gao, Y., Shin, T., Park, P.J., Walsh, C.A., and Lee, E.A. (2021). Whole-genome analysis reveals the contribution of non-coding de novo transposon insertions to autism spectrum disorder. Mob. DNA 12, 28. 10.1186/s13100-021-00256-w.

27. Chuang, N.T., Gardner, E.J., Terry, D.M., Crabtree, J., Mahurkar, A.A., Rivell, G.L., Hong, C.C., Perry, J.A., and Devine, S.E. (2021). Mutagenesis of human genomes by endogenous mobile elements on a population scale. Genome Res. 31, 2225–2235. 10.1101/gr.275323.121.

28. Gardner, E.J., Lam, V.K., Harris, D.N., Chuang, N.T., Scott, E.C., Pittard, W.S., Mills, R.E., 1000 Genomes Project Consortium, and Devine, S.E. (2017). The Mobile Element Locator Tool (MELT): population-scale mobile element discovery and biology. Genome Res 27, 1916–1929. 10.1101/gr.218032.116.

29. Mir, A.A., Philippe, C., and Cristofari, G. (2015). euL1db: the European database of L1HS retrotransposon insertions in humans. Nucleic Acids Res. 43, D43–7. 10.1093/nar/gku1043.

30. Ewing, A.D., and Kazazian, H.H. (2010). High-throughput sequencing reveals extensive variation in human-specific L1 content in individual human genomes. Genome Res. 20, 1262–1270. 10.1101/gr.106419.110.

31. Boissinot, S., and Furano, A.V. (2001). Adaptive evolution in LINE-1 retrotransposons. Mol Biol Evol 18, 2186–2194. 10.1093/oxfordjournals.molbev.a003765.

32. Campitelli, L.F., Yellan, I., Albu, M., Barazandeh, M., Patel, Z.M., Blanchette, M., and Hughes, T.R. (2022). Reconstruction of full-length LINE-1 progenitors from ancestral genomes. Genetics, iyac074. 10.1093/genetics/iyac074.

33. Jacobs, F.M.J., Greenberg, D., Nguyen, N., Haeussler, M., Ewing, A.D., Katzman, S., Paten, B., Salama, S.R., and Haussler, D. (2014). An evolutionary arms race between KRAB zinc-finger genes ZNF91/93 and SVA/L1 retrotransposons. Nature 516, 242–245. 10.1038/nature13760.

34. Lanciano, S., and Cristofari, G. (2020). Measuring and interpreting transposable element expression. Nat. Rev. Genet. 21, 721–736. 10.1038/s41576-020-0251-y.

35. Castro-Diaz, N., Ecco, G., Coluccio, A., Kapopoulou, A., Yazdanpanah, B., Friedli, M., Duc, J., Jang, S.M., Turelli, P., and Trono, D. (2014). Evolutionally dynamic L1 regulation in embryonic stem cells. Genes Dev 28, 1397–1409. 10.1101/gad.241661.114.

36. Imbeault, M., Helleboid, P.-Y., and Trono, D. (2017). KRAB zinc-finger proteins contribute to the evolution of gene regulatory networks. Nature 543, 550–554. 10.1038/nature21683.

37. Alves, G., Tatro, A., and Fanning, T. (1996). Differential methylation of human LINE-1 retrotransposons in malignant cells. Gene 176, 39–44. 10.1016/0378-1119(96)00205-3.

38. Deniz, Ö., Frost, J.M., and Branco, M.R. (2019). Regulation of transposable elements by DNA modifications. Nat. Rev. Genet. 76, 65. 10.1038/s41576-019-0106-6.

39. Jönsson, M.E., Ludvik Brattås, P., Gustafsson, C., Petri, R., Yudovich, D., Pircs, K., Verschuere, S., Madsen, S., Hansson, J., Larsson, J., et al. (2019). Activation of neuronal genes via LINE-1 elements upon global DNA demethylation in human neural progenitors. Nat. Commun. 10, 3182–11. 10.1038/s41467-019-11150-8.

40. Molaro, A., Malik, H.S., and Bourc’his, D. (2020). Dynamic Evolution of De Novo DNA Methyltransferases in Rodent and Primate Genomes. Mol. Biol. Evol. 37, 1882–1892. 10.1093/molbev/msaa044.

41. Muotri, A.R., Marchetto, M.C.N., Coufal, N.G., Oefner, R., Yeo, G., Nakashima, K., and Gage, F.H. (2010). L1 retrotransposition in neurons is modulated by MeCP2. Nature 468, 443–446. 10.1038/nature09544.

42. Thayer, R.E., Singer, M.F., and Fanning, T.G. (1993). Undermethylation of specific LINE-1 sequences in human cells producing a LINE-1-encoded protein. Gene 133, 273–277. 10.1016/0378-1119(93)90651-i.

43. Liu, N., Lee, C.H., Swigut, T., Grow, E., Gu, B., Bassik, M.C., and Wysocka, J. (2018). Selective silencing of euchromatic L1s revealed by genome-wide screens for L1 regulators. Nature 553, 228–232. 10.1038/nature25179.

44. Robbez-Masson, L., Tie, C.H.C., Conde, L., Tunbak, H., Husovsky, C., Tchasovnikarova, I.A., Timms, R.T., Herrero, J., Lehner, P.J., and Rowe, H.M. (2018). The HUSH complex cooperates with TRIM28 to repress young retrotransposons and new genes. Genome Res. 28, 836–845. 10.1101/gr.228171.117.

45. Tunbak, H., Enriquez-Gasca, R., Tie, C.H.C., Gould, P.A., Mlcochova, P., Gupta, R.K., Fernandes, L., Holt, J., van der Veen, A.G., Giampazolias, E., et al. (2020). The HUSH complex is a gatekeeper of type I interferon through epigenetic regulation of LINE-1s. Nat. Commun. 11, 5387. 10.1038/s41467-020-19170-5.

46. Faulkner, G.J., and Billon, V. (2018). L1 retrotransposition in the soma: a field jumping ahead. Mob. DNA 9, 22. 10.1186/s13100-018-0128-1.

47. Burns, K.H. (2017). Transposable elements in cancer. Nat Rev Cancer 17, 415–424. 10.1038/nrc.2017.35.

48. Rodríguez-Martín, B., Alvarez, E.G., Baez-Ortega, A., Zamora, J., Supek, F., Demeulemeester, J., Santamarina, M., Ju, Y.S., Temes, J., Garcia-Souto, D., et al. (2020). Pan-cancer analysis of whole genomes identifies driver rearrangements promoted by LINE-1 retrotransposition. Nat. Genet. 52, 306–319. 10.1038/s41588-019-0562-0.

49. Jang, H.S., Shah, N.M., Du, A.Y., Dailey, Z.Z., Pehrsson, E.C., Godoy, P.M., Zhang, D., Li, D., Xing, X., Kim, S., et al. (2019). Transposable elements drive widespread expression of oncogenes in human cancers. Nat Genet 51, 611–617. 10.1038/s41588-019-0373-3.

50. Weber, B., Kimhi, S., Howard, G., Eden, A., and Lyko, F. (2010). Demethylation of a LINE-1 antisense promoter in the cMet locus impairs Met signalling through induction of illegitimate transcription. Oncogene 29, 5775–5784. 10.1038/onc.2010.227.

51. Wolff, E.M., Byun, H.-M.M., Han, H.F., Sharma, S., Nichols, P.W., Siegmund, K.D., Yang, A.S., Jones, P.A., and Liang, G. (2010). Hypomethylation of a LINE-1 promoter activates an alternate transcript of the MET oncogene in bladders with cancer. PLoS Genet. 6, e1000917. 10.1371/journal.pgen.1000917.

52. Kapusta, A., Kronenberg, Z., Lynch, V.J., Zhuo, X., Ramsay, L., Bourque, G., Yandell, M., and Feschotte, C. (2013). Transposable elements are major contributors to the origin, diversification, and regulation of vertebrate long noncoding RNAs. PLoS Genet. 9, e1003470. 10.1371/journal.pgen.1003470.

53. Hall, L.L., Carone, D.M., Gomez, A.V., Kolpa, H.J., Byron, M., Mehta, N., Fackelmayer, F.O., and Lawrence, J.B. (2014). Stable C0T-1 repeat RNA is abundant and is associated with euchromatic interphase chromosomes. Cell 156, 907–919. 10.1016/j.cell.2014.01.042.

54. Jachowicz, J.W., Bing, X., Pontabry, J., Bošković, A., Rando, O.J., and Torres-Padilla, M.-E. (2017). LINE-1 activation after fertilization regulates global chromatin accessibility in the early mouse embryo. Nat Genet 49, 1502–1510. 10.1038/ng.3945.

55. Percharde, M., Lin, C.-J., Yin, Y., Guan, J., Peixoto, G.A., Bulut-Karslioglu, A., Biechele, S., Huang, B., Shen, X., and Ramalho-Santos, M. (2018). A LINE1-Nucleolin Partnership Regulates Early Development and ESC Identity. Cell 174, 391–405.e19. 10.1016/j.cell.2018.05.043.

56. Baylin, S.B., and Jones, P.A. (2016). Epigenetic Determinants of Cancer. Cold Spring Harb. Perspect. Biol. 8, a019505. 10.1101/cshperspect.a019505.

57. Nguyen, T.H.M., Carreira, P.E., Sánchez-Luque, F.J., Schauer, S.N., Fagg, A.C., Richardson, S.R., Davies, C.M., Jesuadian, J.S., Kempen, M.-J.H.C., Troskie, R.-L., et al. (2018). L1 Retrotransposon Heterogeneity in Ovarian Tumor Cell Evolution. Cell Rep 23, 3730–3740. 10.1016/j.celrep.2018.05.090.

58. Schauer, S.N., Carreira, P.E., Shukla, R., Gerhardt, D.J., Gerdes, P., Sánchez-Luque, F.J., Nicoli, P., Kindlova, M., Ghisletti, S., Santos, A.D., et al. (2018). L1 retrotransposition is a common feature of mammalian hepatocarcinogenesis. Genome Res 28, 639–653. 10.1101/gr.226993.117.

59. Coufal, N.G., Garcia-Perez, J.L., Peng, G.E., Yeo, G.W., Mu, Y., Lovci, M.T., Morell, M., O’Shea, K.S., Moran, J.V., and Gage, F.H. (2009). L1 retrotransposition in human neural progenitor cells. Nature 460, 1127– 1131. 10.1038/nature08248.

60. Klawitter, S., Fuchs, N.V., Upton, K.R., Muñoz-Lopez, M., Shukla, R., Wang, J., Garcia-Cañadas, M., Lopez-Ruiz, C., Gerhardt, D.J., Sebe, A., et al. (2016). Reprogramming triggers endogenous L1 and Alu retrotransposition in human induced pluripotent stem cells. Nat. Commun. 7, 10286. 10.1038/ncomms10286.

61. Macia, A., Widmann, T.J., Heras, S.R., Ayllon, V., Sanchez, L., Benkaddour-Boumzaouad, M., Muñoz-Lopez, M., Rubio, A., Amador-Cubero, S., Blanco-Jimenez, E., et al. (2017). Engineered LINE-1 retrotransposition in nondividing human neurons. Genome Res. 27, 335–348. 10.1101/gr.206805.116.

62. Salvador-Palomeque, C., Sánchez-Luque, F.J., Fortuna, P.R.J., Ewing, A.D., Wolvetang, E.J., Richardson, S.R., and Faulkner, G.J. (2019). Dynamic Methylation of an L1 Transduction Family during Reprogramming and Neurodifferentiation. Mol. Cell. Biol. 39. 10.1128/mcb.00499-18.

63. Sánchez-Luque, F.J., Kempen, M.-J.H.C., Gerdes, P., Vargas-Landin, D.B., Richardson, S.R., Troskie, R.-L., Jesuadian, J.S., Cheetham, S.W., Carreira, P.E., Salvador-Palomeque, C., et al. (2019). LINE-1 Evasion of Epigenetic Repression in Humans. Mol. Cell 75, 590–604.e12. 10.1016/j.molcel.2019.05.024.

64. Wissing, S., Muñoz-Lopez, M., Macia, A., Yang, Z., Montano, M., Collins, W., Garcia-Perez, J.L., Moran, J.V., and Greene, W.C. (2012). Reprogramming somatic cells into iPS cells activates LINE-1 retroelement mobility. Hum Mol Genet 21, 208–218. 10.1093/hmg/ddr455.

65. Deininger, P., Morales, M.E., White, T.B., Baddoo, M., Hedges, D.J., Servant, G., Srivastav, S., Smither, M.E., Concha, M., deHaro, D.L., et al. (2017). A comprehensive approach to expression of L1 loci. Nucleic Acids Res. 45, e31–e31. 10.1093/nar/gkw1067.

66. Flasch, D.A., Macia, A., Sanchez, L., Ljungman, M., Heras, S.R., Garcia-Perez, J.L., Wilson, T.E., and Moran, J.V. (2019). Genome-wide de novo L1 Retrotransposition Connects Endonuclease Activity with Replication. Cell 177, 837–851.e28. 10.1016/j.cell.2019.02.050.

67. Sultana, T., van Essen, D., Siol, O., Bailly-Bechet, M., Philippe, C., Zine El Aabidine, A., Pioger, L., Nigumann, P., Saccani, S., Andrau, J.-C., et al. (2019). The Landscape of L1 Retrotransposons in the Human Genome Is Shaped by Pre-insertion Sequence Biases and Post-insertion Selection. Mol. Cell 74, 555–570.e7. 10.1016/j.molcel.2019.02.036.

68. O’Neill, K., Brocks, D., and Hammell, M.G. (2020). Mobile genomics: tools and techniques for tackling transposons. Philos. Trans. R. Soc. Lond. B. Biol. Sci. 375, 20190345. 10.1098/rstb.2019.0345.

69. Alexandrova, E.A., Olovnikov, I.A., Malakhova, G.V., Zabolotneva, A.A., Suntsova, M.V., Dmitriev, S.E., and Buzdin, A.A. (2012). Sense transcripts originated from an internal part of the human retrotransposon LINE-1 5ʹ UTR. Gene 511, 46–53. 10.1016/j.gene.2012.09.026.

70. Swergold, G.D. (1990). Identification, characterization, and cell specificity of a human LINE-1 promoter. Mol. Cell. Biol. 10, 6718–6729. 10.1128/mcb.10.12.6718.

71. Woodcock, D.M., Lawler, C.B., Linsenmeyer, M.E., Doherty, J.P., and Warren, W.D. (1997). Asymmetric methylation in the hypermethylated CpG promoter region of the human L1 retrotransposon. J. Biol. Chem. 272, 7810–7816. 10.1074/jbc.272.12.7810.

72. Badge, R.M., Alisch, R.S., and Moran, J.V. (2003). ATLAS: a system to selectively identify human-specific L1 insertions. Am. J. Hum. Genet. 72, 823–838. 10.1086/373939.

73. Hata, K., and Sakaki, Y. (1997). Identification of critical CpG sites for repression of L1 transcription by DNA methylation. Gene 189, 227–234. 10.1016/s0378-1119(96)00856-6.

74. Taylor, D., Lowe, R., Philippe, C., Cheng, K.C.L., Grant, O.A., Zabet, N.R., Cristofari, G., and Branco, M.R. (2022). Locus-specific chromatin profiling of evolutionarily young transposable elements. Nucleic Acids Res. 50, e33. 10.1093/nar/gkab1232.

75. Hoffmann, M.J., and Schulz, W.A. (2005). Causes and consequences of DNA hypomethylation in human cancer. Biochem. Cell Biol. 83, 296–321. 10.1139/o05-036.

76. Schulz, W.A., Steinhoff, C., and Florl, A.R. (2006). Methylation of Endogenous Human Retroelements in Health and Disease. In DNA Methylation: Development, Genetic Disease and Cancer Current Topics in Microbiology and Immunology., W. Doerfler and P. Böhm, eds. (Springer Berlin Heidelberg), pp. 211–250. 10.1007/3-540-31181-5_11.

77. Shademan, M., Zare, K., Zahedi, M., Mosannen Mozaffari, H., Bagheri Hosseini, H., Ghaffarzadegan, K., Goshayeshi, L., and Dehghani, H. (2020). Promoter methylation, transcription, and retrotransposition of LINE-1 in colorectal adenomas and adenocarcinomas. Cancer Cell Int. 20, 426. 10.1186/s12935-020-01511-5.

78. Bird, A.P. (1980). DNA methylation and the frequency of CpG in animal DNA. Nucleic Acids Res. 8, 1499– 1504. 10.1093/nar/8.7.1499.

79. Walser, J.-C., Ponger, L., and Furano, A.V. (2008). CpG dinucleotides and the mutation rate of non-CpG DNA. Genome Res. 18, 1403–1414. 10.1101/gr.076455.108.

80. Pehrsson, E.C., Choudhary, M.N.K., Sundaram, V., and Wang, T. (2019). The epigenomic landscape of transposable elements across normal human development and anatomy. Nat. Commun. 10, 5640. 10.1038/s41467-019-13555-x.

81. Baubec, T., Colombo, D.F., Wirbelauer, C., Schmidt, J., Burger, L., Krebs, A.R., Akalin, A., and Schübeler, D. (2015). Genomic profiling of DNA methyltransferases reveals a role for DNMT3B in genic methylation. Nature 520, 243–247. 10.1038/nature14176.

82. Lister, R., Pelizzola, M., Dowen, R.H., Hawkins, R.D., Hon, G., Tonti-Filippini, J., Nery, J.R., Lee, L., Ye, Z., Ngo, Q.-M., et al. (2009). Human DNA methylomes at base resolution show widespread epigenomic differences. Nature 462, 315–322. 10.1038/nature08514.

83. Morselli, M., Pastor, W.A., Montanini, B., Nee, K., Ferrari, R., Fu, K., Bonora, G., Rubbi, L., Clark, A.T., Ottonello, S., et al. (2015). In vivo targeting of de novo DNA methylation by histone modifications in yeast and mouse. eLife 4, e06205. 10.7554/eLife.06205.

84. Neri, F., Rapelli, S., Krepelova, A., Incarnato, D., Parlato, C., Basile, G., Maldotti, M., Anselmi, F., and Oliviero, S. (2017). Intragenic DNA methylation prevents spurious transcription initiation. Nature 543, 72–77. 10.1038/nature21373.

85. Jeziorska, D.M., Murray, R.J.S., De Gobbi, M., Gaentzsch, R., Garrick, D., Ayyub, H., Chen, T., Li, E., Telenius, J., Lynch, M., et al. (2017). DNA methylation of intragenic CpG islands depends on their transcriptional activity during differentiation and disease. Proc. Natl. Acad. Sci. U. S. A. 114, E7526–E7535. 10.1073/pnas.1703087114.

86. Gkountela, S., Zhang, K.X., Shafiq, T.A., Liao, W.-W., Hargan-Calvopiña, J., Chen, P.-Y., and Clark, A.T. (2015). DNA Demethylation Dynamics in the Human Prenatal Germline. Cell 161, 1425–1436. 10.1016/j.cell.2015.05.012.

87. Grandi, F.C., Rosser, J.M., Newkirk, S.J., Yin, J., Jiang, X., Xing, Z., Whitmore, L., Bashir, S., Ivics, Z., Izsvák, Z., et al. (2015). Retrotransposition creates sloping shores: a graded influence of hypomethylated CpG islands on flanking CpG sites. Genome Res. 25, 1135–1146. 10.1101/gr.185132.114.

88. Gilpatrick, T., Lee, I., Graham, J.E., Raimondeau, E., Bowen, R., Heron, A., Downs, B., Sukumar, S., Sedlazeck, F.J., and Timp, W. (2020). Targeted nanopore sequencing with Cas9-guided adapter ligation. Nat Biotechnol. 1712, 43–46. 10.1038/s41587-020-0407-5.

89. Sarkar, A., Lanciano, S., and Cristofari, G. (2023). Targeted Nanopore Resequencing and Methylation Analysis of LINE-1 Retrotransposons. Methods Mol. Biol. 2607, 173–198. 10.1007/978-1-0716-2883-6_10.

90. Puig, R.R., Boddie, P., Khan, A., Castro-Mondragon, J.A., and Mathelier, A. (2021). UniBind: maps of high-confidence direct TF-DNA interactions across nine species. BMC Genomics 22, 482. 10.1186/s12864-021-07760-6.

91. Jiang, J.-C., Rothnagel, J.A., and Upton, K.R. (2021). Widespread Exaptation of L1 Transposons for Transcription Factor Binding in Breast Cancer. Int. J. Mol. Sci. 22, 5625. 10.3390/ijms22115625.

92. Sun, X., Wang, X., Tang, Z., Grivainis, M., Kahler, D., Yun, C., Mita, P., Fenyö, D., and Boeke, J.D. (2018). Transcription factor profiling reveals molecular choreography and key regulators of human retrotransposon expression. Proc. Natl. Acad. Sci. U. S. A. 115, E5526–E5535. 10.1073/pnas.1722565115.

93. Athanikar, J.N., Badge, R.M., and Moran, J.V. (2004). A YY1-binding site is required for accurate human LINE-1 transcription initiation. Nucleic Acids Res 32, 3846–3855. 10.1093/nar/gkh698.

94. Becker, K.G., Swergold, G., Ozato, K., and Thayer, R.E. (1993). Binding of the ubiquitous nuclear transcription factor YY1 to a cis regulatory sequence in the human LINE-1 transposable element. Hum. Mol. Genet. 2, 1697–1702. 10.1093/hmg/2.10.1697.

95. Kurose, K., Hata, K., Hattori, M., and Sakaki, Y. (1995). RNA polymerase III dependence of the human L1 promoter and possible participation of the RNA polymerase II factor YY1 in the RNA polymerase III transcription system. Nucleic Acids Res. 23, 3704–3709. 10.1093/nar/23.18.3704.

96. Minakami, R., Kurose, K., Etoh, K., Furuhata, Y., Hattori, M., and Sakaki, Y. (1992). Identification of an internal cis-element essential for the human L1 transcription and a nuclear factor(s) binding to the element. Nucleic Acids Res. 20, 3139–3145. 10.1093/nar/20.12.3139.

97. Speek, M. (2001). Antisense promoter of human L1 retrotransposon drives transcription of adjacent cellular genes. Mol. Cell. Biol. 21, 1973–1985. 10.1128/mcb.21.6.1973-1985.2001.

98. Shah, N.M., Jang, H.J., Liang, Y., Maeng, J.H., Tzeng, S.-C., Wu, A., Basri, N.L., Qu, X., Fan, C., Li, A., et al. (2023). Pan-cancer analysis identifies tumor-specific antigens derived from transposable elements. Nat. Genet. 55, 631–639. 10.1038/s41588-023-01349-3.

99. Broome, R., Chernukhin, I., Jamieson, S., Kishore, K., Papachristou, E.K., Mao, S.-Q., Tejedo, C.G., Mahtey, A., Theodorou, V., Groen, A.J., et al. (2021). TET2 is a component of the estrogen receptor complex and controls 5mC to 5hmC conversion at estrogen receptor cis-regulatory regions. Cell Rep. 34. 10.1016/j.celrep.2021.108776.

100. Thennavan, A., Beca, F., Xia, Y., Garcia-Recio, S., Allison, K., Collins, L.C., Tse, G.M., Chen, Y.-Y., Schnitt, S.J., Hoadley, K.A., et al. (2021). Molecular analysis of TCGA breast cancer histologic types. Cell Genomics 1, 100067. 10.1016/j.xgen.2021.100067.

101. Rodić, N., Sharma, R., Sharma, R., Zampella, J., Dai, L., Taylor, M.S., Hruban, R.H., Iacobuzio-Donahue, C.A., Maitra, A., Torbenson, M.S., et al. (2014). Long interspersed element-1 protein expression is a hallmark of many human cancers. Am J Pathol 184, 1280–1286. 10.1016/j.ajpath.2014.01.007.

102. Harris, C.R., Normart, R., Yang, Q., Stevenson, E., Haffty, B.G., Ganesan, S., Cordon-Cardo, C., Levine, A.J., and Tang, L.H. (2010). Association of nuclear localization of a long interspersed nuclear element-1 protein in breast tumors with poor prognostic outcomes. Genes Cancer 1, 115–124. 10.1177/1947601909360812.

103. Donaghey, J., Thakurela, S., Charlton, J., Chen, J.S., Smith, Z.D., Gu, H., Pop, R., Clement, K., Stamenova, E.K., Karnik, R., et al. (2018). Genetic determinants and epigenetic effects of pioneer-factor occupancy. Nat. Genet. 50, 250–258. 10.1038/s41588-017-0034-3.

104. McKerrow, W., and Fenyö, D. (2019). L1EM: A tool for accurate locus specific LINE-1 RNA quantification. Bioinformatics 544, 115–1173. 10.1093/bioinformatics/btz724.

105. de Mendoza, A., Nguyen, T.V., Ford, E., Poppe, D., Buckberry, S., Pflueger, J., Grimmer, M.R., Stolzenburg, S., Bogdanovic, O., Oshlack, A., et al. (2022). Large-scale manipulation of promoter DNA methylation reveals context-specific transcriptional responses and stability. Genome Biol. 23, 1–31. 10.1186/s13059-022-02728-5.

106. Jones, P.A., and Taylor, S.M. (1980). Cellular differentiation, cytidine analogs and DNA methylation. Cell 20, 85–93. 10.1016/0092-8674(80)90237-8.

107. Yang, X., Han, H., De Carvalho, D.D., Lay, F.D., Jones, P.A., and Liang, G. (2014). Gene Body Methylation Can Alter Gene Expression and Is a Therapeutic Target in Cancer. Cancer Cell 26, 577–590. 10.1016/j.ccr.2014.07.028.

108. Hoffman, M.M., Ernst, J., Wilder, S.P., Kundaje, A., Harris, R.S., Libbrecht, M., Giardine, B., Ellenbogen, P.M., Bilmes, J.A., Birney, E., et al. (2013). Integrative annotation of chromatin elements from ENCODE data. Nucleic Acids Res. 41, 827–841. 10.1093/nar/gks1284.

109. Ernst, J., and Kellis, M. (2012). ChromHMM: automating chromatin-state discovery and characterization. Nat. Methods 9, 215–216. 10.1038/nmeth.1906.

110. Ewing, A.D., Smits, N., Sánchez-Luque, F.J., Faivre, J., Brennan, P.M., Richardson, S.R., Cheetham, S.W., and Faulkner, G.J. (2020). Nanopore Sequencing Enables Comprehensive Transposable Element Epigenomic Profiling. Mol. Cell 80, 915–928.e5. 10.1016/j.molcel.2020.10.024.

111. McDonald, T.L., Zhou, W., Castro, C.P., Mumm, C., Switzenberg, J.A., Mills, R.E., and Boyle, A.P. (2021). Cas9 targeted enrichment of mobile elements using nanopore sequencing. Nat. Commun. 12, 3586. 10.1038/s41467-021-23918-y.

112. Ardeljan, D., Taylor, M.S., Ting, D.T., and Burns, K.H. (2017). The Human Long Interspersed Element-1 Retrotransposon: An Emerging Biomarker of Neoplasia. Clin. Chem. 63, 816–822. 10.1373/clinchem.2016.257444.

113. Arnaud, P., Goubely, C., Pélissier, T., and Deragon, J.-M. (2000). SINE Retroposons Can Be Used In Vivo as Nucleation Centers for De Novo Methylation. Mol. Cell. Biol. 20, 3434–3441.

114. Turker, M.S. (2002). Gene silencing in mammalian cells and the spread of DNA methylation. Oncogene 21, 5388–5393. 10.1038/sj.onc.1205599.

115. Martin, A., Troadec, C., Boualem, A., Rajab, M., Fernandez, R., Morin, H., Pitrat, M., Dogimont, C., and Bendahmane, A. (2009). A transposon-induced epigenetic change leads to sex determination in melon. Nature 461, 1135–1138. 10.1038/nature08498.

116. Jähner, D., and Jaenisch, R. (1985). Retrovirus-induced de novo methylation of flanking host sequences correlates with gene inactivity. Nature 315, 594–597. 10.1038/315594a0.

117. Rebollo, R., Karimi, M.M., Bilenky, M., Gagnier, L., Miceli-Royer, K., Zhang, Y., Goyal, P., Keane, T.M., Jones, S., Hirst, M., et al. (2011). Retrotransposon-induced heterochromatin spreading in the mouse revealed by insertional polymorphisms. PLoS Genet. 7, e1002301. 10.1371/journal.pgen.1002301.

118. Rebollo, R., Miceli-Royer, K., Zhang, Y., Farivar, S., Gagnier, L., and Mager, D.L. (2012). Epigenetic interplay between mouse endogenous retroviruses and host genes. Genome Biol. 13, R89. 10.1186/gb-2012-13-10-r89.

119. Quadrana, L., Bortolini Silveira, A., Mayhew, G.F., LeBlanc, C., Martienssen, R.A., Jeddeloh, J.A., and Colot, V. (2016). The Arabidopsis thaliana mobilome and its impact at the species level. eLife 5, 6919. 10.7554/elife.15716.

120. Erdmann, R.M., and Picard, C.L. (2020). RNA-directed DNA Methylation. PLOS Genet. 16, e1009034. 10.1371/journal.pgen.1009034.

121. Brind’Amour, J., Kobayashi, H., Richard Albert, J., Shirane, K., Sakashita, A., Kamio, A., Bogutz, A., Koike, T., Karimi, M.M., Lefebvre, L., et al. (2018). LTR retrotransposons transcribed in oocytes drive species-specific and heritable changes in DNA methylation. Nat. Commun. 9, 3331. 10.1038/s41467-018-05841-x.

122. Ferraj, A., Audano, P.A., Balachandran, P., Czechanski, A., Flores, J.I., Radecki, A.A., Mosur, V., Gordon, D.S., Walawalkar, I.A., Eichler, E.E., et al. (2023). Resolution of structural variation in diverse mouse genomes reveals chromatin remodeling due to transposable elements. Cell Genomics 3, 100291. 10.1016/j.xgen.2023.100291.

123. Fukuda, K., and Shinkai, Y. (2020). SETDB1-Mediated Silencing of Retroelements. Viruses 12, 596. 10.3390/v12060596.

124. Karimi, M.M., Goyal, P., Maksakova, I.A., Bilenky, M., Leung, D., Tang, J.X., Shinkai, Y., Mager, D.L., Jones, S., Hirst, M., et al. (2011). DNA methylation and SETDB1/H3K9me3 regulate predominantly distinct sets of genes, retroelements, and chimeric transcripts in mESCs. Cell Stem Cell 8, 676–687. 10.1016/j.stem.2011.04.004.

125. Matsui, T., Leung, D., Miyashita, H., Maksakova, I.A., Miyachi, H., Kimura, H., Tachibana, M., Lorincz, M.C., and Shinkai, Y. (2010). Proviral silencing in embryonic stem cells requires the histone methyltransferase ESET. Nature 464, 927–931. 10.1038/nature08858.

126. He, J., Fu, X., Zhang, M., He, F., Li, W., Abdul, M.Md., Zhou, J., Sun, L., Chang, C., Li, Y., et al. (2019). Transposable elements are regulated by context-specific patterns of chromatin marks in mouse embryonic stem cells. Nat. Commun. 10, 34. 10.1038/s41467-018-08006-y.

127. Ecco, G., Imbeault, M., and Trono, D. (2017). KRAB zinc finger proteins. Dev. Camb. Engl. 144, 2719– 2729. 10.1242/dev.132605.

128. de la Rica, L., Deniz, Ö., Cheng, K.C.L., Todd, C.D., Cruz, C., Houseley, J., and Branco, M.R. (2016). TET-dependent regulation of retrotransposable elements in mouse embryonic stem cells. Genome Biol. 17, 234. 10.1186/s13059-016-1096-8.

129. Freeman, B., White, T., Kaul, T., Stow, E.C., Baddoo, M., Ungerleider, N., Morales, M., Yang, H., Deharo, D., Deininger, P., et al. (2022). Analysis of epigenetic features characteristic of L1 loci expressed in human cells. Nucleic Acids Res. 50, 1888–1907. 10.1093/nar/gkac013.

130. Bonté, P.-E., Arribas, Y.A., Merlotti, A., Carrascal, M., Zhang, J.V., Zueva, E., Binder, Z.A., Alanio, C., Goudot, C., and Amigorena, S. (2022). Single-cell RNA-seq-based proteogenomics identifies glioblastoma-specific transposable elements encoding HLA-I-presented peptides. Cell Rep. 39, 110916. 10.1016/j.celrep.2022.110916.

131. Burbage, M., Rocañín-Arjó, A., Baudon, B., Arribas, Y.A., Merlotti, A., Rookhuizen, D.C., Heurtebise-Chrétien, S., Ye, M., Houy, A., Burgdorf, N., et al. (2023). Epigenetically controlled tumor antigens derived from splice junctions between exons and transposable elements. Sci. Immunol. 8, eabm6360. 10.1126/sciimmunol.abm6360.

132. Merlotti, A., Sadacca, B., Arribas, Y.A., Ngoma, M., Burbage, M., Goudot, C., Houy, A., Rocañín-Arjó, A., Lalanne, A., Seguin-Givelet, A., et al. (2023). Noncanonical splicing junctions between exons and transposable elements represent a source of immunogenic recurrent neo-antigens in patients with lung cancer. Sci. Immunol. 8, eabm6359. 10.1126/sciimmunol.abm6359.

133. Shah, N.M., Jang, H.J., Liang, Y., Maeng, J.H., Tzeng, S.-C., Wu, A., Basri, N.L., Qu, X., Fan, C., Li, A., et al. (2023). Pan-cancer analysis identifies tumor-specific antigens derived from transposable elements. Nat. Genet. 55, 631–639. 10.1038/s41588-023-01349-3.

134. Grundy, E.E., Diab, N., and Chiappinelli, K.B. (2022). Transposable element regulation and expression in cancer. FEBS J. 289, 1160–1179. 10.1111/febs.15722.

135. Gu, Z., Liu, Y., Zhang, Y., Cao, H., Lyu, J., Wang, X., Wylie, A., Newkirk, S.J., Jones, A.E., Lee, M., et al. (2021). Silencing of LINE-1 retrotransposons is a selective dependency of myeloid leukemia. Nat. Genet. 53, 672–682. 10.1038/s41588-021-00829-8.

136. Jones, P.A., Ohtani, H., Chakravarthy, A., and De Carvalho, D.D. (2019). Epigenetic therapy in immune-oncology. Nat. Rev. Cancer 19, 151–161. 10.1038/s41568-019-0109-9.

137. Jung, H., Choi, J.K., and Lee, E.A. (2018). Immune signatures correlate with L1 retrotransposition in gastrointestinal cancers. Genome Res. 28, 1136–1146. 10.1101/gr.231837.117.

138. Kong, Y., Rose, C.M., Cass, A.A., Williams, A.G., Darwish, M., Lianoglou, S., Haverty, P.M., Tong, A.-J., Blanchette, C., Albert, M.L., et al. (2019). Transposable element expression in tumors is associated with immune infiltration and increased antigenicity. Nat. Commun. 10, 5228. 10.1038/s41467-019-13035-2.

139. Altemose, N., Maslan, A., Smith, O.K., Sundararajan, K., Brown, R.R., Mishra, R., Detweiler, A.M., Neff, N., Miga, K.H., Straight, A.F., et al. (2022). DiMeLo-seq: a long-read, single-molecule method for mapping protein–DNA interactions genome wide. Nat. Methods 19, 711–723. 10.1038/s41592-022-01475-6.

140. Cheetham, S.W., Jafrani, Y.M.A., Andersen, S.B., Jansz, N., Kindlova, M., Ewing, A.D., and Faulkner, G.J. (2022). Single-molecule simultaneous profiling of DNA methylation and DNA-protein interactions with Nanopore-DamID. bioRxiv, 2021.08.09.455753. 10.1101/2021.08.09.455753.

141. Battaglia, S., Dong, K., Wu, J., Chen, Z., Najm, F.J., Zhang, Y., Moore, M.M., Hecht, V., Shoresh, N., and Bernstein, B.E. (2022). Long-range phasing of dynamic, tissue-specific and allele-specific regulatory elements. Nat. Genet. 54, 1504–1513. 10.1038/s41588-022-01188-8.

142. Jin, Y., Tam, O.H., Paniagua, E., and Hammell, M. (2015). TEtranscripts: a package for including transposable elements in differential expression analysis of RNA-seq datasets. Bioinformatics 31, 3593–3599. 10.1093/bioinformatics/btv422.

143. Du, Q., Smith, G.C., Luu, P.L., Ferguson, J.M., Armstrong, N.J., Caldon, C.E., Campbell, E.M., Nair, S.S., Zotenko, E., Gould, C.M., et al. (2021). DNA methylation is required to maintain both DNA replication timing precision and 3D genome organization integrity. Cell Rep. 36, 109722. 10.1016/j.celrep.2021.109722.

144. Martin, M. (2011). Cutadapt removes adapter sequences from high-throughput sequencing reads. EMBnet.journal 17, pp-10. 10.14806/ej.17.1.200.

145. Bolger, A.M., Lohse, M., and Usadel, B. (2014). Trimmomatic: a flexible trimmer for Illumina sequence data. Bioinformatics 30, 2114–2120. 10.1093/bioinformatics/btu170.

146. Langmead, B., and Salzberg, S.L. (2012). Fast gapped-read alignment with Bowtie 2. Nat. Methods 9, 357–359. 10.1038/nmeth.1923.

147. Krueger, F., and Andrews, S.R. (2011). Bismark: a flexible aligner and methylation caller for Bisulfite-Seq applications. Bioinformatics 27, 1571–1572. 10.1093/bioinformatics/btr167.

148. Dobin, A., Davis, C.A., Schlesinger, F., Drenkow, J., Zaleski, C., Jha, S., Batut, P., Chaisson, M., and Gingeras, T.R. (2013). STAR: ultrafast universal RNA-seq aligner. Bioinformatics 29, 15–21. 10.1093/bioinformatics/bts635.

149. Li, H. (2018). Minimap2: pairwise alignment for nucleotide sequences. Bioinformatics 34, 3094–3100. 10.1093/bioinformatics/bty191.

150. Loman, N.J., Quick, J., and Simpson, J.T. (2015). A complete bacterial genome assembled de novo using only nanopore sequencing data. Nat. Methods 12, 733–735. 10.1038/nmeth.3444.

151. Quinlan, A.R., and Hall, I.M. (2010). BEDTools: a flexible suite of utilities for comparing genomic features. Bioinformatics 26, 841–842. 10.1093/bioinformatics/btq033.

152. Danecek, P., Bonfield, J.K., Liddle, J., Marshall, J., Ohan, V., Pollard, M.O., Whitwham, A., Keane, T., McCarthy, S.A., Davies, R.M., et al. (2021). Twelve years of SAMtools and BCFtools. GigaScience 10, giab008. 10.1093/gigascience/giab008.

153. Wong, N.C., Pope, B.J., Candiloro, I.L., Korbie, D., Trau, M., Wong, S.Q., Mikeska, T., Zhang, X., Pitman, M., Eggers, S., et al. (2016). MethPat: a tool for the analysis and visualisation of complex methylation patterns obtained by massively parallel sequencing. BMC Bioinformatics 17, 98. 10.1186/s12859-016-0950-8.

154. Thorvaldsdóttir, H., Robinson, J.T., and Mesirov, J.P. (2013). Integrative Genomics Viewer (IGV): high-performance genomics data visualization and exploration. Brief Bioinform 14, 178–192. 10.1093/bib/bbs017.

155. Ramirez, F., Dündar, F., Diehl, S., Grüning, B.A., and Manke, T. (2014). deepTools: a flexible platform for exploring deep-sequencing data. Nucleic Acids Res 42, W187–91. 10.1093/nar/gku365.

156. Zhang, Y., Liu, T., Meyer, C.A., Eeckhoute, J., Johnson, D.S., Bernstein, B.E., Nusbaum, C., Myers, R.M., Brown, M., Li, W., et al. (2008). Model-based analysis of ChIP-Seq (MACS). Genome Biol. 9, R137. 10.1186/gb-2008-9-9-r137.

157. Love, M.I., Huber, W., and Anders, S. (2014). Moderated estimation of fold change and dispersion for RNA-seq data with DESeq2. Genome Biol. 15, 550. 10.1186/s13059-014-0550-8.

158. Josephson, R., Ording, C.J., Liu, Y., Shin, S., Lakshmipathy, U., Toumadje, A., Love, B., Chesnut, J.D., Andrews, P.W., Rao, M.S., et al. (2007). Qualification of embryonal carcinoma 2102Ep as a reference for human embryonic stem cell research. Stem Cells 25, 437–446. 10.1634/stemcells.2006-0236.

159. Mallon, B.S., Hamilton, R.S., Kozhich, O.A., Johnson, K.R., Fann, Y.C., Rao, M.S., and Robey, P.G. (2014). Comparison of the molecular profiles of human embryonic and induced pluripotent stem cells of isogenic origin. Stem Cell Res 12, 376–386. 10.1016/j.scr.2013.11.010.

160. Philippe, C., and Cristofari, G. (2023). Genome-Wide Young L1 Methylation Profiling by bs-ATLAS-seq. Methods Mol. Biol. 2607, 127–150. 10.1007/978-1-0716-2883-6_8.

161. Smit, A.F., Hubley, R., and Green, P. (1996). RepeatMasker Open-3.0.

162. Boissinot, S., Chevret, P., and Furano, A.V. (2000). L1 (LINE-1) retrotransposon evolution and amplification in recent human history. Mol Biol Evol 17, 915–928. 10.1093/oxfordjournals.molbev.a026372.

163. Concordet, J.-P., and Haeussler, M. (2018). CRISPOR: intuitive guide selection for CRISPR/Cas9 genome editing experiments and screens. Nucleic Acids Res. 46, W242–W245. 10.1093/nar/gky354.

164. Doench, J.G., Fusi, N., Sullender, M., Hegde, M., Vaimberg, E.W., Donovan, K.F., Smith, I., Tothova, Z., Wilen, C., Orchard, R., et al. (2016). Optimized sgRNA design to maximize activity and minimize off-target effects of CRISPR-Cas9. Nat. Biotechnol. 34, 184–191. 10.1038/nbt.3437.

165. Moreno-Mateos, M.A., Vejnar, C.E., Beaudoin, J.-D., Fernandez, J.P., Mis, E.K., Khokha, M.K., and Giraldez, A.J. (2015). CRISPRscan: designing highly efficient sgRNAs for CRISPR-Cas9 targeting in vivo. Nat. Methods 12, 982–988. 10.1038/nmeth.3543.

